# A three filament mechanistic model of musculotendon force and impedance

**DOI:** 10.1101/2023.03.27.534347

**Authors:** Matthew Millard, David W. Franklin, Walter Herzog

## Abstract

The force developed by actively lengthened muscle depends on different structures across different scales of lengthening. For small perturbations, the active response of muscle is well captured by a linear-time-invariant (LTI) system: a stiff spring in parallel with a light damper. The force response of muscle to longer stretches is better represented by a compliant spring that can fix its end when activated. Experimental work has shown that the stiffness and damping (impedance) of muscle in response to small perturbations is of fundamental importance to motor learning and mechanical stability, while the huge forces developed during long active stretches are critical for simulating and predicting injury. Outside of motor learning and injury, muscle is actively lengthened as a part of nearly all terrestrial locomotion. Despite the functional importance of impedance and active lengthening, no single muscle model has all of these mechanical properties. In this work, we present the viscoelastic-crossbridge active-titin (VEXAT) model that can replicate the response of muscle to length changes great and small. To evaluate the VEXAT model, we compare its response to biological muscle by simulating experiments that measure the impedance of muscle, and the forces developed during long active stretches. In addition, we have also compared the responses of the VEXAT model to a popular Hill-type muscle model. The VEXAT model more accurately captures the impedance of biological muscle and its responses to long active stretches than a Hill-type model and can still reproduce the force-velocity and force-length relations of muscle. While the comparison between the VEXAT model and biological muscle is favorable, there are some phenomena that can be improved: the low frequency phase response of the model, and a mechanism to support passive force enhancement.

## 1 Introduction

The stiffness and damping of muscle are properties of fundamental importance for motor control, and the accurate simulation of muscle force. The central nervous system (CNS) exploits the activation-dependent stiffness and damping (impedance) of muscle when learning new movements [1], and when moving in unstable [2] or noisy environments [3]. Reaching experiments using haptic manipulanda show that the CNS uses co-contraction to increase the stiffness of the arm when perturbed by an unstable force field [4]. With time and repetition, the force field becomes learned and co-contraction is reduced [1].

The force response of muscle is not uniform, but varies with both the length and time of perturbation. Under constant activation and at a consistent nominal length, Kirsch et al. [5] were able to show that muscle behaves like a linear-time-invariant (LTI) system in response to small^1^ perturbations: a spring-damper of best fit captured over 90% of the observed variation in muscle force for small perturbations (1-3.8% optimal length) over a wide range of bandwidths (4-90Hz). When active muscle is stretched appreciably, titin can develop enormous forces [7], [8], which may prevent further lengthening and injury. The stiffness that best captures the response of muscle to the small perturbations of Kirsch et al. [5] is far greater than the stiffness that best captures the response of muscle to large perturbations [7], [8]. Since everyday movements are often accompanied by both large and small kinematic perturbations, it is important to accurately capture these two processes.

However, there is likely no single muscle model that can replicate the force response of muscle to small [5] and large perturbations [7], [8] while also retaining the capability to reproduce the experiments of Hill [9] and Gordon et al. [10]. Unfortunately, this means that simulation studies that depend on an accurate representation of muscle impedance may reach conclusions well justified in simulation but not in reality. In this work, we focus on formulating a mechanistic muscle model^2^ that can replicate the force response of active muscle to length perturbations both great and small.

There are predominantly three classes of models that are used to simulate musculoskeletal responses: phenomenological models constructed using Hill’s famous force-velocity relationship [9], mechanistic Huxley [11]–[13] models in which individual elastic crossbridges are incorporated, and linearized muscle models [14], [15] which are accurate for small changes in muscle length. Kirsch et al. [5] demonstrated that, for small perturbations, the force response of muscle is well represented by a spring in parallel with a damper. Neither Hill nor Huxley models are likely to replicate Kirsch et al.’s [5] experiments because a Hill muscle model [16], [17] does not contain any active spring elements; while a Huxley model lacks an active damping element. Although linearized muscle models can replicate Kirsch et al.’s experiment [5], these models are only accurate for small changes in length and cannot replicate the Hill’s nonlinear force-velocity relation [9], nor Gordon et al.’s [10] nonlinear force-length relation. However, there have been significant improvements to the canonical forms of phenomenological, mechanistic, and linearized muscle models that warrant closer inspection.

Several novel muscle models have been proposed to improve upon the accuracy of Hill-type muscle models during large active stretches. Forcinito et al. [18] modeled the velocity dependence of muscle using a rheological element^3^ and an elastic rack rather than embedding the force-velocity relationship in equations directly, as is done in a typical Hill model [16], [17]. This modification allows Forcinito et al.’s [18] model to more faithfully replicate the force development of active muscle, as compared to a Hill-type model, during ramp length changes of *≈* 10%^4^ of the optimal CE length, and across velocities of 4 *−* 11% of the maximum contraction velocity^5^. Haeufle et al. [20] made use of a serial-parallel network of spring-dampers to allow their model to reproduce Hill’s force-velocity relationship [9] mechanistically rather than embedding the experimental curve directly in their model. This modification allowed Haeufle et al.’s model to simulate high speed reaching movements that agree more closely with experimental data [21] than is possible with a typical Hill model. Günther et al. [22] evaluated how accurately a variety of spring-damper models were able to reproduce the microscopic increases in crossbridge force in response to small length changes. While each of these models improves upon the force response of the Hill model to ramp length changes, none are likely to reproduce Kirsch et al.’s experiment [5] because the linearized versions of these models lead to a serial, rather than a parallel, connection of a spring and a damper: Kirsch et al. [5] specifically showed (see Figure 3 of [5]) that a serial connection of a spring-damper fails to reproduce the phase shift between force and length present in their experimental data.

Titin [23], [24] has been more recently investigated to explain how lengthened muscle can develop active force when lengthened both within, and beyond, actin-myosin overlap [8]. Titin is a gigantic multi-segmented protein that spans a half-sarcomere, attaching to the Z-line at one end and the middle of the thick filament at the other end [25]. In skeletal muscle, the two sections nearest to the Z-line, the proximal immunoglobulin (IgP) segment and the PEVK segment — rich in the amino acids proline (P), glutamate (E), valine (V) and lysine (K) — are the most compliant [26] since the distal immunoglobulin (IgD) segments bind strongly to the thick filament [27]. Titin has proven to be a complex filament, varying in composition and geometry between different muscle types [28], [29], widely between species [30], and can apply activation dependent forces to actin [31]. It has proven challenging to determine which interactions dominate between the various segments of titin and the other filaments in a sarcomere. Experimental observations have reported titin-actin interactions at myosin-actin binding sites [32], [33], between titin’s PEVK region and actin [34], [35], between titin’s N2A region and actin [36], and between the PEVK-IgD regions of titin and myosin [37]. This large variety of experimental observations has led to a correspondingly large number of proposed hypotheses and models, most of which involve titin interacting with actin [38]–[43], and more recently with myosin [44].

The addition of a titin element to a model will result in more accurate force production during large active length changes, but does not affect the stiffness and damping of muscle at modest sarcomere lengths because of titin’s relatively low stiffness. At sarcomere lengths of 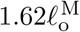 or less, the stiffness of the actin-myosin load path with a single attached crossbridge (0.22 *−* 1.15 pN*/*nm) equals or exceeds the stiffness of 6 passive titin filaments (0.0348 *−* 0.173 pN*/*nm), and our estimated stiffness of 6 active titin filaments (0.0696 *−* 0.346 pN*/*nm, see Appendix A for further details). When fully activated, the stiffness of the actin-myosin load path (4.05 *−* 18.4 pN*/*nm) far exceeds that of both the passive titin (0.0348 *−* 0.173 pN*/*nm), and our estimated active titin (0.0696 *−* 0.346 pN*/*nm) load paths. Since titin-focused models have not made any changes to the modeled myosin-actin interaction beyond a Hill [16], [17] or Huxley [11], [12] model, it is unlikely that these models would be able to replicate Kirsch et al.’s experiments [5].

Although most motor control simulations [2], [45]–[48] make use of the canonical linearized muscle model, phenomenological muscle models have also been used and modified to include stiffness. Sartori et al. [49] modeled muscle stiffness by evaluating the partial derivative of the force developed by a Hill-type muscle model with respect to the contractile element (CE) length. Although this approach is mathematically correct, the resulting stiffness is heavily influenced by the shape of the force-length curve and can lead to inaccurate results: at the optimal CE length this approach would predict an active muscle stiffness of zero since the slope of the force-length curve is zero; on the descending limb this approach would predict a negative active muscle stiffness since the slope of the force-length curve is negative. In contrast, CE stiffness is large and positive near the optimal length [5], and there is no evidence for negative stiffness on the descending limb of the force-length curve [7]. Although the stiffness of the CE can be kept positive by shifting the passive force-length curve, which is at times used in finite-element-models of muscle [50], this introduces a new problem: the resulting passive CE stiffness cannot be lowered to match a more flexible muscle. In contrast, De Groote et al. [51], [52] modeled short-range-stiffness using a stiff spring in parallel with the active force element of a Hill-type muscle model. While the approach of De Groote et al. [51], [52] likely does improve the response of a Hill-type muscle model for small perturbations, there are several drawbacks: the short-range-stiffness of the muscle sharply goes to zero outside of the specified range whereas in reality the stiffness is only reduced [5] (see Fig. 9A); the damping of the canonical Hill-model has been left unchanged and likely differs substantially from biological muscle [5].

In this work, we propose a model that can capture the force development of muscle to perturbations that vary in size and timescale, and yet is described using only a few states making it well suited for large-scale simulations. When active, the response of the model to perturbations within actin-myosin overlap is dominated by a viscoelastic crossbridge element that has different dynamics across time-scales: over brief time-scales the viscoelasticity of the lumped crossbridge dominates the response of the muscle [5], while over longer time-scales the force-velocity [9] and force-length [10] properties of muscle dominate. To capture the active forces developed by muscle beyond actin-myosin overlap we added an active titin element which, similar to existing models [38], [40], features an activation-dependent^6^ interaction between titin and actin. To ensure that the various parts of the model are bounded by reality, we have estimated the physical properties of the viscoelastic crossbridge element as well as the active titin element using data from the literature.

While our main focus is to develop a more accurate muscle model, we would like the model to be well suited to simulating systems that contain tens to hundreds of muscles. Although Huxley models have been used to simulate whole-body movements such as jumping [53], the memory and processing requirements associated with simulating a single muscle with thousands of states is high. Instead of modeling the force development of individual crossbridges, we lump all of the crossbridges in a muscle together so that we have a small number of states to simulate per muscle.

To evaluate the proposed model, we compare simulations of experiments to original data. We examine the response of active muscle to small perturbations over a wide band-width by simulating the stochastic perturbation experiments of Kirsch et al. [5]. Herzog et al.’s [7] active-lengthening experiments are used to evaluate the response of the model when it is actively lengthened within actin-myosin overlap. Next, we use Leonard et al.’s [8] active lengthening experiments to see how the model compares to reality when it is actively lengthened beyond actin-myosin overlap. In addition, we examine how well the model can reproduce the force-velocity experiments of Hill [9] and force-length experiments of Gordon et al. [10]. Since Hill-type models are so commonly used, we also replicate all of the simulated experiments using Millard et al.’s [17] Hill-type muscle model to make the differences between these two types of models clear.

## 2 Model

We begin by treating whole muscle as a scaled half-sarcomere that is pennated at an angle *α* with respect to a tendon (Fig. 1A). The assumption that mechanical properties scale with size is commonly used when modeling muscle [16] and makes it possible to model vastly different musculotendon units (MTUs) by simply changing the architectural and contraction properties: the maximum isometric force 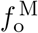, the optimal CE length 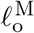 (at which the CE develops 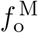), the pennation angle *α*_o_ of the CE (at a length of 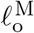) with respect to the tendon, the maximum shortening velocity 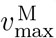 of the CE, and the slack length of the tendon 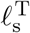. Many properties of sarcomeres scale with 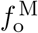 and 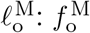 scales with physiological cross-sectional area [54], the force-length property scales with 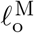 [55], the maximum normalized shortening velocity of different CE types scales with 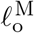 across animals great and small [56], and titin’s passive-force-length properties scale from single molecules to myofibrils [57], [58]

**Figure 1.**
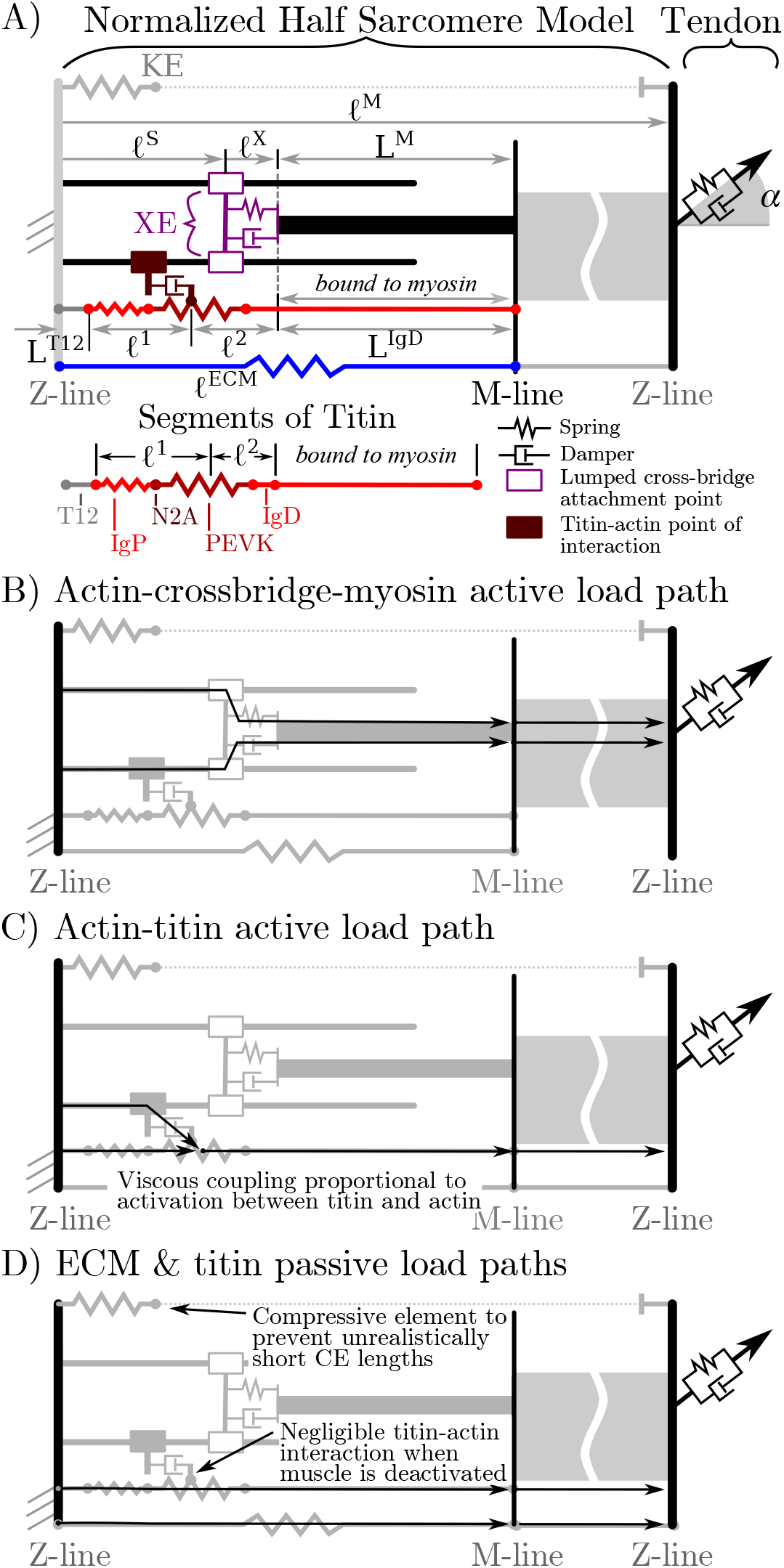
The name of the VEXAT model comes from the viscoelastic crossbridge and active titin elements (A.) in the model. Active tension generated by the lumped crossbridge flows through actin, myosin, and the adjacent sarcomeres to the attached tendon (B.). Titin is modeled as two springs of length *𝓁* ^1^ and *𝓁* ^2^ in series with the rigid segments L^T12^ and L^IgD^. Viscous forces act between titin and actin in proportion to the activation of the muscle (C.), which reduces to negligible values in a purely passive muscle (D.). We modeled actin and myosin as rigid elements; the XE, titin, and the tendon as viscoelastic elements; and the ECM as an elastic element.

The proposed model has several additional properties that we assume scale with 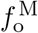 and inversely with 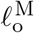 the maximum active isometric stiffness 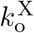 and damping 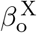, the passive forces due to the extracellular matrix (ECM), and passive forces due to titin. As crossbridge stiffness is well studied [59], we assume that muscle stiffness due to crossbridges scales such that

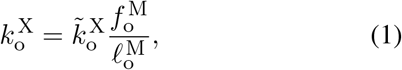

where 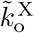 is the maximum normalized stiffness. This scaling is just what would be expected when many crossbridges [59] act in parallel across the cross-sectional area of the muscle, and act in series along the length of the muscle. Although the intrinsic damping properties of crossbridges are not well studied, we assume that the linear increase in damping with activation observed by Kirsch et al. [5] is due to the intrinsic damping properties of individual cross-bridges which will also scale linearly with 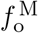 and inversely with 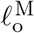

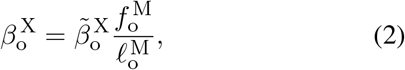

where 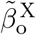 is the maximum normalized damping. For the remainder of the paper, we refer to the proposed model as the VEXAT model due to the viscoelastic (VE) crossbridge (X) and active-titin (AT) elements of the model.

To reduce the number of states needed to simulate the VEXAT model, we lump all of the attached crossbridges into a single lumped crossbridge element (XE) that attaches at *𝓁* ^S^ (Fig. 1A) and has intrinsic stiffness and damping properties that vary with the activation and force-length properties of muscle. The active force developed by the XE at the attachment point to actin is transmitted to the main myosin filament, the M-line, and ultimately to the tendon (Fig. 1B). In addition, since the stiffness of actin [60] and myosin filaments [61] greatly exceeds that of crossbridges [62], we treat actin and myosin filaments as rigid to reduce the number of states needed to simulate this model. Similarly, we have lumped the six titin filaments per half-sarcomere (Fig. 1A) together to further reduce the number of states needed to simulate this model.

The addition of a titin filament to the model introduces an additional active load-path (Fig. 1C) and an additional passive load-path (Fig. 1D). As is typical [16], [17], we assume that the passive elasticity of these structures scale linearly with 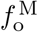 and inversely with 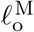. Since the VEXAT model has two passive load paths (Fig. 1D), we further assume that the proportion of the passive force due to the extra-cellular-matrix (ECM) and titin does not follow a scale dependent pattern, but varies from muscle-to-muscle as observed by Prado et al. [58].

As previously mentioned, there are several theories to explain how titin interacts with the other filaments in activated muscle. While there is evidence for titin-actin interaction near titin’s N2A region [36], there is also support for a titin-actin interaction occurring near titin’s PEVK region [34], [35], and for a titin-myosin interaction near the PEVK-IgD region [37]. For the purposes of our model, we will assume a titin-actin interaction because current evidence weighs more heavily towards a titin-actin interaction than a titin-myosin interaction. Next, we assume that the titin-actin interaction takes place somewhere in the PEVK segment for two reasons: first, there is evidence for a titin-actin interaction [34], [35] in the PEVK segment; and second, there is evidence supporting an interaction at the proximal end of the PEVK segment (N2A-actin interaction) [36]. We have left the point within the PEVK segment that attaches to actin as a free variable since there is some uncertainty about what part of the PEVK segment interacts with actin. The nature of the mechanical interaction between titin and the other filaments in an active sarcomere remains uncertain. Here we assume that this interaction is not a rigid attachment, but instead is an activation dependent damping to be consistent with the observations of Kellermayer and Granzier [31] and Dutta et al. [36]: adding titin filaments and calcium slowed, but did not stop, the progression of actin filaments across a plate covered in active crossbridges (an in-vitro motility assay). When activated, we assume that the amount of damping between titin and actin scales linearly with 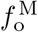 and inversely with 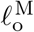

After lumping all of the crossbridges and titin filaments together we are left with a rigid-tendon MTUmodel that has two generalized positions

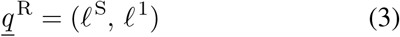

and an elastic-tendon MTUmodel that has three generalized positions

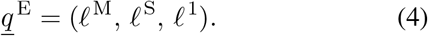

Given these generalized positions, the path length *𝓁* ^P^, and a pennation model, all other lengths in the model can be calculated. Here we use a constant thickness

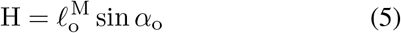

pennation model to evaluate the pennation angle

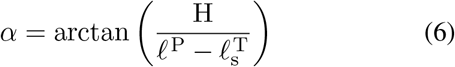

of a CE with a rigid-tendon, and

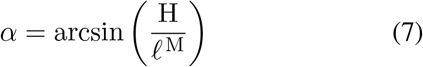

to evaluate the pennation angle of a CE with an elastic-tendon. We have added a small compressive element KE (Fig. 1A) to prevent the model from reaching the numerical singularity that exists as 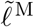 approaches 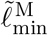, the length at which *α →* 90^*°*^ in Eqns. 6 and 7. The tendon length

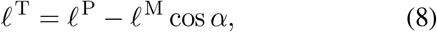

of an elastic-tendon model is the difference between the path length and the CE length along the tendon. The length of the XE

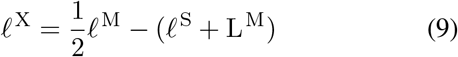

is the difference between the half-sarcomere length and the sum of the average point of attachment *𝓁* ^S^ and the length of the myosin filament L^M^. The length of *𝓁* ^2^, the lumped PEVK-IgD segment, is

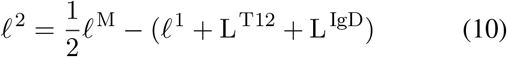

the difference between the half-sarcomere length and the sum of the length from the Z-line to the actin binding site on titin (*𝓁* ^1^) and the length of the IgD segment that is bound to myosin (L^IgD^). Finally, the length of the extra-cellular-matrix *𝓁* ^ECM^ is simply

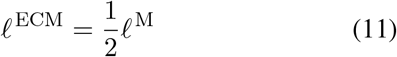

half the length of the CE since we are modeling a half-sarcomere.

We have some freedom to choose the state vector of the model and the differential equations that define how the muscle responds to length and activation changes. The experiments we hope to replicate depend on phenomena that take place at different time-scales: Kirsch et al.’s [5] stochastic perturbations evolve over brief time-scales, while all of the other experiments take place at much longer time-scales. Here we mathematically decouple phenomena that affect brief and long time-scales by making a second-order model that has states of the average point of crossbridge attachment *𝓁* ^S^, and velocity *v* ^S^. When the activation *a* state and the titin-actin interaction model are included, the resulting rigid-tendon model that has a total of four states

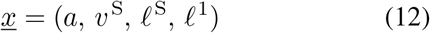

and the elastic-tendon model has

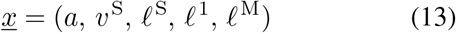

five states. For the purpose of comparison, a Hill-type muscle model with a rigid-tendon has a single state (*a*), while an elastic-tendon model has two states (*a* and *𝓁* ^M^) [17].

Before proceeding, a small note on notation: throughout this work we will use an underbar to indicate a vector, bold font to indicate a curve, a tilde for a normalized quantity, and a capital letter to indicate a constant. Unless indicated otherwise, curves are constructed using 𝒞^2^ continuous^7^ Bézier splines so that the model is compatible with gradient-based optimization. Normalized quantities within the CE follow a specific convention: lengths and velocities are normalized by the optimal CE length 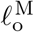, forces by the maximum active isometric tension 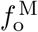, stiffness and damping by 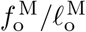. Velocities used as input to the force-velocity relation **f** ^V^ are further normalized by 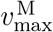 annotated using a hat: 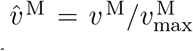. Tendon lengths and velocities are normalized by 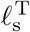 tendon slack length, while forces are normalized by 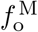.

To evaluate the state derivative of the model, we require equations for 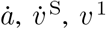 and *v* ^1^, and *v* ^M^ if the tendon is elastic. For 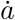 we use of the first order activation dynamics model described in Millard et al. [17]^8^ which uses a lumped first order ordinary-differential-equation (ODE) to describe how a fused tetanus electrical excitation leads to force development in an isometric muscle. We formulated the equation for 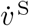 with the intention of having the model behave like a spring-damper over small time-scales, but to converge to the tension developed by a Hill-type model

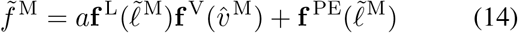

over longer time-scales, where **f** ^L^(·) is the active-force-length curve (Fig. 2A), **f** ^PE^(·) is the passive-force-length curve (Fig. 2A), and **f** ^V^(·) is the force-velocity (Fig. 2B).

**Figure 2.**
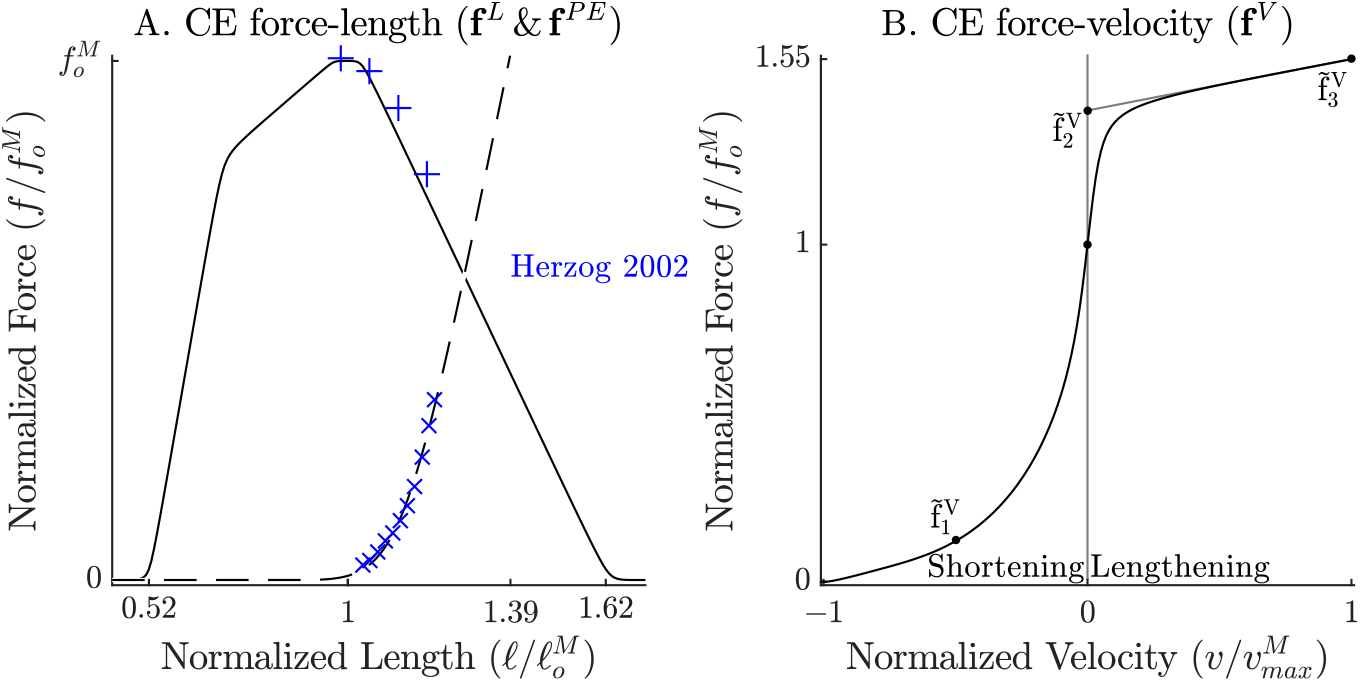
The model relies on Bézier curves to model the nonlinear effects of the active-force-length curve, the passive-force-length curves (A.), and the force-velocity curve (B.). Since nearly all of the reference experiments used in Sec. 3 have used cat soleus, we have fit the active-force-length curve (**f** ^L^(·)) and passive-force-length curves (**f** ^PE^(·)) to the cat soleus data of Herzog and Leonard 2002 [7]. The concentric side of the force-velocity curve (**f** ^V^(·)) has been fitted to the cat soleus data of Herzog and Leonard 1997 [63].

**Figure 3.**
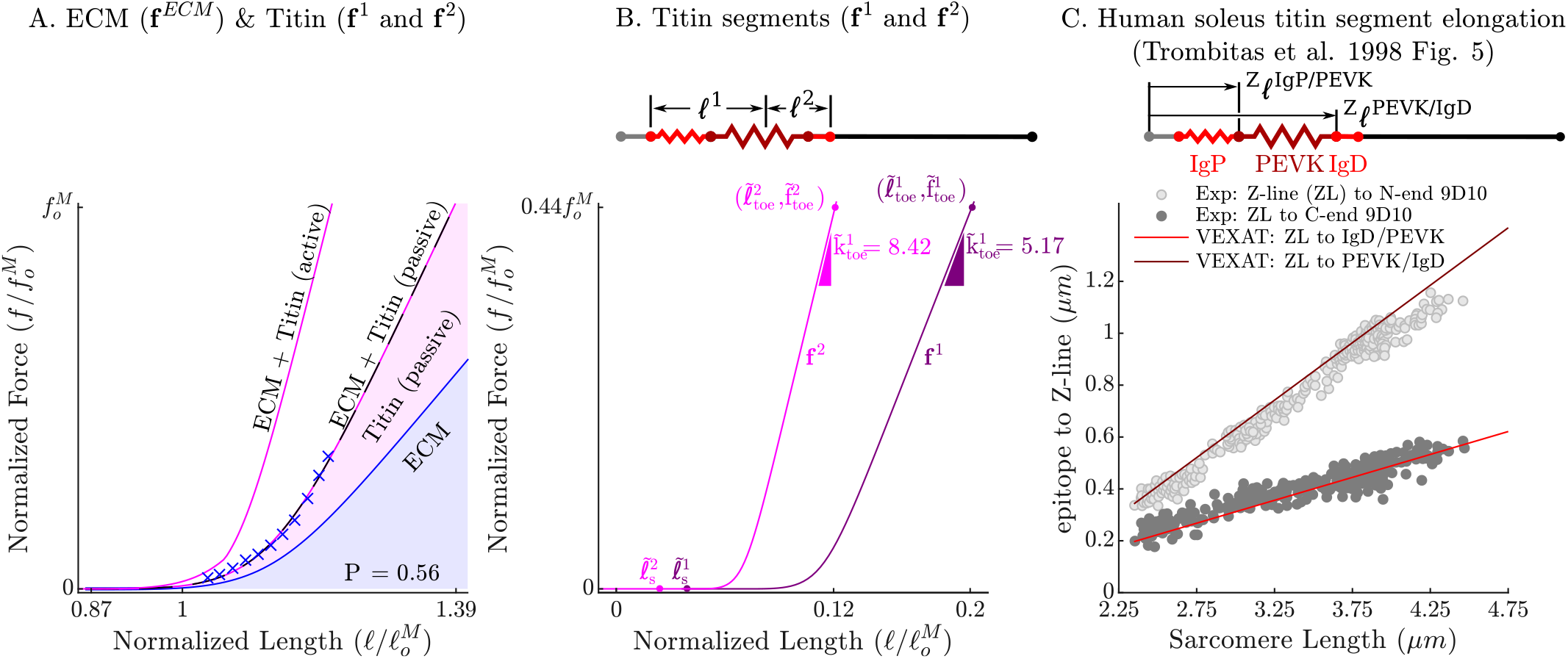
The passive force-length curve has been decomposed such that 56% of it comes from the ECM while 44% comes from titin to match the average of ECM-titin passive force distribution (which ranges from 43%-76%) reported by Prado et al. [58] (A.). The elasticity of the titin segment has been further decomposed into two serially connected sections: the proximal section consisting of the T12, proximal IgP segment and part of the PEVK segment, and the distal section consisting of the remaining PEVK section and the distal Ig segment (B.). The stiffness of the IgP and PEVK segments has been chosen so that the model can reproduce the movements of IgP/PEVK and PEVK/IgD boundaries that Trombitás et al. [26] (C.) observed in their experiments. The curves that appear in subplots A. and B. come from scaling the two-segmented human soleus titin model to cat soleus muscle. The curves that appear in subplot C compare the human soleus titin model’s IgP, PEVK, and IgD force-length relations to the data of Trombitás et al. [26] (see in Appendix B for details).

The normalized tension developed by the VEXAT model

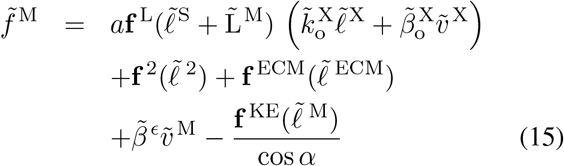

differs from that of a Hill model, Eqn. 14, because it has no explicit dependency on 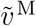, includes four passive terms, and a lumped viscoelastic crossbridge element. The four passive terms come from the ECM element 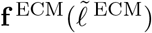 (Fig. 3A), the PEVK-IgD element 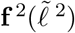 (Fig. 3A and B), the compressive term 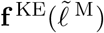 (prevents 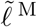 cos *α* from reaching a length of 0), and a numerical damping term 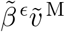 (where 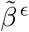 is small). The active force developed by the XE’s lumped crossbridge 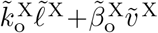 is scaled by the fraction of the XE that is active and attached, 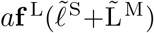, where **f** ^L^(·) is the active-force-length relation (Fig. 2A). We evaluate **f** ^L^ using 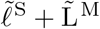 in Eqn. 15, rather than 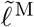 as in Eqn. 14, since actin-myosin overlap is independent of crossbridge strain. With 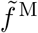 derived, we can proceed to model the acceleration of the CE, 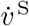, so that it is driven over time by the force imbalance between the XE’s active tension and that of a Hill model.

We set the first term of 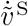 so that, over time, the CE is driven to develop the same active tension as a Hill-type model [17] (terms highlighted in blue)

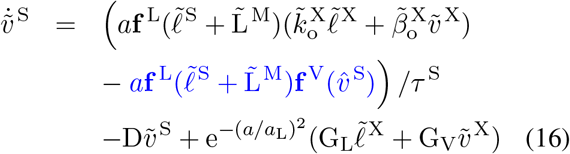

where *τ* ^S^ is a time constant and 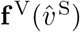 is the force-velocity curve (Fig. 2B). The rate of adaptation of the model’s tension, to the embedded Hill model, is set by the time constant *τ* ^S^: as *τ* ^S^ is decreased the VEXAT model converges more rapidly to a Hill-type model; as *τ* ^S^ is increased the active force produced by the model will look more like a spring-damper. Our preliminary simulations indicate that there is a trade-off to choosing *τ* ^S^: when *τ* ^S^ is large the model will not shorten rapidly enough to replicate Hill’s experiments, while if *τ* ^S^ is small the low-frequency response of the model is compromised when Kirsch et al.’s [5] experiments are simulated.

The remaining two terms, 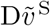 and 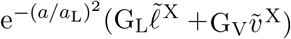, have been included for numerical reasons specific to this model formulation rather than muscle physiology. We include a term that damps the rate of actin-myosin translation, 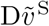, to prevent this second-order system from unrealistically oscillating^9^. The final term 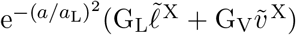, where G_L_ and G_V_ are scalar gains, and *a*_L_ is a low-activation threshold (*a*_L_ is 0.05 in this work). This final term has been included as a consequence of the generalized positions we have chosen. When the CE is nearly deactivated (as *a* approaches *a*_L_), this term forces 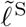 and 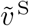 to shadow the location and velocity of the XE attachment point. This ensures that if the XE is suddenly activated, that it attaches with little strain. We had to include this term because we made *𝓁* ^S^ a state variable, rather than *𝓁* ^X^. We chose *𝓁* ^S^ as a state variable, rather than *𝓁* ^X^, so that the states are more equally scaled for numerical integration.

The passive force developed by the CE in Eqn. 15 is the sum of the elastic forces (Fig. 3A) developed by the force-length curves of titin 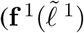 and 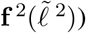 and the 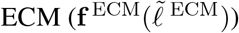. We model titin’s elasticity as being due to two serially connected elastic segments: the first elastic segment 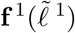 is formed by lumping together the IgP segment and a fraction Q of the PEVK segment, while the second elastic segment 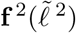 is formed by lumping together the remaining (1 − Q) of the PEVK segment with the free IgD section. Our preliminary simulations of Herzog and Leonard’s active lengthening experiment [7] indicate that a Q value of 0.5, positioning the PEVK-actin attachment point that is near the middle of the PEVK segment, allows the model to develop sufficient tension when actively lengthened. The large section of the IgD segment that is bound to myosin is treated as rigid.

The curves that form 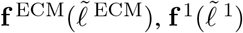, and 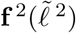 have been carefully constructed to satisfy three experimental observations: that the total passive force-length curve of titin and the ECM match the observed passive force-length curve of the muscle (Fig. 2A and Fig. 3A) [58]; that the proportion of the passive force developed by titin and the ECM is within experimental observations [58] (Fig. 3A); and that the Ig domains and PEVK residues show the same relative elongation as observed by Trombitás et al. [26] (Fig. 3C). Even though Trombitás et al.’s [26] measurements come from human soleus titin, we can construct the force-length curves of other titin isoforms if the number of proximal Ig domains, PEVK residues, and distal Ig domains are known (see Appendix B.3). Since the passive–force-length relation and the results of Trombitás et al. [26] are at modest lengths, we consider two different extensions to the force-length relation to simulate extreme lengths: first, a simple linear extrapolation; second, we extend the force-length relation of each of titin’s segments to follow a worm-like-chain (WLC) model [65] (see Appendix B.3 for details on the WLC model). With titin’s passive force-length relations defined, we can next consider what happens to titin in active muscle.

When active muscle is lengthened, it produces an enhanced force that persists long after the lengthening has ceased called residual force enhancement (RFE) [7]. For the purposes of the VEXAT model, we assume that RFE is produced by titin. Experiments have demonstrated RFE on both the ascending limb [66] and descending limb of the force-length [7] relation. The amount of RFE depends both on the final length of the stretch [67] and the magnitude of the stretch: on the ascending limb the amount of RFE varies with the final length but not with stretch magnitude, while on the descending limb RFE varies with stretch magnitude. To develop RFE, we assume that some point of the PEVK segment bonds with actin through an activation-dependent damper. The VEXAT model’s distal segment of titin, *𝓁* ^2^, can contribute to RFE when the titin-actin bond is formed and CE is lengthened beyond 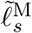, the shortest CE length at which the PEVK-actin bond can form. In this work, we set 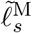 to be equal to the slack length of the CE’s force-length relation 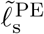 (see Table 1E and H). To incorporate the asymmetric length dependence of RFE [67], we introduce a smooth step-up function

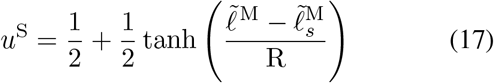

that transitions from zero to one as 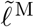 extends beyond 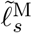, where the sharpness of the transition is controlled by R. Similar to Hisey et al.’s experimental work [67], active lengthening on the ascending limb will produce similar amounts of RFE since 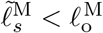 and the titin-actin bond is prevented from forming below 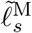. In contrast, the amount of RFE on the descending limb increases with increasing stretch magnitudes since the titin-actin bond is able to form across the entire descending limb.

**Table 1:**
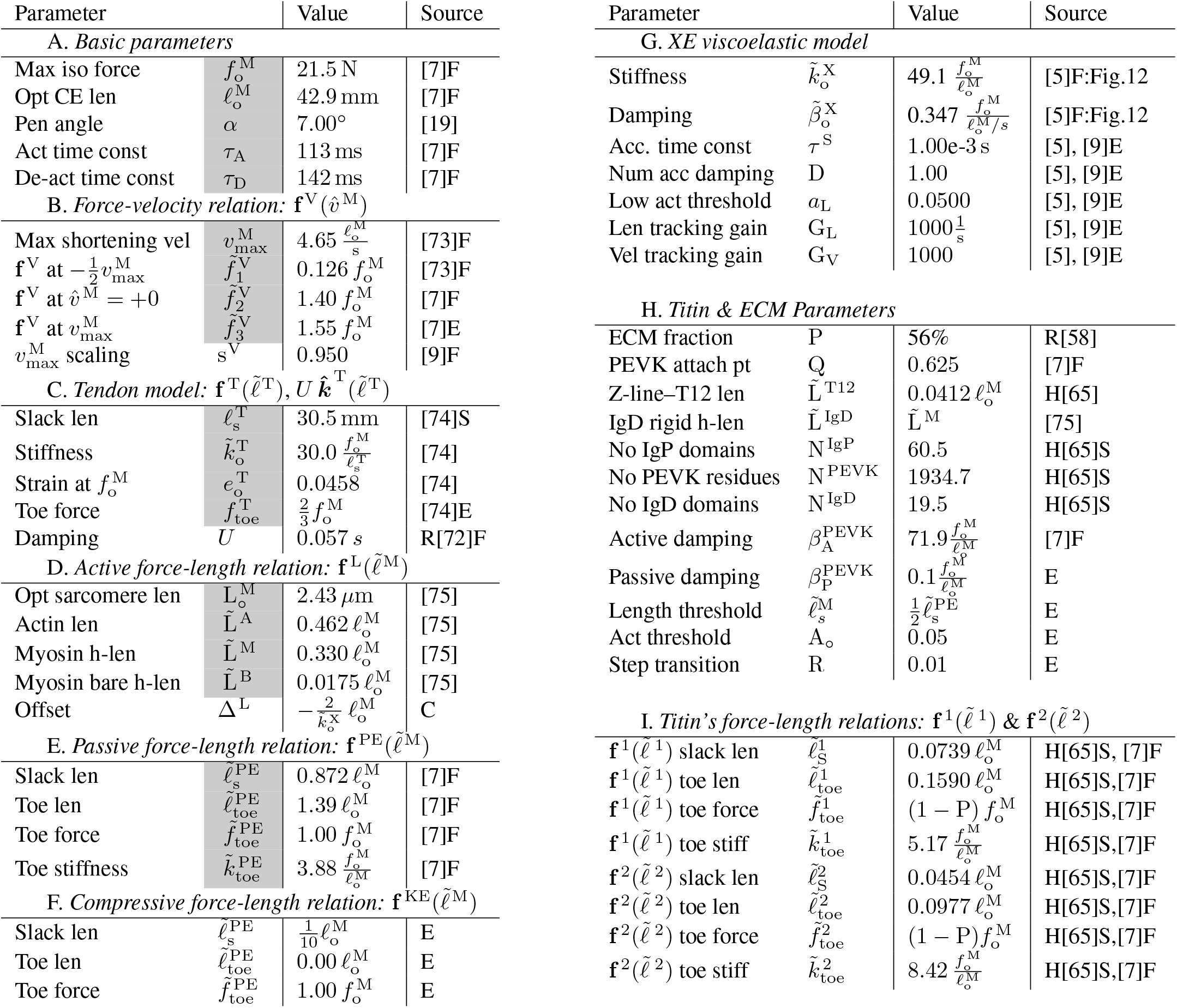
The VEXAT and Hill model’s elastic-tendon cat soleus MTU parameters. The VEXAT model uses all of the Hill model’s parameters which are highlighted in grey. Short forms are used to indicate: length ‘len’, velocity ‘vel’, acceleration ‘acc’, half ‘h’, activation ‘act’, segment ‘seg’, threshold ‘thr’, and stiffness ‘stiff’. The letters ‘R’ or ‘H’ in front of a reference mean the data is from a rabbit or a human, otherwise the data is from cat soleus. The letters following a reference indicate how the data was used to create the parameter: ‘C’ calculated, ‘F’ fit, ‘E’ estimated, or ‘S’ scaled. Most of the VEXAT model’s XE and titin parameters can be used as rough parameter guesses for other muscles because we have expressed these parameters in a normalized space: the values will scale appropriately with changes to 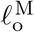 and 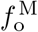. Titin’s force-length curves, however, should be updated if N ^IgP^, N ^PEVK^, or N ^IgD^ differ from the values shown below (see Appendix B.3 for details). Note that the rigid-tendon cat soleus parameters differ from the table below because tendon elasticity affects the fitting of 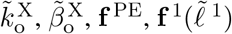, and 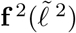). Finally, the parameters related to the compressive element (F.), the XE (G.), and titin (H. and I.) can be used as initial values when simulating the MTU’s other mammals. By making use of these defaults the VEXAT model requires the same number of parameters as a Hill-type muscle model (A.—E.).

At very long CE lengths, the modeled titin-actin bond can literally slip off of the end of the actin filament (Fig. 1A) when the distance between the Z-line and the bond, 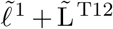, exceeds the length of the actin filament, 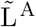. To break the titin-actin bond at long CE lengths we introduce a smooth step-down function

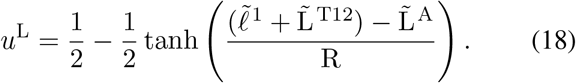

The step-down function *u*^L^ transitions from one to zero when the titin-actin bond 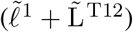 reaches 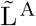, the end of the actin filament.

The strength of the titin-actin bond also appears to vary nonlinearly with activation. Fukutani and Herzog [68] observed that the absolute RFE magnitude produced by actively lengthened fibers is similar between normal and reduced contractile force states. Since these experiments [68] were performed beyond the optimal CE length, titin could be contributing to the observed RFE as previously described. The consistent pattern of absolute RFE values observed by Fukutani and Herzog [68] could be produced if the titin-actin bond saturated at its maximum strength even at a reduced contractile force state. To saturate the titin-actin bond, we use a final smooth step function

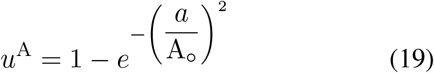

where A_*°*_ is the threshold activation level at which the bond saturates. While we model the strength of the titin-actin bond as being a function of activation, which is proportional Ca^2+^ concentration [69], this is a mathematical convenience. The work of Leonard et al. [8] makes it clear that both Ca^2+^ and crossbridge cycling are needed to allow titin to develop enhanced forces during active lengthening: no enhanced forces are observed in the presence of Ca^2+^ when crossbridge cycling is chemically inhibited. Putting this all together, the active damping acting between the titin and actin filaments is given by the product of 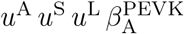 where 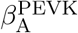 is the maximum damping coefficient.

With a model of the titin-actin bond derived, we can focus on how the bond location moves in response to applied forces. Since we are ignoring the mass of the titin filament, the PEVK-attachment point is balanced by the forces applied to it and the viscous forces developed between titin and actin

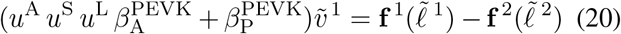

due to the active 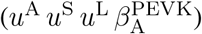 and a small amount of passive damping 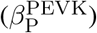 Since Eqn. 20 is linear in 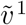, we can solve directly for it

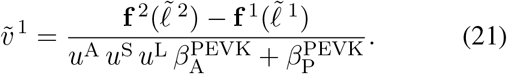

The assumption of whether the tendon is rigid or elastic affects how the state derivative is evaluated and how expensive it is to compute. While all of the position dependent quantities can be evaluated using Eqns. 6-11 and the generalized positions, evaluating the generalized velocities of a rigid-tendon and elastic-tendon model differ substantially.

The CE velocity *v* ^M^ and pennation angular velocity 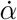 of a rigid-tendon model can be evaluated directly given the path length, velocity, and the time derivatives of Eqns. 6 and 8. After *v* ^1^ is evaluated using Eqn. 21, the velocities of the remaining segments can be evaluated using the time derivatives of Eqns. 9-11.

Evaluating the CE rate of lengthening, *v* ^M^, for an elastic-tendon muscle model is more involved. As is typical of lumped parameter muscle models [16], [17], [70], here we assume that difference in tension, 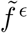, between the CE and the tendon

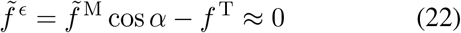

is negligible^10^. During our preliminary simulations it became clear that treating the tendon as an idealized spring degraded the ability of the model to replicate the experiment of Kirsch et al. [5] particularly at high frequencies. Kirsch et al. [5] observed a linear increase in the gain and phase profile between the output force and the input perturbation applied to the muscle. This pattern in gain and phase shift can be accurately reproduced by a spring in parallel with a damper. Due to the way that impedance combines in series ^11^, the models of both the CE and the tendon need to have parallel spring and damper elements so that the entire MTU, when linearized, appears to be a spring in parallel with a damping element. We model tendon force using a nonlinear spring and damper model

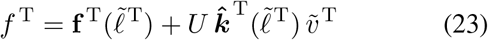

where the damping coefficient 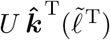, is a linear scal, ing of the normalized tendon stiffness 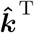 by *U*, a constant scaling coefficient. We have chosen this specific damping model because it fits the data of Netti et al. [72] and captures the structural coupling between tendon stiffness and damping (see Appendix B.1 and Fig. 15 for further details). Now that all of the terms in Eqn. 22 have been explicitly defined, we can use Eqn. 22 to solve for *v* ^M^. Equation 22 becomes linear in *v* ^M^ after substituting the force models described in Eqns. 23 and 15, and the kinematic model equations described in Eqns. 8, 9 and 11 (along with the time derivatives of Eqns. 8-11). After some simplification we arrive at

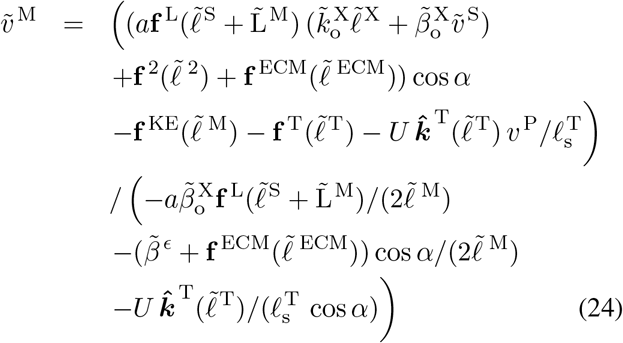

allowing us to evaluate the final state derivative in 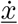. During simulation the denominator of 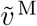 will always be finite since 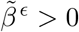, and *α <* 90 ° due to the compressive element. The evaluation of 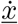 in the VEXAT model is free of numerical singularities, giving it an advantage over a conventional Hill-type muscle model [17]. In addition, the VEXAT’s 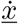 does not require iteration to numerically solve a root, giving it an advantage over a singularity-free formulation of the Hill model [17]. As with previous models, initializing the model’s state is not trivial and required the derivation of a model-specific method (see Appendix C for details).

## 3 Biological Benchmark Simulations

In order to evaluate the model, we have selected three experiments that capture the responses of active muscle to small, medium, and large length changes. The small 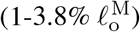 stochastic perturbation experiment of Kirsch et al. [5] demonstrates that the impedance of muscle is well described by a stiff spring in parallel with a damper, and that the spring-damper coefficients vary linearly with active force. The active impedance of muscle is such a fundamental part of motor learning that the amount of impedance, as indicated by co-contraction, is used to define how much learning has actually taken place [1], [77]: co-contraction is high during initial learning, but decreases over time as a task becomes familiar. The active lengthening experiment of Herzog and Leonard [7] shows that modestly stretched 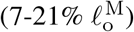 biological muscle has positive stiffness even on the descending limb of the active force-length curve 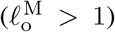. In contrast, a conventional Hill model [16], [17] can have negative stiffness on the descending limb of the active-force-length curve, a property that is both mechanically unstable and unrealistic. The final active lengthening experiment of Leonard et al. [8] unequivocally demonstrates that the CE continues to develop active forces during extreme lengthening 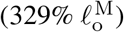 which exceeds actin-myosin overlap. Active force development beyond actin-myosin overlap is made possible by titin, and its activation dependent interaction with actin [8]. The biological benchmark simulations conclude with a replication of the force-velocity experiments of Hill [9] and the force-length experiments of Gordon et al. [10].

The VEXAT model requires the architectural muscle parameters (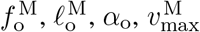, and 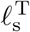) needed by a conventional Hill-type muscle model as well as additional parameters. The additional parameters are needed for these component models: the compressive element (Eqn. 15 and 24), the lumped viscoelastic XE (Eqn. 1 and 2), XE-actin dynamics (Eqn. 16), the two-segment active titin model (Fig. 3), titin-actin dynamics (Eqns. 21), and the tendon damping model (Eqn. 23). Fortunately, there is enough experimental data in the literature that values can be found, fitted, or estimated directly for our simulations of experiments on cat soleus (Table 1), and rabbit psoas fibrils (see Appendix B for fitting details and Appendix H for rabbit psoas fibril model parameters). The parameter values we have established for the cat soleus (Table 1 F.-I.) can serve as initial values when modeling other mammalian MTU’s because these parameters have been normalized (by 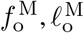, and 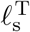 where appropriate) and will scale appropriately given the architectural properties of a different MTU. By making use of these default values, the VEXAT model can be made to represent another MTU using exactly the same number of parameters as a Hill-type muscle model (Table 1A.-E.).

### 3.1 Stochastic Length Perturbation Experiments

In Kirsch et al.’s [5] in-situ experiment, the force response of a cat’s soleus muscle under constant stimulation was measured as its length was changed by small amounts. Kirsch et al. [5] applied stochastic length perturbations (Fig. 4A) to elicit force responses from the muscle (in this case a spring-damper Fig. 4B) across a broad range of frequencies (4-90 Hz) and across a range of small length perturbations 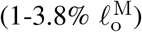. The resulting time-domain signals can be quite complicated (Fig. 4A and B) but contain rich measurements of how muscle transforms changes in length into changes in force.

**Figure 4.**
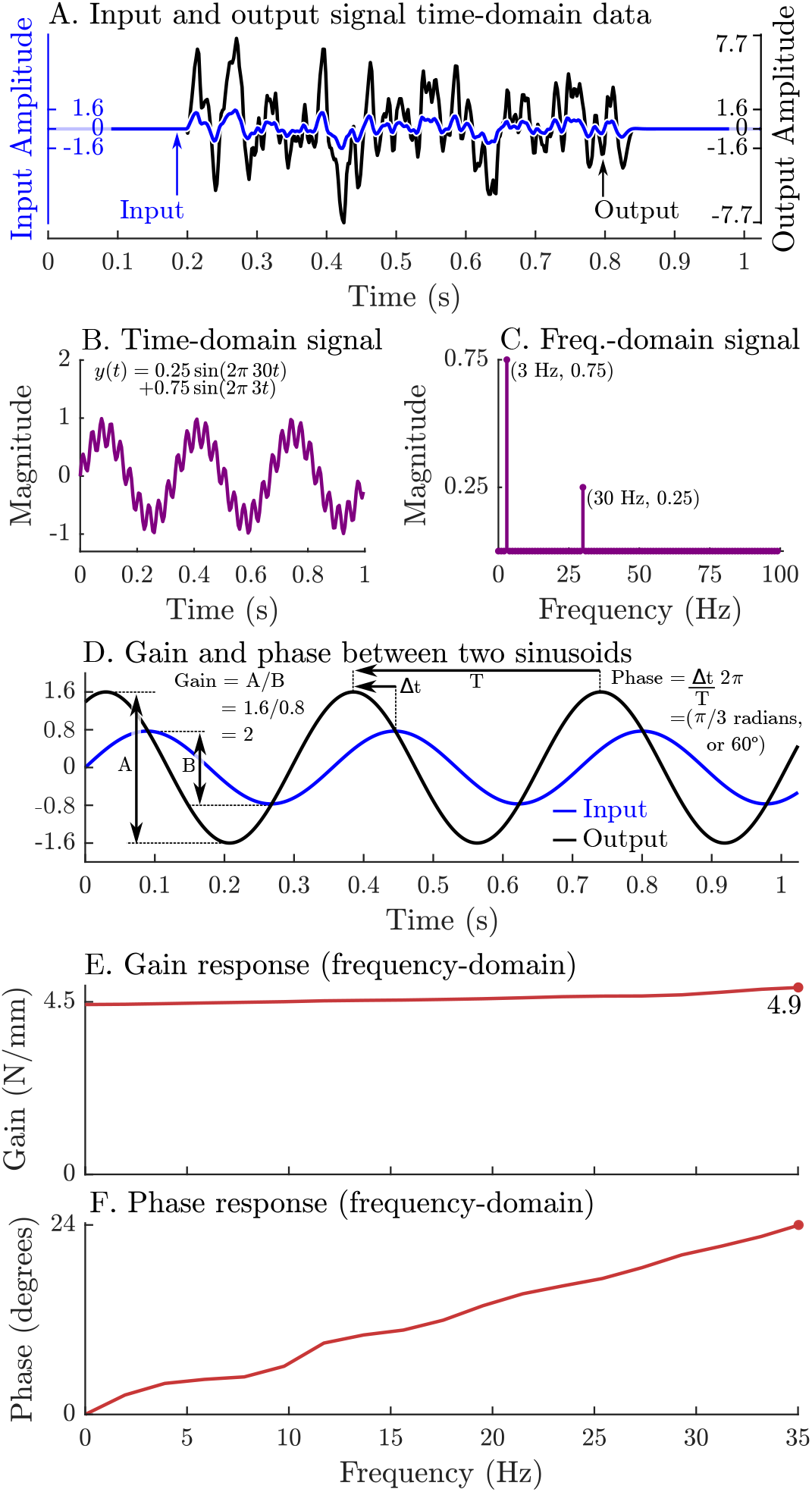
Evaluating a system’s gain and phase response begins by applying a pseudo-random input signal to the system and measuring its output (A). Both the input and output signals (A) are transformed into the frequency domain by expressing these signals as an equivalent sum of scaled and shifted sinusoids (simple example shown in B and C). Each individual input sinusoid is compared with the output sinusoid of the same frequency to evaluate how the system scales and shifts the input to the output (D). This process is repeated across all sinusoid pairs to produce a function that describes how an input sinusoid is scaled (E) and shifted (F) to an output sinusoid using only the measured data (A).

As long as muscle can be considered to be linear (a sinusoidal change in length produces a sinusoidal change in force), then system identification methods [78], [79] can be applied to extract a relationship between length *x*(*t*) and force *y*(*t*). We will give a brief overview of system identification methods here to make methods and results clearer. First, the time-domain signals (*x*(*t*) and *y*(*t*)) are transformed into an equivalent representation in the frequency-domain (*X*(*s*) and *Y* (*s*)) as a sum of scaled and shifted sine curves (Fig. 4B and C) using a Fourier transform [78].

In the frequency domain, we identify an LTI system of best fit *H*(*s*) that describes how muscle transforms changes in length into changes in force such that *Y* (*s*) = *H*(*s*) *X*(*s*). Next, we evaluate how *H*(*s*) scales the magnitude (gain) and shifts the timing (phase) of a sinusoid in *X*(*s*) into a sinusoid of the same frequency in *Y* (*s*) (Fig. 4D). This process is repeated across all frequency-matched pairs of input and output sinusoids to build a function of how muscle scales (Fig. 4E) and shifts (Fig. 4F) input length sinusoids into output force sinusoids. The resulting transformation turns two complicated time-domain signals (Fig. 4A) into a clear relationship in the frequency-domain that describes how muscle transforms length changes into force changes: a very slow (0Hz) length change will result in an output force that is scaled by 4.5 and is in phase (Fig. 4E and F), a 35 Hz sinusoidal length change will produce an output force that is scaled by 4.9 and leads the input signal by 24^*°*^ (Fig. 4E and F), and frequencies between 0 Hz and 35 Hz will be between these two signals in terms of scaling and phase. These patterns of gain and phase can be used to identify a network of spring-dampers that is equivalent to the underlying linear system (the system in Fig. 4 A, E, and F is a 4.46 N/mm spring in parallel with a 0.0089 Ns/mm damper). Since experimental measurements often contain noise and small nonlinearities, the mathematical procedure used to estimate *H*(*s*) and the corresponding gain and phase profiles is more elaborate than we have described (see Appendix D for details).

Kirsch et al. [5] used system identification methods to identify LTI mechanical systems that best describes how muscle transforms input length waveforms to output force waveforms. The resulting LTI system, however, is only accurate when the relationship between input and output is approximately linear. Kirsch et al. [5] used the coherence squared, (*C*_*xy*_)^2^, between the input and output to evaluate the degree of linearity: *Y*(*s*) is a linear transformation of *X*(*s*) at frequencies in which (*C*_*xy*_)^2^ is near one, while *Y* (*s*) cannot be described as a linear function of *X*(*s*) at frequencies in which (*C*_*xy*_)^2^ approaches zero. By calculating (*C*_*xy*_)^2^ between the length perturbation and force waveforms, Kirsch et al. [5] identified the bandwidth in which the muscle’s response is approximately linear. Kirsch et al. [5] set the lower frequency of this band to 4 Hz, and Fig. 3 of Kirsch et al. [5] suggests that this corresponds to (*C*_*xy*_)^2^ *≥* 0.67 though the threshold for (*C*_*xy*_)^2^ is not reported. The upper frequency of this band was set to the cutoff frequency of the low-pass filter applied to the input (15, 35, or 90 Hz). Within this bandwidth, Kirsch et al. [5] compared the response of the specimen to several candidate models and found that a parallel spring-damper fit the muscle’s frequency response best. Next, they evaluated the stiffness and damping coefficients that best fit the muscle’s frequency response [5]. Finally, Kirsch et al. evaluated how much of the muscle’s time-domain response was captured by the spring-damper of best fit by evaluating the variance-accounted-for (VAF) between the two time-domain signals

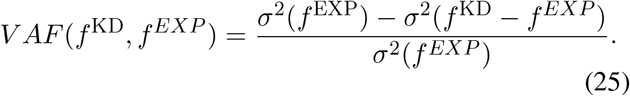

Astonishingly, Kirsch et al. [5] found that a spring-damper of best fit has a VAF of between 78-99%^12^ when compared to the experimentally measured forces *f* ^*EXP*^. By repeating this experiment over a variety of stimulation levels (using both electrical stimulation and the crossed-extension reflex) Kirsch et al. [5] showed that these stiffness and damping coefficients vary linearly with the active force developed by the muscle. Further, Kirsch et al. [5] repeated the experiment using perturbations that had a variety of lengths (0.4 mm, 0.8mm, and 1.6mm) and bandwidths (15Hz, 35Hz, and 90Hz) and observed a peculiar quality of muscle: the damping coefficient of best fit increases as the bandwidth of the perturbation decreases (See Figures 3 and 10 of Kirsch et al. [5] for details). Here we simulate Kirsch et al.’s experiment [5] to determine, first, the VAF of the VEXAT model and the Hill model in comparison to a spring-damper of best fit; second, to compare the gain and phase response of the models to biological muscle; and finally, to see if the spring-damper coefficients of best fit for both models increase with active force in a manner that is similar to the cat soleus that Kirsch et al. studied [5].

To simulate the experiments of Kirsch et al. [5] we begin by creating the 9 stochastic perturbation waveforms used in the experiment that vary in perturbation amplitude (0.4mm, 0.8mm, and 1.6mm) and bandwidth (0-15 Hz, 0-35 Hz, and 0-90 Hz)^13^. The waveform is created using a vector that is composed of random numbers with a range of [*−*1, 1] that begins and ends with a series of zero-valued padding points.

Next, a forward pass of a 2^nd^ order Butterworth filter is applied to the waveform and finally the signal is scaled to the appropriate amplitude (Fig. 5). The muscle model is then activated until it develops a constant tension at a length of 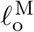. The musculotendon unit is then simulated as the length is varied using the previously constructed waveforms while activation is held constant. To see how impedance varies with active force, we repeated these simulations at ten evenly spaced tensions from 2.5N to 11.5N. Ninety simulations are required to evaluate the nine different perturbation waveforms at each of the ten tension levels. The time-domain length perturbations and force responses of the modeled muscles are used to evaluate the coherence squared of the signal, gain response, and phase responses of the models in the frequency-domain. Since the response of the models might be more nonlinear than biological muscle, we select a bandwidth that meets (*C*_*xy*_)^2^ *>* 0.67 but otherwise follows the bandwidths analyzed by Kirsch et al. [5] (see Appendix D for details).

**Figure 5.**
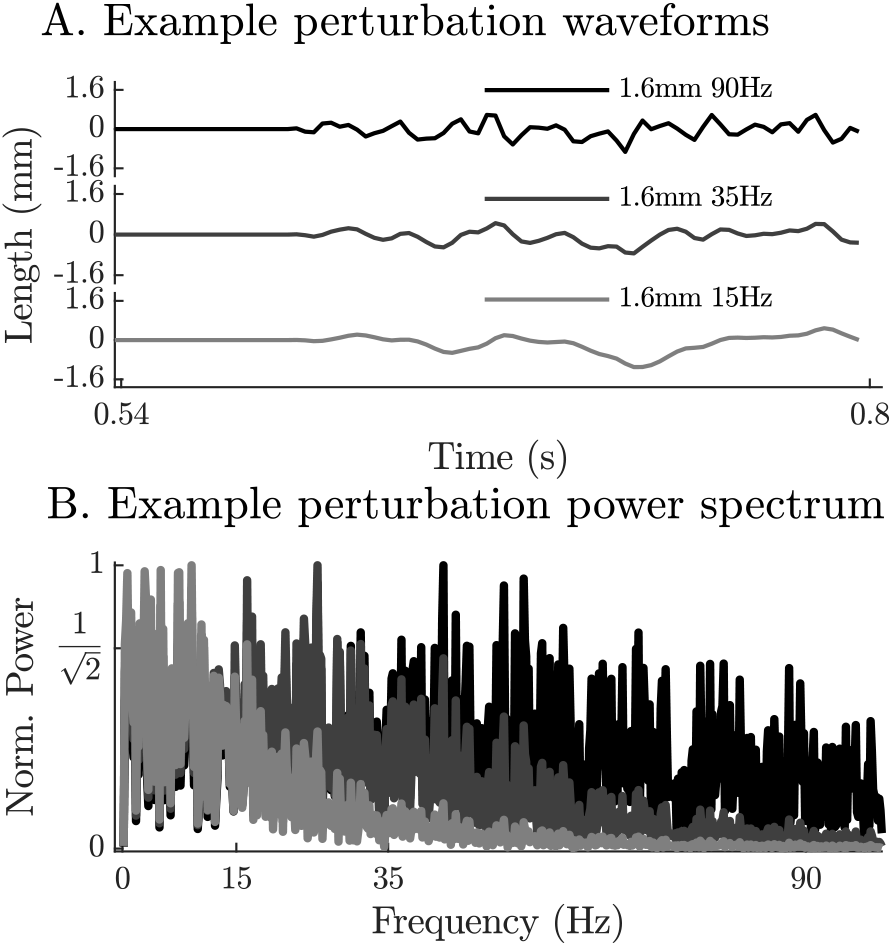
The perturbation waveforms are constructed by generating a series of pseudo-random numbers, padding the ends with zeros, by filtering the signal using a 2nd order low-pass filter (wave forms with -3dB cut-off frequencies of 90 Hz, 35 Hz and 15 Hz appear in A.) and finally by scaling the range to the desired limit (1.6mm in A.). Although the power spectrum of the resulting signals is highly variable, the filter ensures that the frequencies beyond the -3dB point have less than half their original power (B.).

When coupled with an elastic-tendon, the 15 Hz perturbations show that neither model can match the VAF of Kirsch et al.’s analysis [5] (compare Fig. 6A to G), while at 90Hz the VEXAT model reaches a VAF of 89% (Fig. 6D) which is within the range of 78-99% reported by Kirsch et al. [5]. In contrast, the Hill model’s VAF at 90 Hz remains low at 58% (Fig. 6J). While the VEXAT model has a gain profile in the frequency-domain that closer to Kirsch et al.’s data [5] than the Hill model (compare Fig. 6B to H and E to K), both models have a greater phase shift than Kirsch et al.’s data [5] at low frequencies (compare Fig. 6C to I and F to L). The phase response of the VEXAT model to the 90 Hz perturbation (Fig. 6F) shows the consequences of Eqn. 16: at low frequencies the phase response of the VEXAT model is similar to that of the Hill model, while at higher frequencies the model’s response becomes similar to a spring-damper. This frequency dependent response is a consequence of the first term in Eqn. 16: the value of *τ* ^S^ causes the response of the model to be similar to a Hill model at lower frequencies and mimic a spring-damper at higher frequencies. Both models show the same perturbation-dependent phase-response, as the damping coefficient of best fit increases as the perturbation bandwidth decreases: compare the damping coefficient of best fit for the 15Hz and 90Hz profiles for the VEXAT model (listed on Fig. 6A. and D.) and the Hill model (listed on Fig. 6G. and J., respectively).

**Figure 6.**
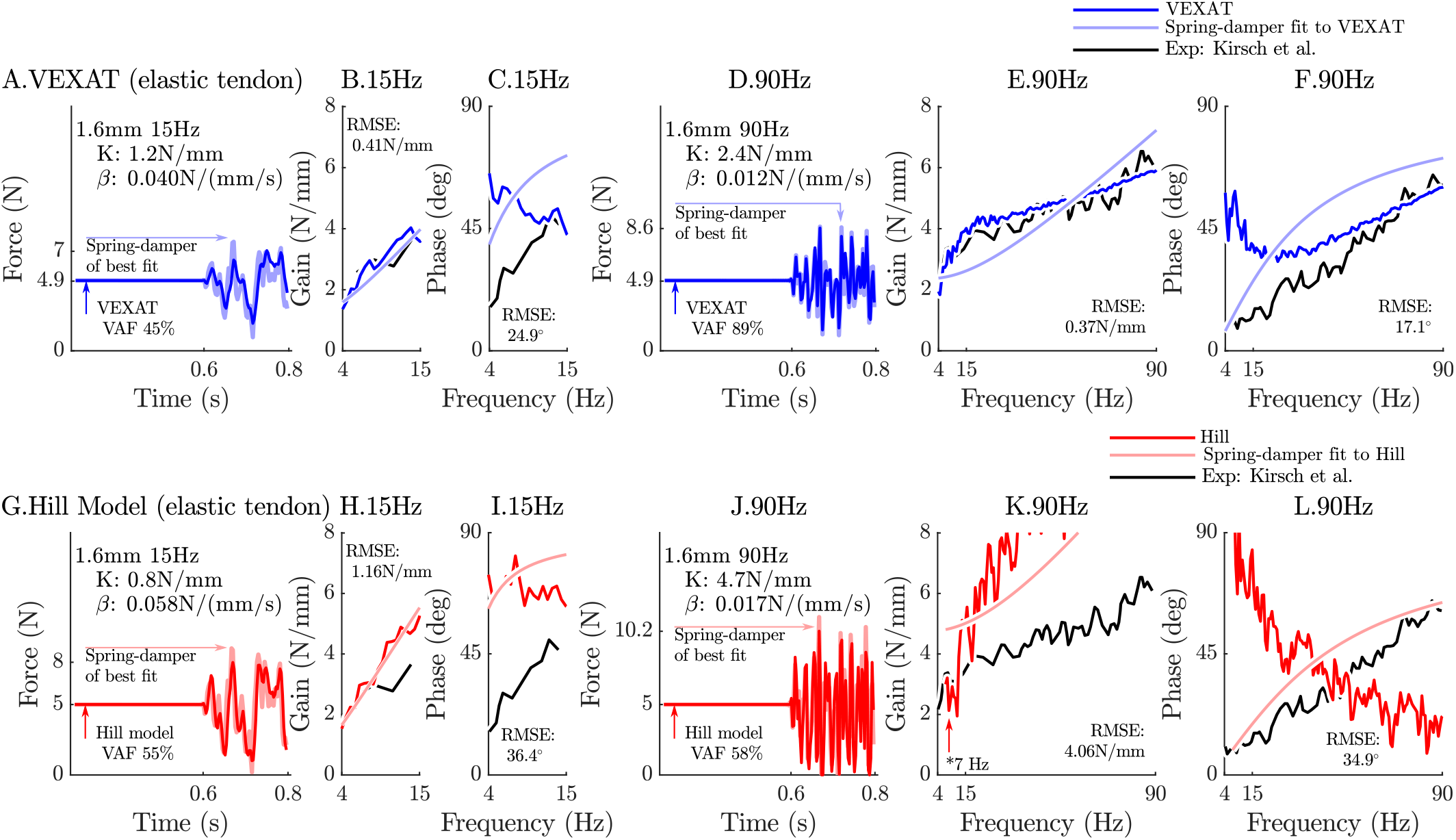
The 15 Hz perturbations show that the VEXAT model’s performance is mixed: in the time-domain (A.) the VAF is lower than the 78-99% analyzed by Kirsch et al. [5]; the gain response (B.) follows the profile in Figure 3 of Kirsch et al. [5], while the phase response differs (C.). The response of the VEXAT model to the 90 Hz perturbations is much better: a VAF of 91% is reached in the time-domain (D.), the gain response follows the response of the cat soleus analyzed by Kirsch et al. [5], while the phase-response follows biological muscle closely for frequencies higher than 30 Hz. Although the Hill’s time-domain response to the 15 Hz signal has a higher VAF than the VEXAT model (G.), the RMSE of the Hill model’s gain response (H.) and phase response (I.) shows it to be more in error than the VEXAT model. While the VEXAT model’s response improved in response to the 90 Hz perturbation, the Hill model’s response does not: the VAF of the time-domain response remains low (J.), neither the gain (K.) nor phase responses (L.) follow the data of Kirsch et al. [5]. Note that the Hill model’s 90 Hz response was so nonlinear that the lowest frequency analyzed had to be raised from 4 Hz to 7 Hz to satisfy the criteria that (*C*_*xy*_)^2^ *≥* 0.67.

The closeness of each model’s response to the spring-damper of best fit changes when a rigid-tendon is used instead of an elastic-tendon. While the VEXAT model’s response to the 15 Hz and 90 Hz perturbations improves slightly (compare Fig. 6A-F to Fig. 16A-F in Appendix F), the response of the Hill model to the 15 Hz perturbation changes dramatically with the time-domain VAF rising from 55% to 85% (compare Fig. 6G-L to Fig. 16G-L in Appendix F). Although the Hill model’s VAF in response to the 15 Hz perturbation improved, the frequency response contains mixed results: the rigid-tendon Hill model’s gain response is better (Fig. 16H in Appendix F), while the phase response is worse in comparison to the elastic-tendon Hill model. While the rigid-tendon Hill model produces a better time-domain response to the 15 Hz perturbation than the elastic-tendon Hill model, this improvement has been made with a larger phase shift between force and length than biological muscle [5].

The gain and phase profiles of both models deviate from the spring-damper of best fit due to the presence of nonlinearities, even for small perturbations. Some of the VEXAT model’s nonlinearities in this experiment come from the tendon model (compare Fig. 6A-F to Fig. 16A-F in Appendix F), since the response of the VEXAT model with a rigid-tendon stays closer to the spring-damper of best fit. The Hill model’s nonlinearities originate from the underlying expressions for stiffness and damping of the Hill model, which are particularly nonlinear with a rigid-tendon model (Fig. 16G-L in Appendix F) The stiffness of a Hill model’s CE

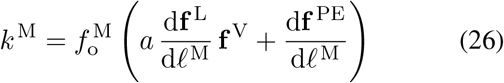

is heavily influenced by the partial derivative of 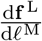 which has a region of negative stiffness. Although 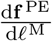 is well approximated as being linear for small length changes, 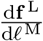 changes sign across 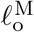. The damping of a Hill model’s CE

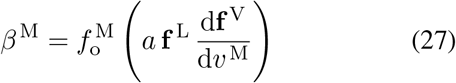

also suffers from high degrees of nonlinearity for small perturbations about *v* ^M^ = 0 since the slope of 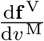 is positive and large when shortening, and positive and small when lengthening (Fig. 2B). While Eqns. 26 and 27 are mathematically correct, the negative stiffness and wide ranging damping values predicted by these equations do not match experimental data [5]. In contrast, the stiffness

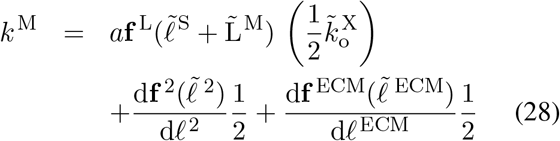

and damping

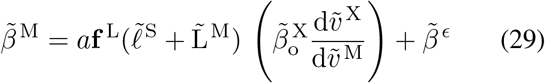

of the VEXAT’s CE do not change so drastically because these terms do not depend on the slope of the force-length relation, or the force-velocity relation (see Appendix B.4 for derivation).

By repeating the stochastic perturbation experiments across a range of isometric forces, Kirsch et al. [5] were able to show that the stiffness and damping of a muscle varies linearly with the active tension it develops (see Figure 12 of [5]). We have repeated our simulations of Kirsch et al.’s [5] experiments at ten nominal forces (spaced evenly between 2.5N and 11.5 N) and compared how the VEXAT model and the Hill model’s stiffness and damping coefficients compare to Figure 12 of Kirsch et al. [5] (Fig. 7). The stiffness and damping profile of the VEXAT model deviates a little from Kirsch et al.’s data [5] because XE’s dynamics at 35 Hz are still influenced by the Hill model embedded in Eqn. 16 (see Appendix B.4). Despite this, the VEXAT model develops similar stiffness and damping profile with either a viscoelastic-tendon (Fig. 7A & B) or a rigid-tendon (Fig. 7C & D). In contrast, when the Hill model is coupled with an elastic-tendon both its stiffness and damping are larger than Kirsch et al.’s data [5] at the higher tensions (Fig. 7A and B). This pattern changes when simulating a Hill model with a rigid-tendon: the model’s stiffness is slightly negative (Fig. 7C), while the model’s final damping coefficient is nearly three times the value measured by Kirsch et al. (Fig. 7D). Though a negative stiffness may seem surprising, Eqn. 26 shows a negative stiffness is possible at the nominal CE length of these simulations: just past 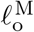 the slope of the active force-length curve is negative and the slope of the passive force-length curve is negligible. The tendon model also affects the VAF of the Hill model to a large degree: the elastic-tendon Hill model has a low VAF 30-51% (Fig. 7A & B) while the rigid-tendon Hill model has a much higher VAF of 86%. Although the VAF of the rigid-tendon Hill model is acceptable, these forces are being generated in a completely different manner than those obtained from biological muscle, as Kirsch et al.’s data [5] indicate (Fig. 7C and D).

**Figure 7.**
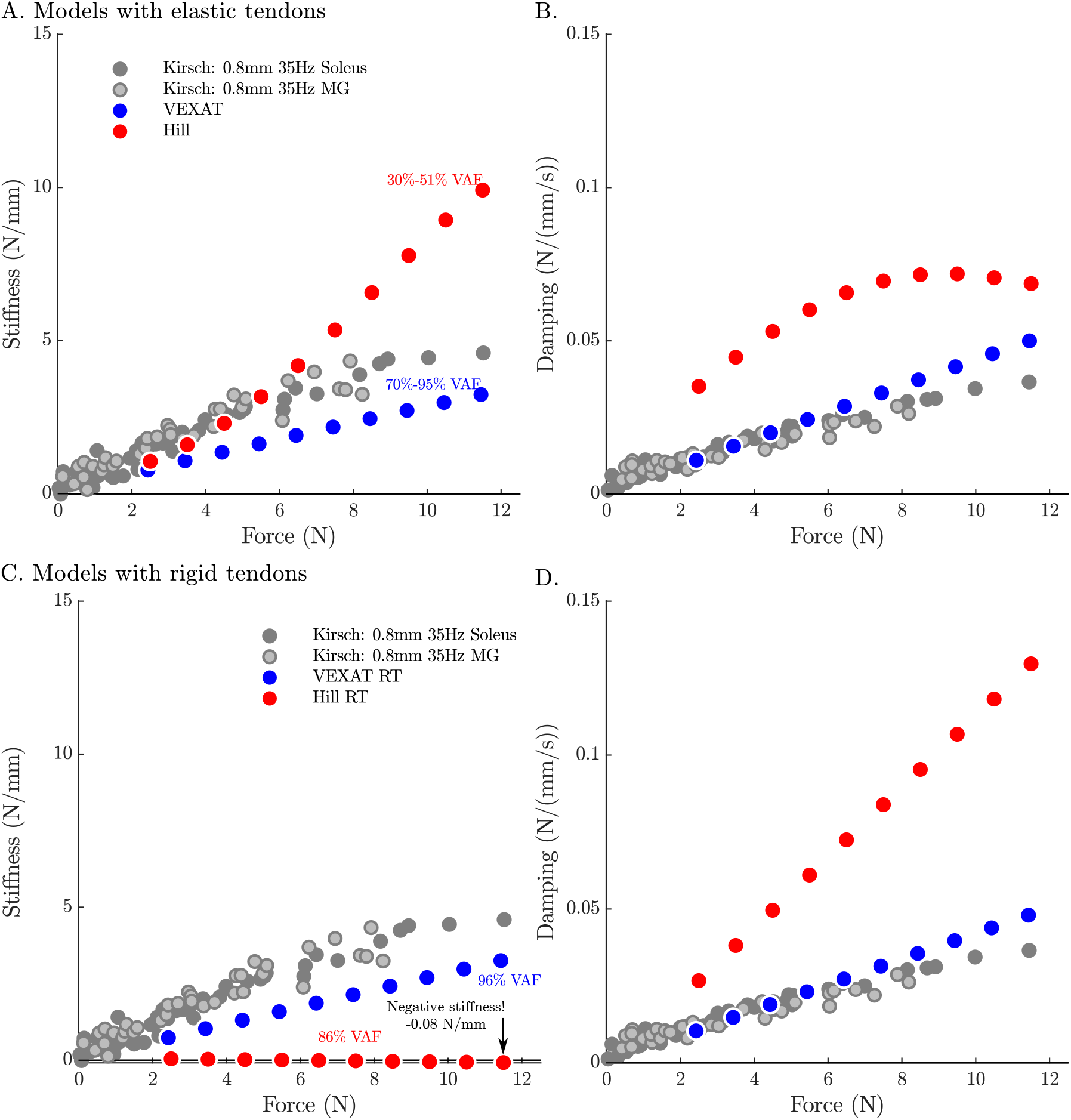
When coupled with an elastic-tendon, the stiffness (A.) and damping (B.) coefficients of best fit of both the VEXAT model and a Hill model increase with the tension developed by the MTU. However, both the stiffness and damping of the elastic-tendon Hill model are larger than Kirsch et al.’s coefficients (from Figure 12 of [5]), particularly at higher tensions. When coupled with rigid-tendon the stiffness (C.) and damping (D.) coefficients of the VEXAT model remain similar, as the values for 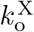 and 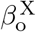 have been calculated to take the tendon model into account (see Appendix B.4 for details). In contrast, the stiffness and damping coefficients of the rigid-tendon Hill model differ dramatically from the elastic-tendon Hill model: while the elastic-tendon Hill model is too stiff and damped, the rigid-tendon Hill model is not stiff enough (compare A. to C.) and far too damped (compare B. to D.). Coupling the Hill model with a rigid-tendon increases the VAF from 30-51% to 86% but this improved accuracy is made using stiffness and damping that deviates from that of biological muscle [5].

When the VAF of the VEXAT and Hill model is evaluated across a range of nominal tensions (ten values from 2.5 to 11.5N), frequencies (15 Hz, 35 Hz, and 90 Hz), amplitudes (0.4mm, 0.8mm, and 1.6mm), and tendon types (rigid and elastic) two things are clear: first, that the VEXAT model’s 64-100% VAF is close to the 78-99% VAF reported by Kirsch et al. [5] while the Hill model’s 28-95% VAF differs (Fig. 8); and second, that there are systematic vari-ations in VAF, stiffness, and damping across the different perturbation magnitudes and frequencies (see Tables 5 and 5 in Appendix E). Both models produce worse VAF values when coupled with an elastic-tendon (Fig. 8A, B, and C), though the Hill model is affected most: the mean VAF of the elastic-tendon Hill model is 67% lower than the mean VAF of the rigid-tendon model for the 0.4 mm 15 Hz perturbations (Fig. 8A). While the VEXAT model’s lowest VAF occurs in response to the low frequency perturbations (Fig. 8A) with both rigid and elastic-tendons, the Hill model’s lowest VAF varies with both tendon type and frequency: the rigid-tendon Hill model has its lowest VAF in response to the 1.6 mm 90 Hz perturbations (Fig. 8C) while the elastic-tendon Hill model has its lowest VAF in response to the 0.4 mm 15 Hz perturbations (Fig. 8A). It is unclear if biological muscle displays systematic shifts in VAF since Kirsch et al. [5] did not report the VAF of each trial.

**Figure 8.**
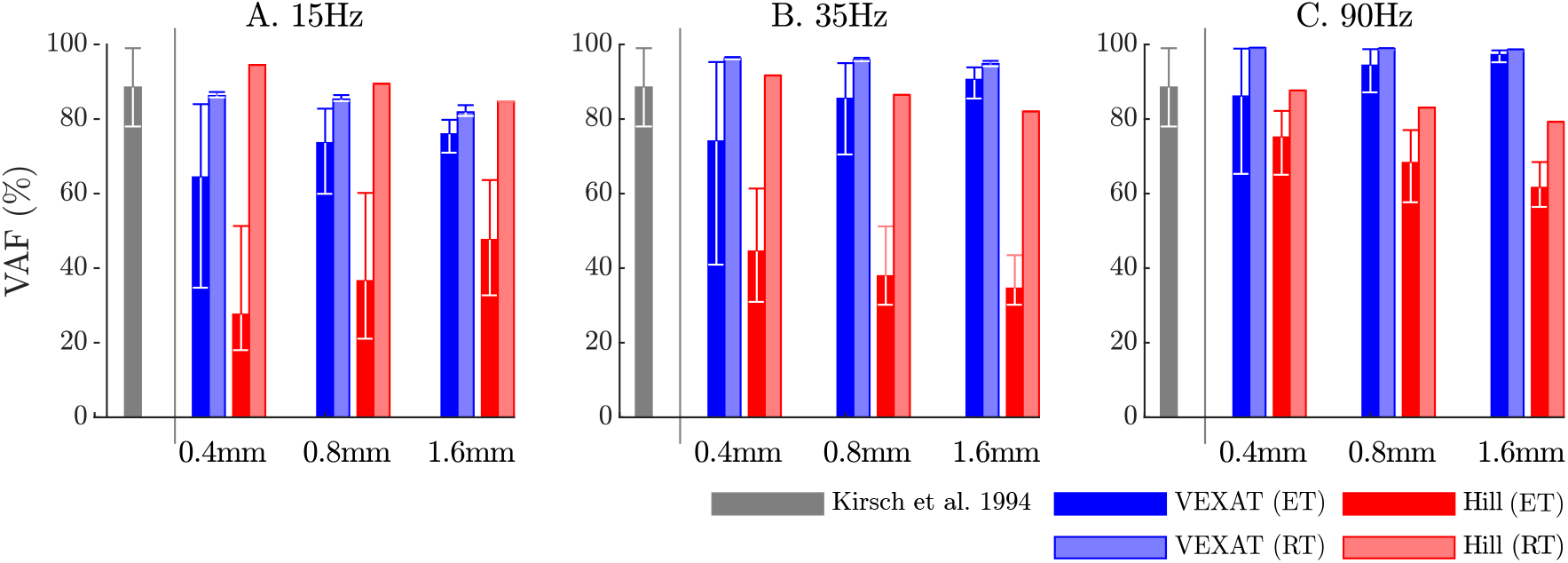
Kirsch et al. [5] noted that the VAF of the spring-damper model of best fit captured between 78-99% across all experiments. We have repeated the perturbation experiments to evaluate the VAF across a range of conditions: two different tendon models, three perturbation bandwidths (15 Hz, 35 Hz, and 90 Hz), three perturbation magnitudes (0.4 mm, 0.8 mm and 1.6 mm), and ten nominal force values (spaced evenly between 2.5N and 11.5N). Each bar in the plot shows the mean VAF across all ten nominal force values, with the whiskers extending to the minimum and maximum value that occurred in each set. The mean VAF of the VEXAT model changes by up to 36% depending on the condition, with the lowest mean VAF occurring in response to the 1.6 mm 15 Hz perturbation with an elastic-tendon (A.), and the highest mean VAF occurring in response to the 90 Hz perturbations with the rigid-tendon (C.). In contrast, the mean VAF of the Hill model varies by up to 67% depending on the condition, with the lowest VAF occurring in the 15 Hz 0.4 mm trial with the elastic-tendon (A.), and the highest value VAF occurring in the 15 Hz 0.4 mm trial with the rigid-tendon (A.).

**Figure 9.**
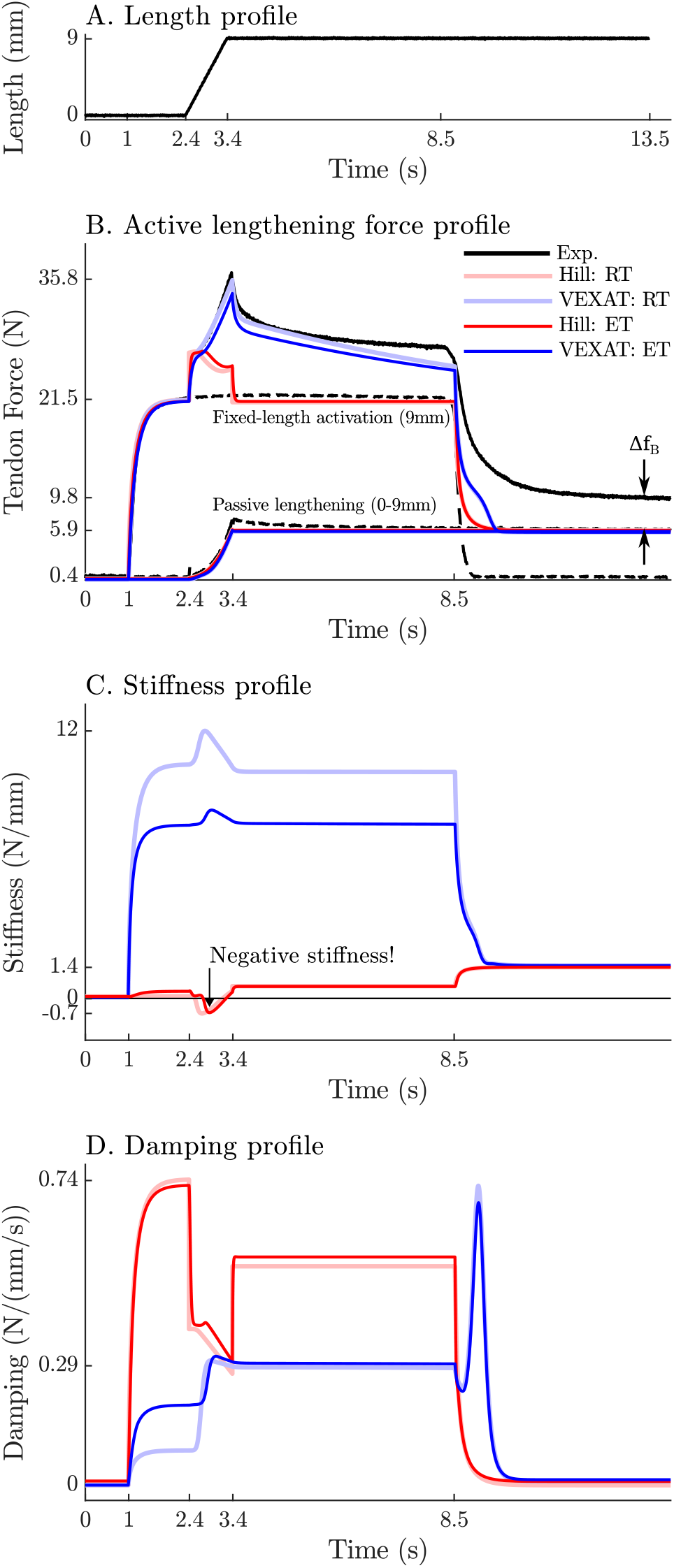
Herzog and Leonard [7] actively lengthened (A.) cat soleus muscles on the descending limb of the force-length curve (where 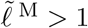 in Fig. 2A) and measured the force response of the MTU (B.). After the initial transient at 2.4s the Hill model’s output force drops (B.) because of the small region of negative stiffness (C.) created by the force-length curve. In contrast, the VEXAT model develops steadily increasing forces between 2.4 − 3.4s and has a consistent level of stiffness (C.) and damping (D.).

**Figure 10.**
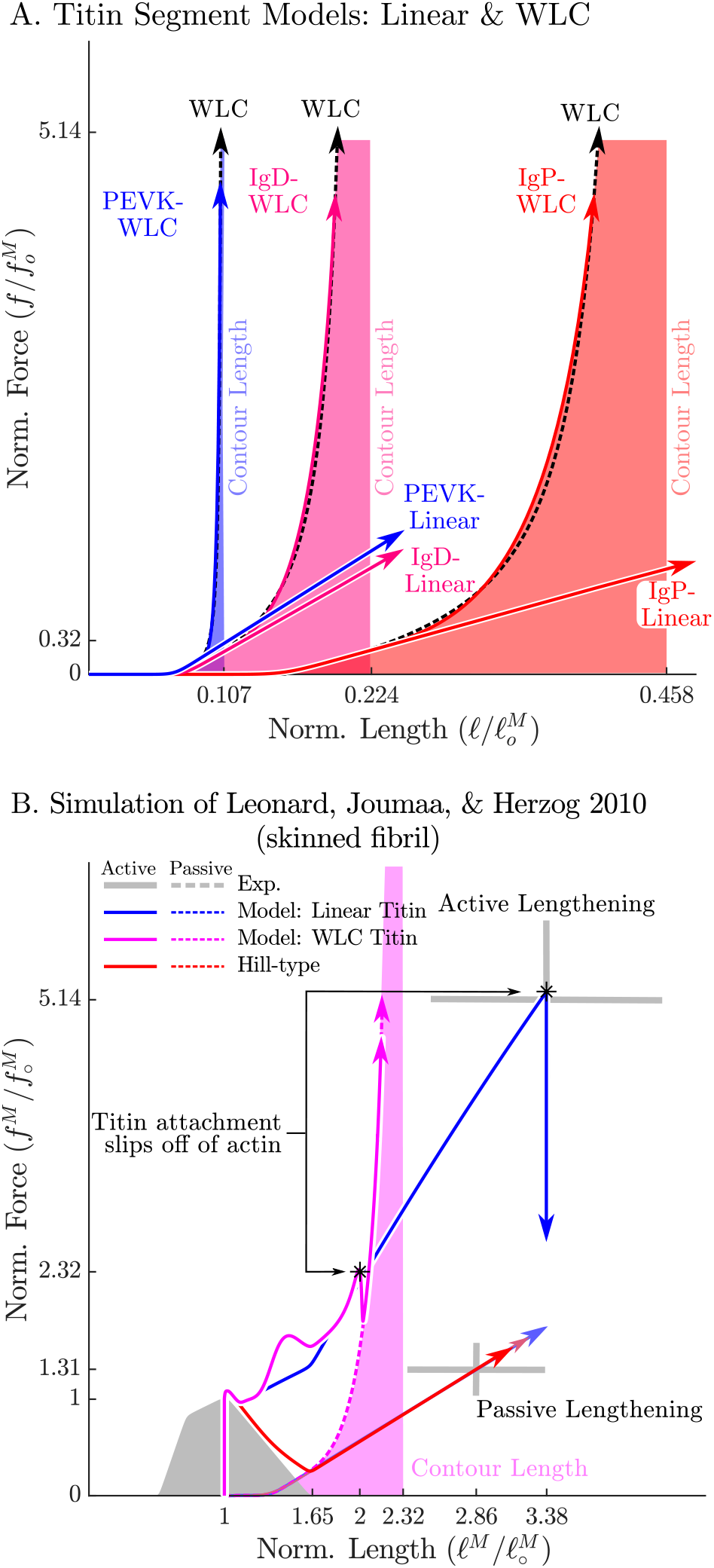
In the VEXAT model we consider two different versions of the force-length relation for each titin segment (A): a linear extrapolation, and a WLC model extrapolation. Leonard et al. [8] observed that active myofibrils continue to develop increasing amounts of tension beyond actin-myosin overlap (B, grey lines with *±*1 standard deviation shown). When this experiment is replicated using the VEXAT model (B., blue & magenta lines) and a Hill model (C. red lines), only the VEXAT model with the linear extrapolated titin model is able to replicate the experiment with the titin-actin bond slipping off of the actin filament at 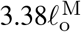 .

### 3.2 Active lengthening on the descending limb

We now turn our attention to the active lengthening insitu experiments of Herzog and Leonard [7]. During these experiments cat soleus muscles were actively lengthened by modest amounts 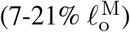 starting on the descending limb of the active-force-length curve (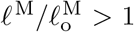 in Fig. 2A). This starting point was chosen specifically because the stiffness of a Hill model may actually change sign and become negative because of the influence of the active-force-length curve on *k* ^M^ as shown in Eqn. 26 as *ℓ* ^M^ extends beyond 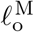. Herzog and Leonard’s [7] experiment is important for showing that biological muscle does not exhibit negative stiffness on the descending limb of the active-force-length curve. In addition, this experiment also highlights the slow recovery of the muscle’s force after stretching has ceased, and the phenomena of passive force enhancement after stimulation is removed. Here we will examine the 9 mm/s ramp experiment in detail because the simulations of the 3 mm/s and 27 mm/s ramp experiments produces similar stereotypical patterns (see Appendix G for details).

When Herzog and Leonard’s [7] active ramp-lengthening (Fig. 9A) experiment is simulated, both models produce a force transient initially (Fig. 9B), but for different reasons. The VEXAT model’s transient is created when the lumped crossbridge spring (the 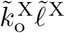 term in Eqn. 15) is stretched. In contrast, the Hill model’s transient is produced, not by spring forces, but by damping produced by the force-velocity curve as shown in Eqn. 26.

After the initial force transient, the response of the two models diverges (Fig. 9B): the VEXAT model continues to develop increased tension as it is lengthened, while the Hill model’s tension drops before recovering. The VEXAT model’s continued increase in force is due to the titin model: when activated, a section of titin’s PEVK region remains approximately fixed to the actin element (Fig. 1C). As a result, the *ℓ* ^2^ element (composed of part of PEVK segment and the distal Ig segment) continues to stretch and generates higher forces than it would if the muscle were being passively stretched. While both the elastic and rigid-tendon versions of the VEXAT model produce the same stereotypical ramp-lengthening response (Fig. 9B), the rigid-tendon model develops slightly more tension because the strain of the MTU is solely borne by the CE.

In contrast, the Hill model develops less force during lengthening when it enters a small region of negative stiffness (Fig. 9B and C) because the passive-force-length curve is too compliant to compensate for the negative slope of the active force-length curve. Similarly, the damping coefficient of the Hill model drops substantially during lengthening (Fig. 9D). Equation 27 and Figure 2B shows the reason that damping drops during lengthening: *d* **f** ^V^*/dv* ^M^, the slope of the line in Fig. 2B, is quite large when the muscle is isometric and becomes quite small as the rate of lengthening increases.

After the ramp stretch is completed (at time 3.4 seconds in Fig. 9B), the tension developed by the cat soleus recovers slowly, following a profile that looks strikingly like a first-order decay. The large damping coefficient acting between the titin-actin bond slows the force recovery of the VEXAT model. We have tuned the value of 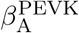 to 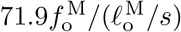 for the elastic-tendon model, and 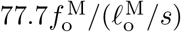 for the rigid-tendon model, to match the rate of force decay of the cat soleus in Herzog and Leonard’s data [7]. The Hill model, in contrast, recovers to its isometric value quite rapidly. Since the Hill model’s force enhancement during lengthening is a function of the rate of lengthening, when the lengthening ceases, so too does the force enhancement.

Once activation is allowed to return to zero, Herzog and Leonard’s data shows that the cat soleus continues to develop a tension that is Δ*f*_*B*_ above passive levels (Fig. 9B for *t >* 8.5*s*). The force Δ*f*_*B*_ is known as passive force enhancement, and is suspected to be caused by titin bind-ing to actin [76]. Since we model titin-actin forces using an activation-dependent damper, when activation goes to zero our titin model becomes unbound from actin. As such, both our model and a Hill model remain Δ*f*_*B*_ below the experimental data of Herzog and Leonard (Fig. 9B) after lengthening and activation have ceased.

### 3.3 Active lengthening beyond actin-myosin overlap

One of the great challenges that remains is to decompose how much tension is developed by titin (Fig. 1C) separately from myosin (Fig. 1B) in an active sarcomere. Leonard et al.’s [8] active-lengthening experiment provides some insight into this force distribution problem because they recorded active forces both within and far beyond actin-myosin overlap. Leonard et al.’s [8] data shows that active force continues to develop linearly during lengthening, beyond actin-myosin overlap, until mechanical failure. When activated and lengthened, the myofibrils failed at a length of 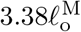 and force of 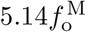, on average. In contrast, during passive lengthening myofibrils failed at a much shorter length of 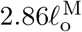 with a dramatically lower tension of of 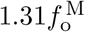. To show that the extraordinary forces beyond actin-myosin overlap can be ascribed to titin, Leonard et al. [8] repeated the experiment but deleted titin using trypsin: the titin-deleted myofibrils failed at short lengths and insignificant stresses. Using the titin model of Eqn. 20 (Fig. 1A) as an interpretive lens, the huge forces developed during active lengthening would be created when titin is bound to actin leaving the distal segment of titin to take up all of the strain. Conversely, our titin model would produce lower forces during passive lengthening because the proximal Ig, PEVK, and distal Ig regions would all be lengthening together (Fig. 3A).

Since Leonard et al.’s experiment [8] was performed on skinned rabbit myofibrils and not on whole muscle, both the VEXAT and Hill models had to be adjusted prior to simulation (see Appendix H for parameter values). To simulate a rabbit myofibril we created a force-length curve [75] consistent with the filament lengths of rabbit skeletal muscle [60] (1.12*µ*m actin, 1.63*µ*m myosin, and 0.07*µ*m z-line width) and fit the force-length relations of the two titin segments to be consistent with the structure measured by Prado et al. [58] of rabbit psoas titin consisting of a 70%-30% mix of a 3300kD and a 3400kD titin isoform (see Appendix B.3 for fitting details and Appendix H for parameter values). Since this is a simulation of a fibril, we used a rigid-tendon of zero length (equivalent to ignoring the tendon), and set the pennation angle to zero.

As mentioned in Sec. 2, because this experiment includes extreme lengths, we consider two different force-length relations for each segment of titin (Fig. 10A): a linear extrapolation, and an extension that follows the WLC model. While both versions of the titin model are identical up to 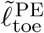, beyond 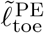 the WLC model continues to develop increasingly large forces until all of the Ig domains and PEVK residues have been unfolded and the segments of titin reach a physical singularity: at this point the Ig domains and PEVK residues cannot be elongated any further without breaking molecular bonds (see Appendix B.3 for details). Our preliminary simulations indicated that the linear titin model’s titin-actin bond was not strong enough to support large tensions, and so we increased the value of 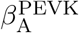 from 71.9 to 975 (compare Tables 1 and 6 section H).

The Hill model was similarly modified, with the pennation angle set to zero and coupled with a rigid-tendon of zero length. Since the Hill model lacks an ECM element the passive-force-length curve was instead fitted to match the passive forces produced in Leonard et al.’s data [8]. No adjustments were made to the active elements of the Hill model.

When the slow active stretch (0.1*µm*/sarcomere/s) of Leonard et al.’s experiment is simulated [8] only the VEXAT model with the linear titin element can match the experimental data of Leonard et al. [8] (Fig. 10B). The Hill model cannot produce active force for lengths greater than 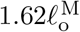 since the active force-length curve goes to zero (Fig. 2A) and the model lacks any element capable of producing force beyond this length. In contrast, the linear titin model continues to develop active force until a length of 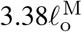 is reached, at which point the titin-actin bond is pulled off the end of the actin filament and the active force is reduced to its passive value.

The WLC titin model is not able to reach the extreme lengths observed by Leonard et al. [8]. The distal segment of the WLC titin model approaches its contour length early in the simulation and ensures that the the titin-actin bond is dragged off the end of the actin filament at 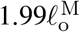 (Fig. 10B). After 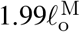 (Fig. 10B), the tension of the WLC titin model drops to its passive value but continues to increase until the contour lengths of all of the segments of titin are reached at 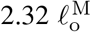. Comparing the response of the linear model to the WLC titin model two things are clear: the linear titin model more faithfully follows the data of Leonard et al. [8], but does so with titin segment lengths that exceed the maximum contour length expected for the isoform of titin in a rabbit myofibril.

This simulation has also uncovered a surprising fact: the myofibrils in Leonard et al.’s [8] experiments do not fail at 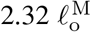, as would be expected by the WLC model of titin, but instead reach much greater lengths (Fig. 2B). Physically, it may be possible for a rabbit myofibril to reach these lengths (without exceeding the contour lengths of the proximal Ig, PEVK, and distal Ig segments) if the bond between the distal segment of titin and myosin breaks down. This would allow the large Ig segment, that is normally bound to myosin, to uncoil and continue to develop the forces observed by Leonard et al. [8]. Unfortunately the mechanism which allowed the samples in Leonard et al.’s experiments to develop tension beyond titin’s contour length remains unknown.

### 3.4 Force-length and force-velocity

Although the active portion of the Hill model is embedded in Eqn. 16, it is not clear if the VEXAT model can still replicate Hill’s force-velocity experiments [9] and Gordon et al.’s [10] force-length experiments. Here we simulate both of these experiments using the cat soleus model that we have used for the simulations described in Sec. 3.1 and compare the results to the force-length and force-velocity curves that are used in the Hill model and in Eqn. 16 of the VEXAT model.

Hill’s force-velocity experiment [9] is simulated by activating the model, and then by changing its length to follow a shortening ramp and a lengthening ramp. During shortening experiments, the CE shortens from 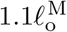 to 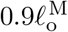 with the measurement of active muscle force is made at 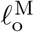 Lengthening experiments are similarly made by measuring muscle force mid-way through a ramp stretch that begins at 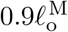 and ends at 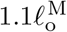. When an elastic-tendon model is used, we carefully evaluate initial and terminal path lengths to accommodate for the stretch of the tendon so that the CE still shortens from 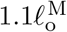 to 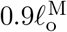 and lengthens from 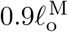 to 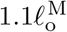.

The VEXAT model produces forces that differ slightly from the **f** ^V^ that is embedded in Eqn. 16 while the Hill model reproduces the curve (Fig. 11). The maximum shortening velocity of the VEXAT model is slightly weaker than the embedded curve due to the series viscoelasticity of the XE element. Although the model can be made to converge to the **f** ^V^ curve more rapidly by decreasing *τ* ^S^ this has the undesirable consequence of degrading the low-frequency response of the model during Kirsch et al.’s experiments [5] (particularly Fig. 6C., and F.). These small differences can be effectively removed by scaling 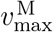 by s^V^ (Fig. 11A has s^V^ = 0.95) to accommodate for the small decrease in force caused by the viscoelastic XE element.

**Figure 11.**
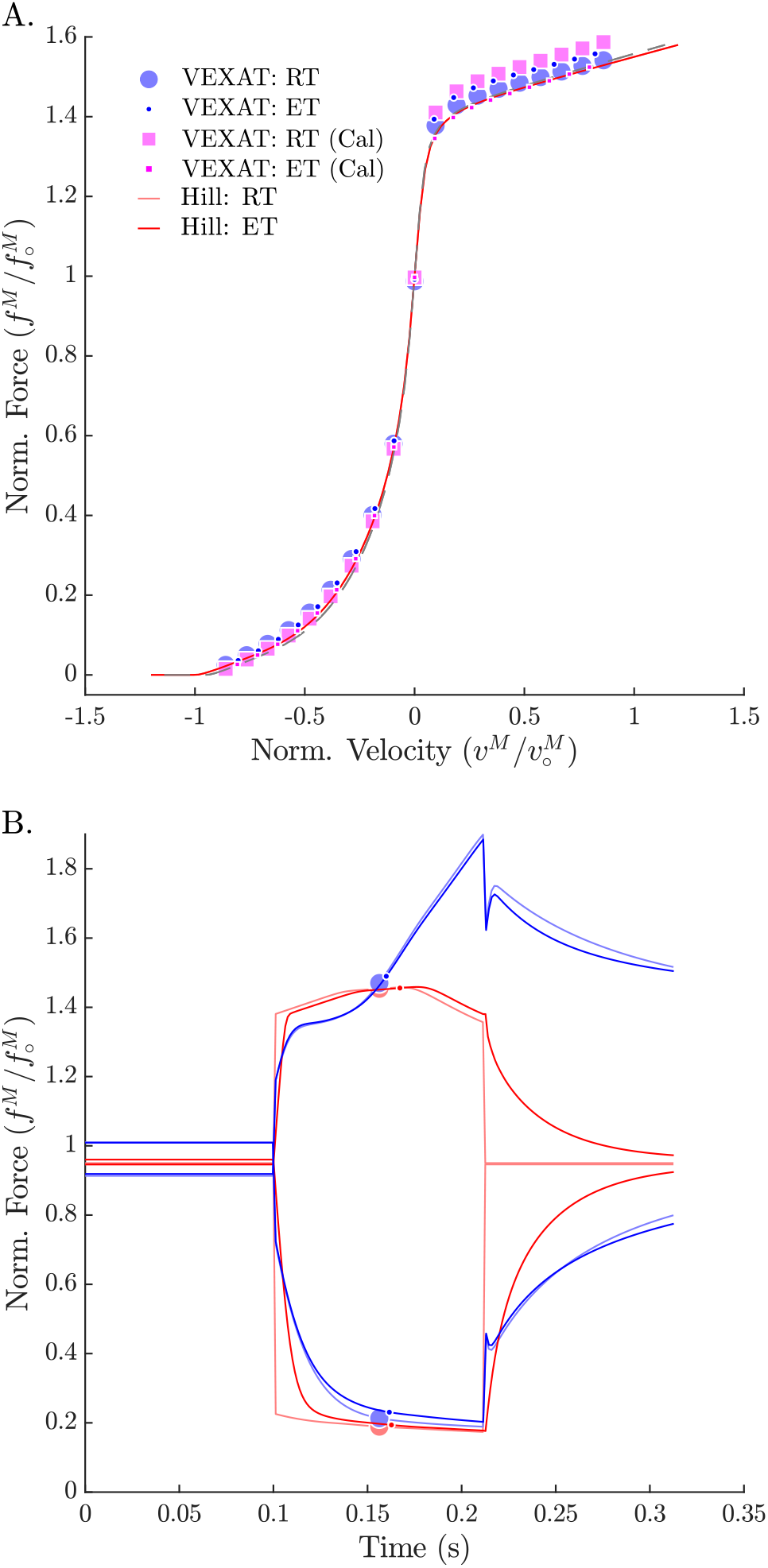
When Hill’s [9] force-velocity experiment is simulated (A.), the VEXAT model produces a force-velocity profile (blue dots) that approaches zero more rapidly during shortening than the embedded profile **f** ^V^(·) (red lines). By scaling 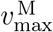 by 0.95 the VEXAT model (magenta squares) is able to closely follow the force-velocity curve of the Hill model. While the force-velocity curves between the two models are similar, the time-domain force response of the two models differs substantially (B.). The rigid-tendon Hill model exhibits a sharp nonlinear change in force at the beginning (0.1s) and ending (0.21s) of the ramp stretch.

**Figure 12.**
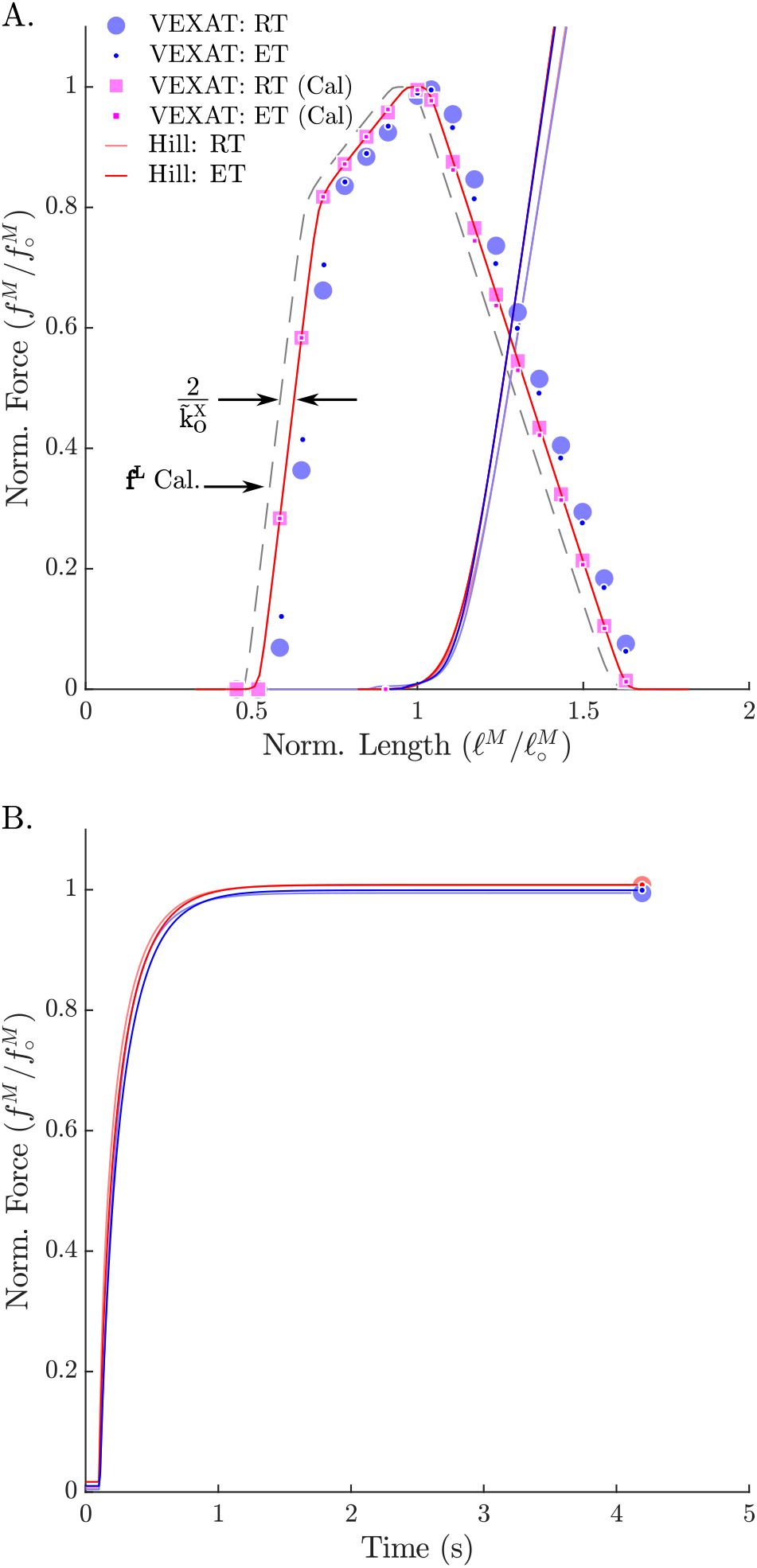
When Gordon et al.’s [10] passive and active force-length experiments are simulated, the VEXAT model (blue dots) and the Hill model (red lines) produce slightly different force-length curves (A.) and force responses in the time-domain (B.). The VEXAT model produces a right shifted active force-length curve, when compared to the Hill model due to the series elasticity of the XE element. By shifting the underlying curve by 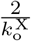 to the left the VEXAT model (magenta squares) can be made to exactly match the force-length characteristic of the Hill model.

Gordon et al.’s [10] force-length experiments were simulated by first passively lengthening the CE, and next by measuring the active force developed by the CE at a series of fixed lengths. Prior to activation, the passive CE was simulated for a brief period of time in a passive state to reduce any history effects due to the active titin element. To be consistent with Gordon et al.’s [10] experiment, we subtracted off the passive force from the active force before producing the active-force-length profile.

The simulation of Gordon et al.’s [10] experiment shows that the VEXAT model (Fig. 12A, blue dots) produces a force-length profile that is shifted to the right of the Hill model (Fig. 12A, red line) due to the series elasticity in-troduced by the XE. We can solve for the size of this right-wards shift by noting that Eqn. 16 will drive the 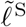 to a length such that the isometric force developed by the XE is equal to that of the embedded Hill model

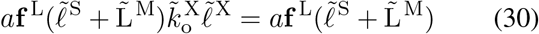

allowing us to solve for

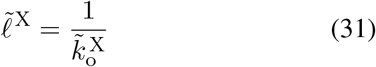

the isometric strain of the XE. Since there are two viscoelastic XE elements per CE, the VEXAT model has an active force-length characteristic that shifted to the right of the embedded **f** ^L^ curve by a constant 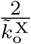. This shift, Δ^L^, can be calibrated out of the model (Fig. 12 upper plot, magenta squares) by adjusting the **f** ^L^(·) curve so that it is 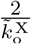 to the left of its normal position. Note that all simulations described in the previous sections made use of the VEXAT model with the calibrated force-length relation and the calibrated force-velocity relation.

## 4 Discussion

A muscle model is defined by the experiments it can replicate and the mechanisms it embodies. We have developed the VEXAT muscle model to replicate the force response of muscle to a wide variety of perturbations [5], [7], [8] while also retaining the ability to reproduce Hill’s force-velocity [9] experiment and Gordon et al.’s [10] force-length experiments. The model we have developed uses two mechanisms to capture the force response of muscle over a large variety of time and length scales: first, a viscoelastic crossbridge element that over brief time-scales appears as a spring-damper, and at longer time-scales mimics a Hill-model; second, a titin element that is capable of developing active force during large stretches.

The viscoelastic crossbridge and titin elements we have developed introduce a number of assumptions into the model. While there is evidence that the activation-dependent stiffness of muscle originates primarily from the stiffness of the attached crossbridges [62], the origins of the activation-dependent damping observed by Kirsch et al. [5] have not yet been established. We assumed that, since the damping observed by Kirsch et al. [5] varies linearly with activation, the damping originates from the attached crossbridges. Whether this damping is intrinsic or is due to some other factor remains to be established. Next, we have also assumed that the force developed by the XE converges to a Hill model [17] given enough time (Eqn. 16). A recent experiment of Tomalka et al. [80] suggests the force developed by the XE might decrease during lengthening rather than increasing as is typical of a Hill model [17]. If Tomalka et al.’s [80] observations can be replicated, the VEXAT model will need to be adjusted so that the the XE element develops less force during active lengthening while the active-titin element develops more force. Finally, we have assumed that actin-myosin sliding acceleration (due to crossbridge cycling) occurs when there is a force imbalance between the external force applied to the XE and the internal force developed by the XE as shown in Eqn. 16. This assumption is a departure from previous models: Hill-type models [16], [17] assume that the tension applied to the muscle instantaneously affects the actin-myosin sliding velocity; Huxley models [11] assume that the actin-myosin sliding velocity directly affects the rate of attachment and detachment of crossbridges.

The active titin model that we have developed makes assumptions similar to Rode et al. [38] and Schappacher-Tilp et al. [40]: some parts of the PEVK segment bond to actin, and this bond cannot do any positive work on titin. The assumption that the bond between titin and actin cannot do positive work means that titin cannot be actively preloaded: it can only develop force when it is stretched. In contrast, two mechanisms have been proposed that make it possible for titin to be preloaded by crossbridge cycling: Nishikawa’s [39] winding filament theory and DuVall et al.’s [44] titin entanglement hypothesis. If titin were significantly preloaded by crossbridge cycling, the titin load path would support higher forces and the myosin-actin load path would bear less force. Accordingly, the overall stiffness of the CE would be reduced, affecting our simulations of Kirsch et al. [5]: lower myosin-actin loads mean fewer attached crossbridges, since crossbridges are stiff in comparison to titin, the stiffness of the CE would decrease (see Appendix A). Hopefully experimental work will clarify if titin can be actively preloaded by crossbridges in the future.

Both the viscoelastic crossbridge and active titin elements include simple myosin-actin and titin-actin bond models that improve accuracy but have limitations. First, the viscoelastic crossbridge element has been made to represent a population of crossbridges in which the contribution of any single crossbridge is negligible. Though it may be possible for the XE model to accurately simulate a maximally activated single sarcomere (which has roughly 20 attached crossbridges per half sarcomere [12], [81]) the accuracy of the model will degrade as the number of attached cross-bridges decreases. When only a single crossbridge remains, the XE model’s output will be inaccurate because it can only generate force continuously while a real crossbridge generates force discretely each time it attaches to, and detaches from, actin. Next, we have used two equations, Eqns. 16 and 21, that assume myosin-actin and titin-actin interactions are temperature-invariant and scale linearly with size (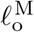 and 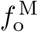). In contrast, myosin-actin interactions and some titin-actin interactions are temperature-sensitive [82], [83] and may not scale linearly with size. In Sec. 3.3 we had to adjust the active titin damping parameter, 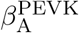 to simulate myofibril experiments [8], perhaps because the assumptions of temperature-invariance and size-linearity were not met: the initial value for 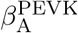 came from fitting to in-situ experimental data [7] from whole muscle that was warmer (35 − 36.5° C vs 20 − 21° C) and larger (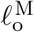 of 42.9 mm vs. 10 − 15 µm) than the myofibrils [8].

While the cat soleus XE and titin model parameters (Table 1 G, H, and I) can be used as rough default values, these parameters should be refit to accurately simulate muscle that differs in scale or temperature from cat soleus. Finally, the VEXAT model in its current form ignores phenomena related to submaximal contractions: the shift in the peak of the force-length relation [84], and the scaling of the maximum shortening velocity [85]. We hope to include these phenomena in a later version of the VEXAT model to more accurately simulate submaximal contractions.

The model we have proposed can replicate phenomena that occur at a wide variety of time and length scales: Kirsch et al.’s experiments [5] which occur over small time and length scales; and the active lengthening experiments of Herzog and Leonard [7] and Leonard et al. [8] which occur over physiological and supra-physiological length scales. In contrast, we have shown in Sec. 3.1 to 3.3 that a Hill-type model compares poorly to biological muscle when the same set of experiments are simulated. We expect that a Huxley model [11] is also likely to have difficulty reproducing Kirsch et al.’s experiment [5] because the model lacks an active damping element. Since titin was discovered [23] long after Huxley’s model was proposed [11], a Huxley model will be unable to replicate any experiment that is strongly influenced by titin such as Leonard et al.’s experiment [8].

Although there have been several more recent muscle model formulations proposed, none have the properties to simultaneously reproduce the experiments of Kirsch et al. [5], Herzog and Leonard [7], Leonard et al. [8], Hill [9], and Gordon et al. [10]. Linearized impedance models [14], [15] can reproduce Kirsch et al.’s experiments [5], but these models lack the nonlinear components needed to reproduce Gordon et al.’s force-length curve [10] and Hill’s force-velocity curve [9]. The models of Forcinito et al. [18], and Tahir et al. [41] have a structure that places a contractile element in series with an elastic-tendon. While this is a commonly used structure, at high frequencies the lack of damping in the tendon will drive the phase shift between length and force to approach zero. The measurements and model of Kirsch et al. [5], in contrast, indicate that the phase shift between length and force approaches ninety degrees with increasing frequencies. Though the Hill-type models of Haeufle et al. [20] and Günther et al. [22] have viscoelastic tendons, these models have no representation, of the viscoelasticity of the CE’s attached crossbridges. Similar to the Hill-type muscle model evaluated in this work [17], it is likely that models of Haeufle et al. [20] and Günther et al. [22] will not be able to match the frequency response of biological muscle. De Groote et al. [51], [52] introduced a short-range-stiffness element in parallel to a Hill model to capture the stiffness of biological muscle. While De Groote et al.’s [51], [52] formulation improves upon a Hill model it is unlikely to reproduce Kirsch et al.’s experiment [5] because we have shown in Sec. 3.1 that a Hill model has a frequency response that differs from biological muscle. Rode et al.’s [38] muscle model also uses a Hill model for the CE and so we expect that this model will have the same difficulties reproducing Kirsch et al.’s [5] experiment. Schappacher-Tilp et al.’s model [40] extends a Huxley model [11] by adding a detailed titin element. Similar to a Huxley model, Schappacher-Tilp et al.’s model [40] will likely have difficulty reproducing Kirsch et al.’s experiment [5] because it is missing an active damping element.

While developing this model, we have come across open questions that we hope can be addressed in the future. How do muscle stiffness and damping change across the force-length curve? Does stiffness and damping change with velocity? What are the physical origins of the active damping observed by Kirsch et al. [5]? What are the conditions that affect passive-force enhancement, and its release? In addition to pursuing these questions, we hope that other researchers continue to contribute experiments that are amenable to simulation, and to develop musculotendon models that overcome the limitations of our model. To help others build upon our work, we have made the source code of the model and all simulations presented in this paper available online^14^.

## Acknowledgements

Financial support is gratefully acknowledged from the Deutsche Forschungsgemeinschaft (DFG, German Research Foundation) under Germany’s Excellence Strategy (EXC 2075 – 390740016) through the Stuttgart Center for Simulation Science (SimTech), from DFG grant no. MI 2109/1-1, the Lighthouse Initiative Geriatronics by StMWi Bayern (Project X, grant no. 5140951), and the Natural Sciences and Engineering Research Council of Canada (RGPIN-2020-03920).

## A The stiffness of the actin-myosin and titin load paths

A single half-myosin can connect to the surrounding six actin filaments through 97.9 crossbridges. A 0.800 *µ*m half-myosin has a pair crossbridges over 0.700 *µ*m of its length every 14.3nm which amounts to 97.9 per halfmyosin [12]. Although 97.9 crossbridges does not make physical sense, here we will evaluate the stiffness of the CE assuming that fractional crossbridges exist and that attached crossbridges can be perfectly distributed among the 6 available actin filaments: the alternative calculation is more complicated and produces stiffness values that differ only in the 3^*rd*^ significant digit. Assuming a duty cycle of 20% [81] (values between 5-90% have been reported [86]), at full actin-myosin overlap there will be 19.6 crossbridges attached to the 6 surrounding actin filaments. Assuming that these 19.6 crossbridges are evenly distributed between the 6 actin filaments, each single actin will be attached to 3.26 attached crossbridges.

At full overlap, the Z-line is 1 actin filament length L^A^ (1.12 *µ*m in rabbits [60]) from the M-line. The average point of crossbridge attachment is in the middle of the halfmyosin at a distance of 0.45 *µ*m from the M-line (0.1 *µ*m is bare and 0.35 *µ*m is half of the remaining length), which is L^A^−0.45 µm from the Z-line. A single actin filament has a stiffness of 46 − 68 pN/nm [60] while a single crossbridge has a stiffness of 0.69 *±* 0.47 pN*/*nm [62]. Since stiffness scales inversely with length, actin’s stiffness between the Zline and the average attachment point is 81.8−121 pN/nm. Finally, the stiffness of each actin filament and its 3.26 attached crossbridges is 0.712 − 3.67 pN*/*nm and all 6 together have a stiffness of 4.27 − 22.0 pN*/*nm.

Myosin has a similar stiffness as a single actin filament [61], with the section between the average attachment point and the M-line having a stiffness of 76.9 − 113 pN/nm. The final active stiffness of half-sarcomere is 4.05 − 18.4 pN*/*nm which comes from comes from the series connection of the group of 6 actin filaments, with 19.6 crossbridges, and finally the single myosin filament. When this procedure is repeated assuming that only a single cross-bridge is attached the stiffness drops to 0.22−1.15 pN/nm, which is slightly less than the stiffness of a single cross-bridge ^15^.

The force-length profile of a single rabbit titin has been measured by Kellermayer et al. [87] using laser tweezers to apply cyclical stretches. By digitizing Fig. 4B (blue line) of Kellermayer et al. [87] we arrive at a stiffness for titin of 0.0058 − 0.0288 pN*/*nm at 2 *µ*m (for a total sarcomere length of 4 *µ*m or 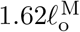), and 0.0505 − 0.0928 pN*/*nm at 4 *µ*m (8 *µ*m or 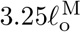). Since there are 6 titin filaments acting in parallel for each half-sarcomere, we end up with the total stiffness for titin ranging between 0.0348 − 0.173 pN*/*nm at 2 *µ*m and 0.303 − 0.557 pN*/*nm at 4 *µ*m.

When activated, the stiffness of our rabbit psoas linear-titin model (described in Sec. 3.3, fitted in Appendix B.3, and with the parameters shown in Appendix H) doubles, which would increase titin’s stiffness to 0.0696 − 0.346 pN*/*nm at 2 *µ*m and 0.606 − 1.11 pN*/*nm at 4 *µ*m.

Comparing the actin-myosin and titin stiffness ranges (Fig. 14) makes it clear that the stiffness of actin-myosin with 1 attached crossbridge (AM:Low in Fig. 14) is comparable to the highest stiffness values we have estimated for titin (TA:High in Fig. 14). When all 20% of the available crossbridges are attached (AM:High in Fig. 14), the average stiffness of the actin-myosin load path is roughly one order of magnitude stiffer than the highest stiffness values of titin (TA:High in Fig. 14), and two to three orders of magnitude higher than the lowest stiffness titin load path (TP:Low in Fig. 14). Similarly, the maximum XE stiffness and titin stiffness in this work are separated by roughly an order of magnitude: the cat soleus model has a XE stiff-ness of 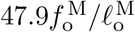 and maximum active titin stiffness of 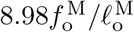 (Table 1); while the rabbit psoas fibril model has a XE stiffness of 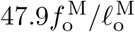 and maximum active titin stiffness of 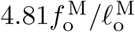 (Appendix H).

**Figure 13.**
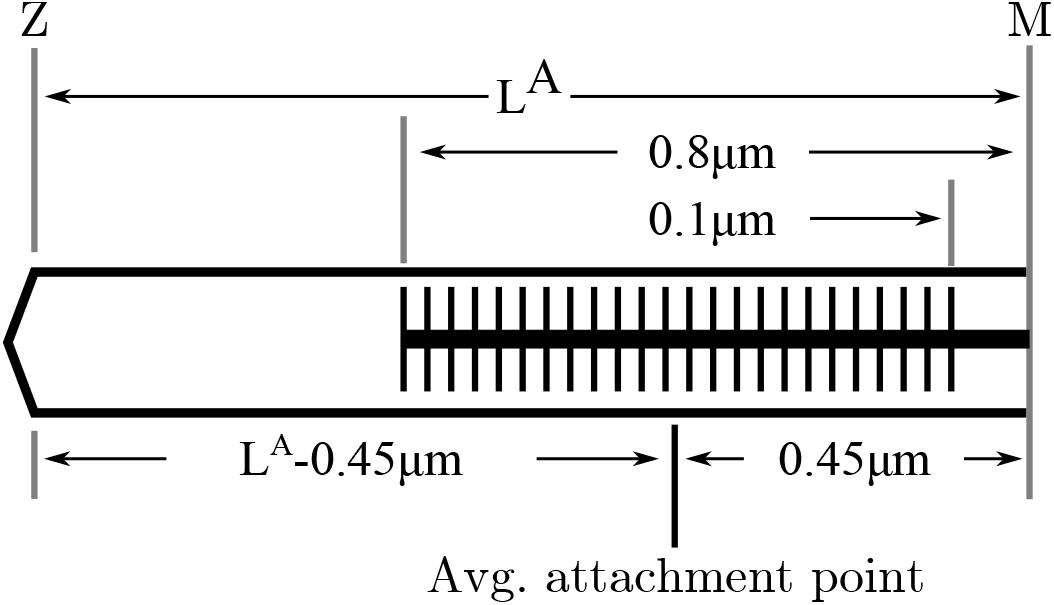
To evaluate the stiffness of the actin-myosin load path, we first determine the average point of attachment. Since the actin filament length varies across species we label it L^A^. Across rabbits, cats and human skeletal muscle myosin geometry is consistent [75]: a half-myosin is 0.8*µm* in length with a 0.1*µm* bare patch in the middle. Thus at full overlap the average point of attachment is 0.45*µm* from the M-line, or L^A^−0.45µm from the Z-line at 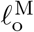. The lumped stiffness of the actin-myosin load path of a half-sarcomere is the stiffness of three springs in series: a spring representing a L^A^ − 0.45µm length of actin, a spring representing the all attached crossbridges, and a spring representing a 0.45*µm* section of myosin.

**Figure 14.**
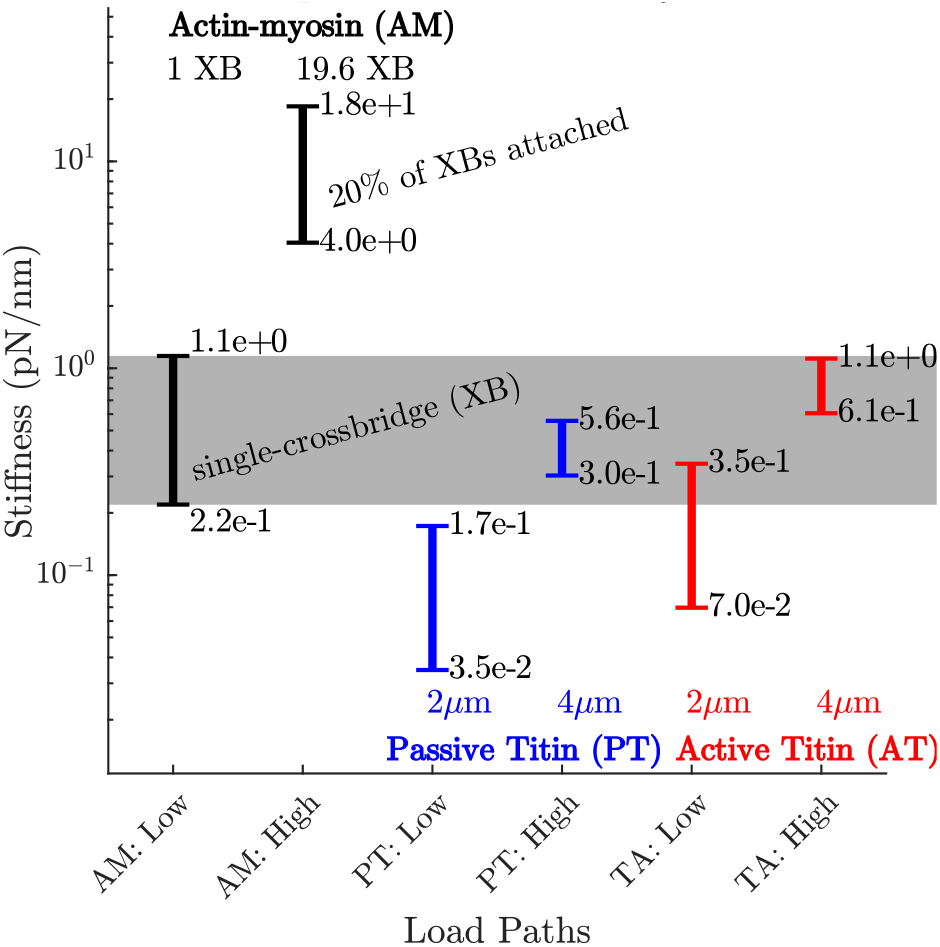
The stiffness of a rabbit’s actin-myosin load path with a single attached crossbridge (1 XB) exceeds the stiff-ness of its titin filament at lengths of 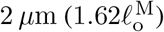 (compare AM:Low to TP:Low and TP:High). Only when titin is stretched to 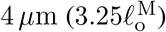 does its stiffness (TP:High and TA:High) become comparable to the actin-myosin with a single attached crossbridge (AM:Low). At higher activations and modest lengths, the stiffness of the actin-myosin load path (AM: High) exceeds the stiffness of titin (TP: Low and TA:Low) by between two and three orders of magnitude. At higher activations and longer lengths, the stiffness of the actin-myosin load path (AM: High) exceeds the stiffness of titin by roughly an order of magnitude (TP:High and TA:High).

**Figure 15.**
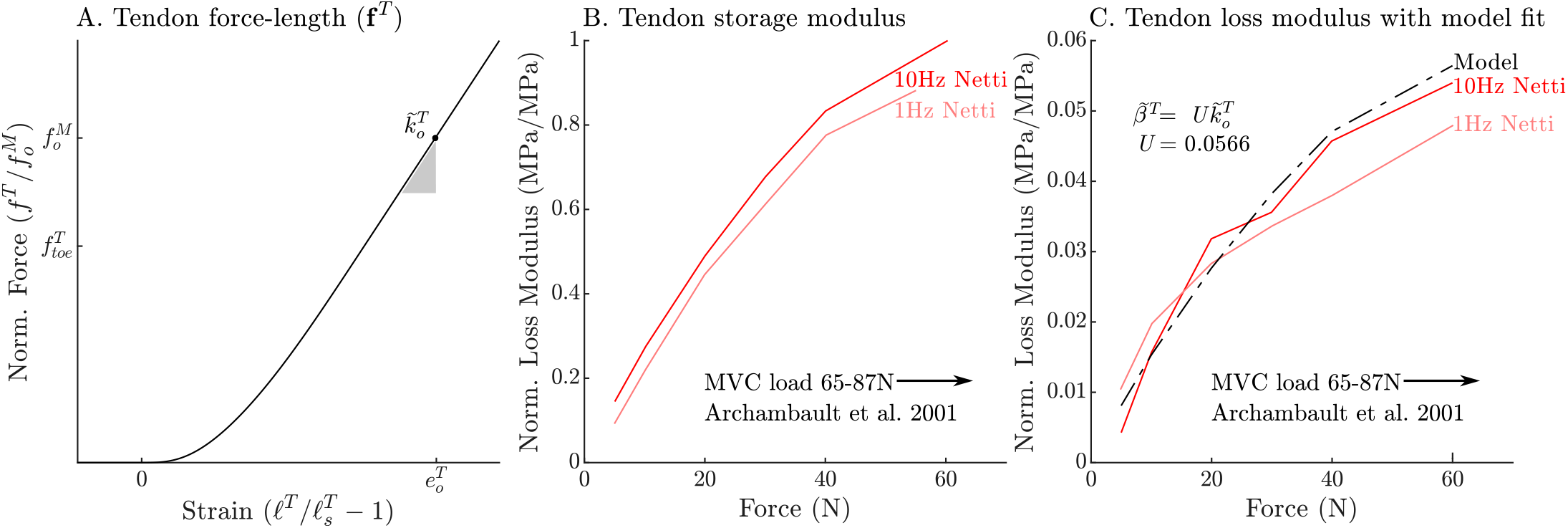
The normalized tendon force-length curve (A) has been been fit to match the cat soleus tendon stiffness measurements of Scott and Loeb [74]. The data of Netti et al. [72] allow us to develop a model of tendon damping as a linear function of tendon stiffness. By normalizing the measurements of Netti et al. [72] by the maximum storage modulus we obtain curves that are equivalent to the normalized stiffness (B) and damping (C) of an Achilles tendon from a rabbit. Both normalized tendon stiffness and damping follow similar curves, but at different scales, allowing us to model tendon damping as a linear function of tendon stiffness (C).

**Figure 16.**
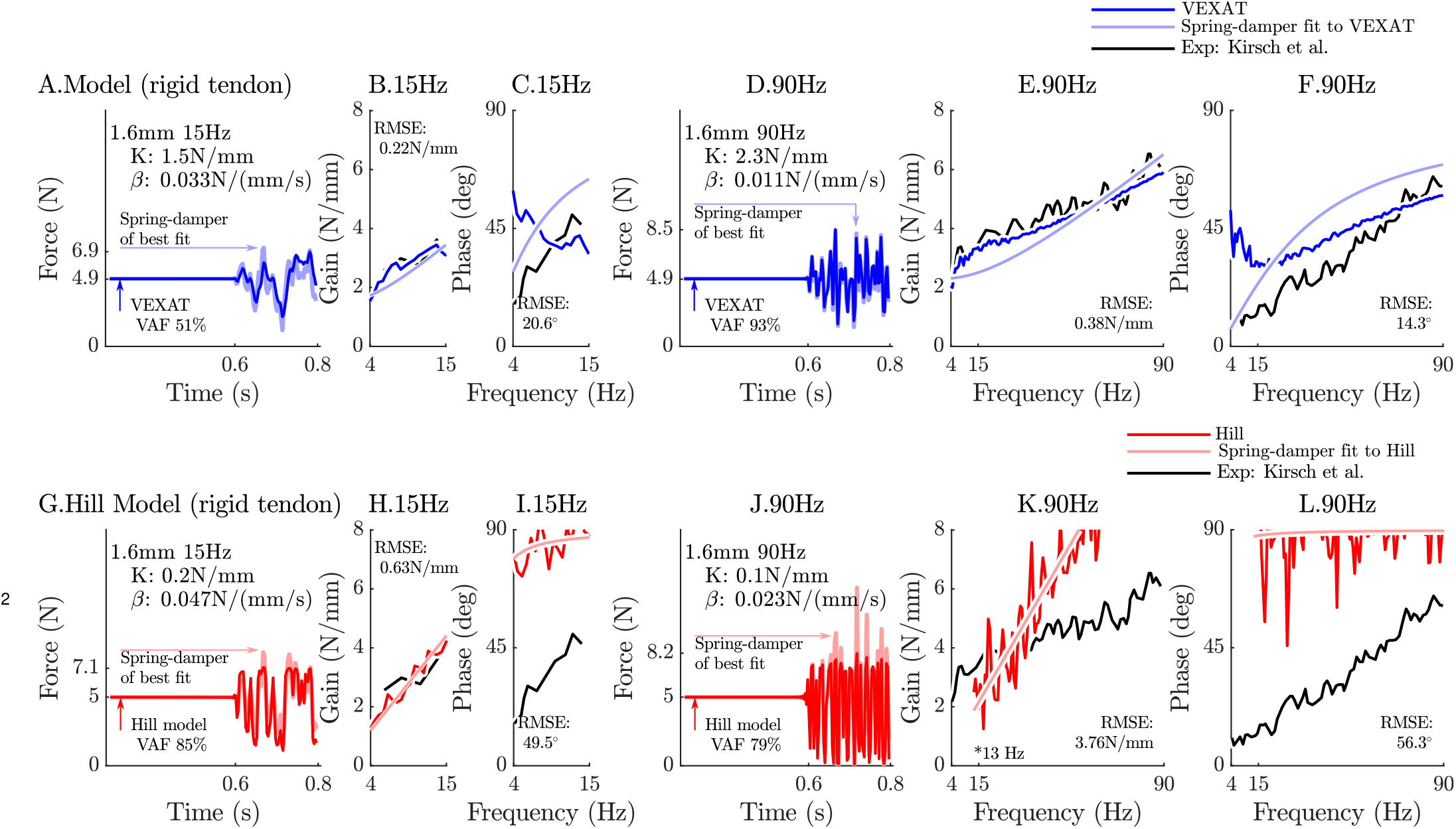
When coupled with a rigid-tendon, the VEXAT model’s VAF (A.), gain response (B.), and phase response (C.) more closely follows the data of Kirsch et al. (Figure 3) [5] than when an elastic-tendon is used. This improvement in accuracy is also observed at the 90 Hz perturbation (D., E., and F.), though the phase response of the model departs from Kirsch et al.’s data [5] for frequencies lower than 30 Hz. Parts of the Hill model’s response to the 15 Hz perturbation are better with a rigid-tendon, with a higher VAF (G.), a lower RMSE gain-response (H.). but have a poor phase-response (I.). In response to the higher frequency perturbations, the Hill model’s response is poor with an elastic (see Fig. 6) or rigid-tendon. The VAF in response to the 90 Hz perturbation remains low (J.), and neither the gain (K.) nor the phase response of the Hill model (L.) follow the data of Kirsch et al. [5]. The rigid-tendon Hill model’s nonlinearity was so strong that the lowest frequency analyzed had to be raised from 4 Hz to 21 Hz to meet the criteria that (*C*_*xy*_)^2^ ≥ 0.67.

## B Model Fitting

Many of the experiments simulated in this work [5], [7] have been performed using cat soleus muscle. While we have been able to extract some architectural parameters directly from the experiments we simulate (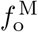 and 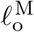 from [7]), we have had to rely on the literature mentioned in Table 1 for the remaining parameters. The remaining properties of the model can be solved by first building a viscoelastic damping model of the tendon; next, by solving for the intrinsic stiffness and damping properties of the CE; and finally, by fitting the passive curves 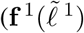 and 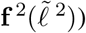 to simultaneously fit the passive force-length curve recorded by Herzog and Leonard [7], using a mixture of tension from titin and the ECM that is consistent with Prado et al.’s data [58], all while maintaining the geometric relationship between **f** ^IgP^ and **f** ^PEVK^ as measured by Trombitás et al. [26].

### B.1 Fitting the tendon’s stiffness and damping

Similar to previous work [17], we model the force-length relation of the tendon using a quintic Bézier spline (Fig. 15A) that begins at 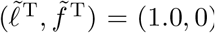 (where 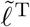 is tendon length normalized by 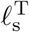, and 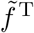 is tension normalized by 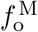), ends at 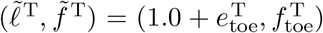 with a nor-malized stiffness of 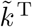, and uses the constants 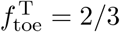 and 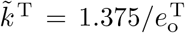 (given 30 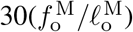 from Scott and Loeb [74], 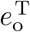 is thus 4.58%). Using the experimental data of Netti et al. [72] we have also constructed a curve to evaluate the damping coefficient of the tendon. The normalized tendon stiffness (termed storage modulus by Netti et al. [72]) and normalized tendon damping (termed loss modulus by Netti et al. [72]) both have a similar shape as the tendon is stretched from slack to 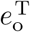 (Fig. 15B and C).

The similarity in shape is likely not a coincidence.

The nonlinear characteristics (Fig. 15) tendon originates from its microscopic structure. Tendon is composed of many fiber bundles with differing resting lengths [72]. Initially the tendon’s fiber bundles begin crimped, but gradually stretch as the tendon lengthens, until finally all fiber bundles are stretched and the tendon achieves its maximum stiffness (Fig. 15B) and damping (Fig. 15C) [72]. Accordingly, in Eqn. 23 we have described the normalized damping of the tendon as being equal to the normalized stiffness of the tendon scaled by a constant *U*. To estimate *U* we have used the measurements of Netti et al. [72] (Fig. 15 B and C) and solved a least-squares problem

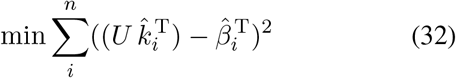

to arrive at a value of *U* = 0.057. The resulting damping model (Fig. 15C) fits the measurements of Netti et al. [72] closely.

### B.2 Fitting the CE’s Impedance

We can now calculate the normalized impedance of the XE using the viscoelastic-tendon model we have constructed and Kirsch et al.’s [5] measurements of the impedance of the entire MTU. Since an MTU is structured with a CE in series with a tendon, the compliance of the MTU is given by

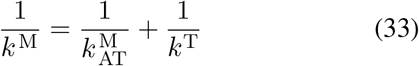

where 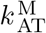 is the stiffness of the CE in the direction of the tendon. We can calculate *k* ^M^ directly by fitting a line to the stiffness vs tension plot that appears in Figure 12 of Kirsch et al. [5] (0.8mm, 0-35 Hz perturbation) and resulting in *k* ^M^ =2.47 N/mm at a nominal force of 5N. Here we use a nominal tension of 5N so that we can later compare our model to the 5N frequency response reported in Figure 3 of Kirsch et al. [5]. Since Kirsch et al. [5] did not report the architectural properties of the experimental specimens, we assume that the architectural properties of the cat used in Kirsch et al.’s experiments are similar to the properties listed in Table 1. We evaluate the stiffness of the tendon model by stretching it until it develops the nominal tension of Kirsch et al.’s Figure 3 data (5N), and then compute its derivative which amounts to *k*^T^ =16.9 N/mm. Finally, using Eqn. 33 we can solve for a value of 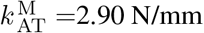. Since the inverse of damping adds for damping elements in series

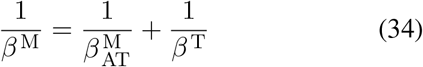

we can use a similar procedure to evaluate 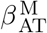, the damping of the CE along the tendon. The value of *β* ^M^ that best fits the damping vs. tension plot that appears in Figure 12 of Kirsch et al. [5] at a nominal tension of 5N is 0.0198 Ns/mm. The tendon damping model we have just constructed develops 0.697 Ns/mm at a nominal load of 5N.

Using Eqn. 34, we arrive at 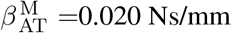 Ns/mm. Due to the pennation model, the stiffness and damping values of the CE differ from those along the tendon.

The stiffness of the CE along the tendon is

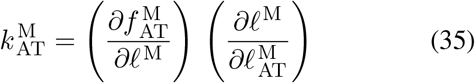

which can be expanded to

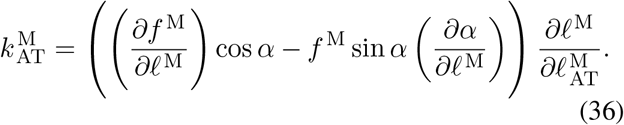

Since we are using a constant thickness pennation model

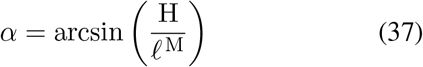

and thus

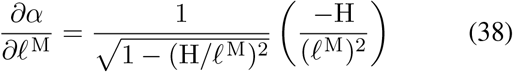

which simplifies to

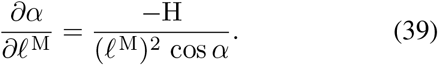

Similarly, the constant thickness pennation model means that

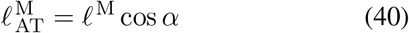

which leads to

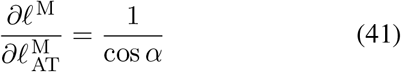

Recognizing that

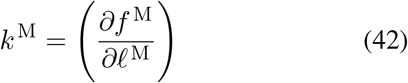

we can solve for *k* ^M^ in terms of 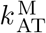 by solving Eqn. 36 for *k* ^M^ and substituting the values of Eqns. 39, and 41. In this case, the values of *k* ^M^ (4.37 N/mm) and 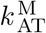 (4.37 N/mm) are the same to three significant figures.

We can use a similar process to transform 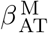 into *β* ^M^ using the pennation model by noting that

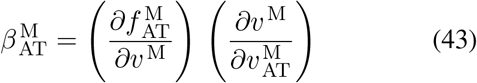

which expands to a much smaller expression

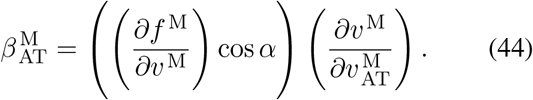

than Eqn. 36 since *α* does not depend on *v* ^M^, and thus *∂α/∂v* ^M^ = 0. By taking a time derivative of Eqn. 40 we arrive at

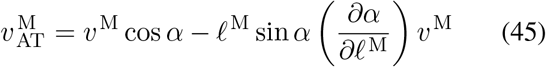

which allows us to solve for

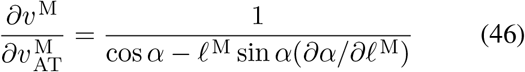

By recognizing that

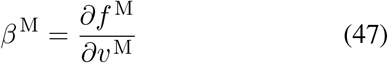

and using Eqns. 44 and 46 we can evaluate *β* ^M^ in terms of 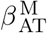. Similar to *k* ^M^, the value of *β* ^M^ (0.020 Ns/mm) is close to 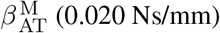. When this same procedure is applied to the stiffness and damping coefficients extracted from the gain and phase profiles from Figure 3 of Kirsch et al. [5], the values of *k* ^M^ and *β* ^M^ differ (4.37 N/mm and 0.0090 Ns/mm) from the results produced using the data of Figure 12 (2.90 N/mm and 0.020 Ns/mm). Likely these differences arise because we have been forced to use a fixed maximum isometric force for all specimens when, in reality, this property varies substantially. We now turn our attention to fitting the titin and ECM elements, since we cannot determine how much of *k* ^M^ and *β* ^M^ are due to the XE until the titin and ECM elements have been fitted.

### B.3 Fitting the force-length curves of titin’s segments

The nonlinear force-length curves used to describe titin (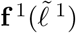 and 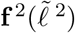 in series), and the ECM 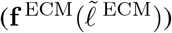 must satisfy three conditions: the total force-length curve produced by the sum of the ECM and titin must match the observed passive-force-length relation[7]; the proportion of titin’s contribution relative to the ECM must be within measured bounds [58]; and finally the stiffness of the 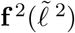 must be a linear scaling of 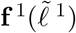 to match the observations of Trombitás et al. [26].

First, we fit the passive force-length curve to the data of Herzog and Leonard [7] to serve as a reference. The curve **f** ^PE^ begins at the normalized length and force coordinates of 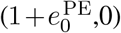 with a slope of 0, ends at 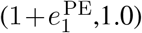 with a slope of 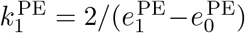, and is linearly extrapolated outside of this region. We solve for the 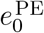 and 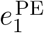 such that

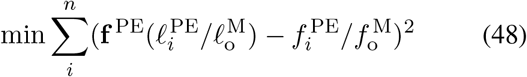

the squared differences between **f** ^PE^ and the passive force-length data of Herzog and Leonard [7] (Fig. 2A shows both the data and the fitted **f** ^PE^ curve) are minimized. While **f** ^PE^ is not used directly in the model, it serves as a useful reference for constructing the ECM and titin force-length curves. We assume that the ECM force-length curve is a linear scaling of **f** ^PE^

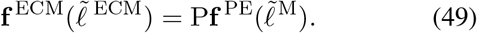

where P is a constant. In this work, we set P to 56% which is the average ECM contribution that Prado et al. [58] measured across 5 different rabbit skeletal muscles^16^. The remaining fraction, 1 − P, of the force-length curve comes from titin.

In mammalian skeletal muscle, titin has three elastic segments [58] connected in series: the proximal Ig segment, the PEVK segment, and the distal Ig segment that is between the PEVK region and the myosin filament (Fig. 1A). Trombitás et al. [26] labelled the PEVK region of titin with antibodies allowing them to measure the distance between the Z-line and the proximal Ig/PEVK boundary (^Z^*ℓ* ^IgP*/*PEVK^), and the distance between the Z-line and the PEVK/distal Ig boundary (^Z^*ℓ* ^PEVK*/*IgD^), while the passive sarcomere was stretched from 2.35 − 4.46µm. By fitting functions to Trombitás et al.’s [26] data we can predict the length of any of titin’s segments under the following assumptions: the T12 segment is rigid (Fig. 1A), the distal Ig segment that overlaps with myosin is rigid (Fig. 1A), and that during passive stretching the tension throughout the titin filament is uniform. Since the sarcomeres in Trombitás et al.’s [26] experiments were passively stretched it is reasonable to assume that tension throughout the free part of the titin filament is uniform because the bond between titin and actin depends on calcium [31], [36] and crossbridge attachment [8].

We begin by digitizing the data of Figure 5 of Trombitás et al. [26] and using the least-squares method to fit lines to ^Z^*ℓ* ^IgP*/*PEVK^ and ^Z^*ℓ* ^PEVK*/*IgD^ (where the superscripts mean ^from^*ℓ*^to^ and so ^Z^*ℓ* ^IgP*/*PEVK^ is the distance from the Z-line to the border of the IgP/PEVK segments). From these lines of best fit we can evaluate the normalized length of the proximal Ig segment

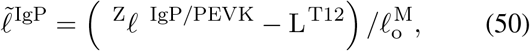

the normalized length of the PEVK segment

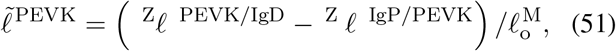

and the normalized length of the distal Ig segment

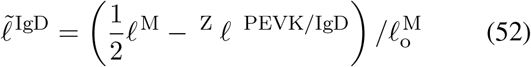

as a function of sarcomere length. Next, we extract the coefficients for linear functions that evaluate the lengths of

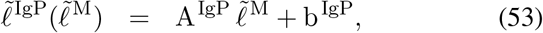

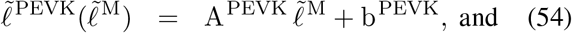

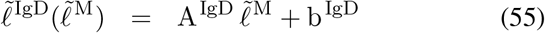

given the 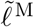. The coefficients that best fit the data from Trombitás et al. [26] appear in Table 2.

**Table 2:** The coefficients of the normalized lengths of 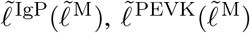, and 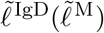 from Eqns. 53-55 under passive lengthening. These coefficients have been extracted from data of Figure 5 of Trombitás et al. [26] using a least-squares fit. Since Figure 5 of Trombitás et al. [26] plots the change in segment length of a single titin filament against the change in length of the entire sarcomere, the resulting slopes are in length normalized units. The slopes sum to 0.5, by construction, to reflect the fact that these three segments of titin stretch at half the rate of the entire sarcomere (assuming symmetry). The cat soleus titin segment coefficients have been formed using a simple scaling of the human soleus titin segment coefficients, and so, are similar. Rabbit psoas titin geometry [58] differs dramatically from human soleus titin [26] and produce a correspondingly large difference in the coefficients that describe the length of the segments of rabbit psoas titin.

| Coefficient | Human<br>Soleus [26] | Feline<br>Soleus | Rabbit<br>Psoas |
| --- | --- | --- | --- |
| $A^{\text{IgP}}$ | 0.177 | 0.177 | 0.262 |
| $b^{\text{IgP}}$ | -0.101 | -0.113 | -0.189 |
| $A^{\text{PEVK}}$ | 0.266 | 0.266 | 0.122 |
| $b^{\text{PEVK}}$ | -0.197 | -0.221 | -0.100 |
| $A^{\text{IgD}}$ | 0.057 | 0.057 | 0.115 |
| $b^{\text{IgD}}$ | -0.033 | -0.033 | -0.083 |

These functions can be scaled to fit a titin filament of a differing geometry. Many of the experiments simulated in this work used cat soleus. Although the lengths of titin filament segments in cat soleus have not been measured, we assume that it is a scaled version of a human soleus titin filament (68 proximal Ig domains, 2174 PEVK residues, and 22 distal Ig domains [26]) since both muscles contain predominately slow-twitch fibers: slow twitch fibers tend to express longer, more compliant titin filaments [58]. Since the optimal sarcomere length in cat skeletal muscle is shorter than in human skeletal muscle (2.43 *µ*m vs. 2.73 *µ*m, [75]) the coefficients for Eqns. 53-55 differ slightly (see the feline soleus column in Table 2). In addition, by assuming that the titin filament of cat skeletal muscle is a scaled version of the titin filament found in human skeletal muscle, we have implicitly assumed that the cat’s skeletal muscle titin filament has 60.5 proximal Ig domains, 1934.7 PEVK residues, and 19.6 distal Ig domains. Although a fraction of a domain does not make physical sense, we have not rounded to the nearest domain and residue to preserve the sarcomere length-based scaling.

In contrast, the rabbit psoas fibril used in the simulation of Leonard et al. [8] has a known titin geometry (50 proximal Ig domains, 800 PEVK residues, and 22 distal Ig domains [58]) which differs substantially from the isoform of titin expressed in the human soleus. To create the rabbit psoas titin length functions 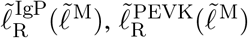, and 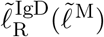 we begin by scaling the human soleus PEVK length function 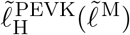 by the relative proportion of PEVK residues of 800*/*2174. The length of the two Ig segments

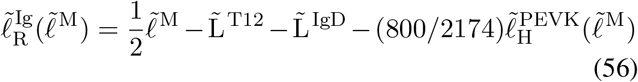

is what remains from the half-sarcomere once the rigid lengths of titin (0.100 *µ*m for L^T12^ and 0.8150 *µ*m for L^IgD^ [60]) and the PEVK segment length have been subtracted away. The function that describes 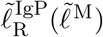 and 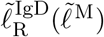 can then be formed by scaling 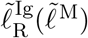 by the proportion of Ig domains in each segment

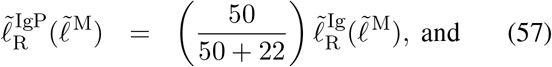

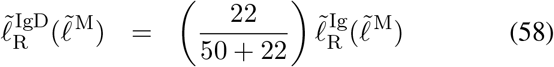

which produce the coefficients that appear in the rabbit psoas column in Table 2. While we have applied this approach to extend Trombitás et al.’s [26] results to a rabbit psoas, in principle this approach can be applied to any isoform of titin provided that its geometry is known, and the Ig domains and PEVK residues in the target titin behave similarly to those in human soleus titin.

The only detail that remains is to establish the shape of the IgP, PEVK, and IgD force-length curves. Studies of individual titin filaments, and of its segments, make it clear that titin is mechanically complex. For example, the tandem Ig segments (the IgD and IgP segments) are composed of many folded domains (titin from human soleus has two Ig segments that together have nearly 100 domains [26]). Each domain appears to be a simple nonlinear spring until it unfolds and elongates by nearly 25 nm in the process [88]. Unfolding events appear to happen individually during lengthening experiments [88], with each unfolding event occurring at a slightly greater tension than the last, giving an Ig segment a force-length curve that is saw-toothed. Although detailed models of titin exist that can simulate the folding and unfolding of individual Ig domains, this level of detail comes at a cost of a state for each Ig domain which can add up to nearly a hundred extra states [40] in total.

Active and passive lengthening experiments at the sarcomere-level hide the complexity that is apparent when studying individual titin filaments. The experiments of Leonard et al. [8] show that the sarcomeres in a filament (from rabbit psoas) mechanically fail when stretched passively to an average length of 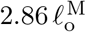, but can reach 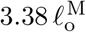 when actively lengthened. Leonard et al. [8] showed that titin was the filament bearing these large forces since the sarcomeres were incapable of developing active or passive tension when the experiment was repeated after the titin filaments were chemically cut. It is worth noting that the forces measured by Leonard et al. [8] contain none of the complex saw-tooth pattern indicative of unfolding events even though 72 of these events would occur as each proximal and distal Ig domain fully unfolded and reached its maximal length^17^. Although we cannot be sure how many unfolding events occurred during Leonard et al.’s experiments [8], due to sarcomere non-homogeneity [89], it is likely that many Ig unfolding events occurred since the average sarcomere length at failure 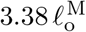 was longer than the maximum length of 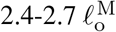 that would be predicted from the geometry of rabbit psoas titin^18^.

Since we are interested in a computationally efficient model that is accurate at the whole muscle level, we model titin as a multi-segmented nonlinear spring but omit the states needed to simulate the folding and unfolding of Ig domains. Simulations of active lengthening using our titin model will exhibit the enhanced force development that appears in experiments [7], [8], but will lack the nonlinear saw-tooth force-length profile that is measured when individual titin filaments are lengthened [88]. To have the broadest possible application, we will fit titin’s force-length curves to provide reasonable results for both moderate [7] and large active stretches [8]. Depending on the application, it may be more appropriate to use a stiffer force-length curve for the Ig segment if normalized sarcomere lengths stays within 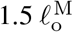 and no unfolding events occur as was done by Trombitás et al. [65].

To ensure that the serially connected force-length curves of 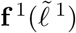 and 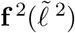 closely reproduce 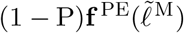, we are going to use affine transformations of **f** ^PE^ to describe 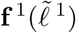 and 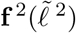. The total stiffness of the half-sarcomere titin model is given by

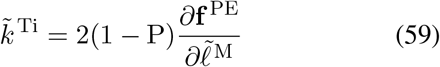

which is formed by the series connection of 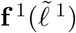 and 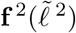

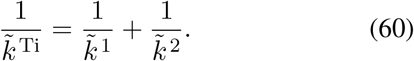

Since each of titin’s segments is exposed to the same tension in Trombitás et al.’s experiment [26] the slopes of the lines that Eqns. 53-55 describe are directly proportional to the relative compliance (inverse of stiffness) between of each of titin’s segments. Using this fact, we can define the normalized stretch rates of the proximal titin segment

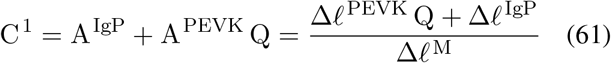

and the distal titin segment

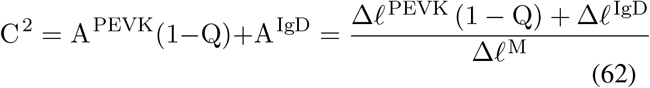

which are proportional to the compliance of two titin segments in our model. When both the 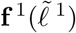 and 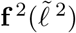 curves are beyond the toe region the stiffness is a constant and is given by

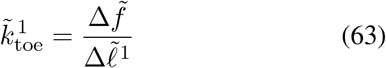

and

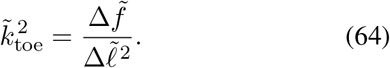

Dividing Eqn. 63 by 64 eliminates the unknown 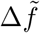 and results in an expression that relates the ratio of the terminal linear stiffness of 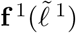 and 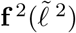

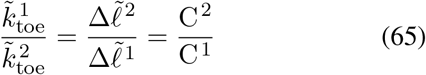

to the relative changes in Eqns. 61 and 62. Substituting Eqns. 65, and 60 into Eqn. 59 yields the expression

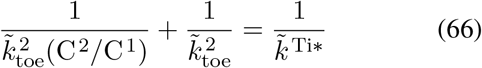

which can be simplified to

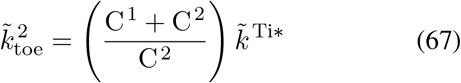

and this expression can be evaluated using the terminal stiffness of titin 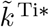 and the coefficients listed in Table 2. Substituting Eqn. 67 into Eqn. 65 yields

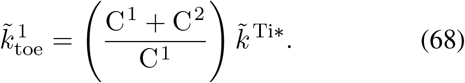

The curves 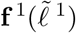 and 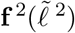 can now be formed by scaling and shifting the total force-length curve of titin (1 −P)**f** ^PE^. By construction, titin’s force-length curve develops a tension of (1 − P), and has reached its terminal stiffness, when the CE reaches a length 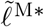 such that 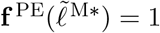. Using Eqns. 53-55 and the appropriate coefficients in Table 2 we can evaluate the normalized length developed by the *ℓ* ^1^ segment

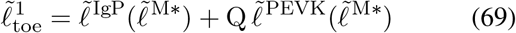

and *ℓ* ^2^ segment

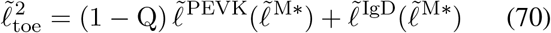

at a CE length of 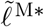. The 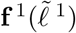 curve is formed by shifting and scaling the (1−P)**f** ^PE^ curve so that it develops a normalized tension of (1 − P) and a stiffness of 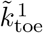 a length of 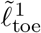. Similarly, the 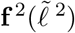 curve is made by shifting and scaling the (1 − P)**f** ^PE^ curve to develop a normalized tension of (1 − P) and a stiffness of 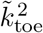 at a length of 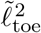.

By construction, the spring network formed by the 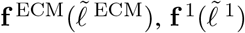, and 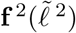 curves follows the fitted **f** ^PE^ curve (Fig. 3A) such that the ECM curve makes up 56% of the contribution. When the CE is active and 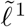 is effectively fixed in place, the distal segment of titin contributes higher forces since 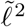 undergoes higher strains (Fig. 3A). Finally, when the experiment of Trombitás et al. [26] are simulated the movements of the IgP/PEVK and PEVK/IgD boundaries in the titin model closely follow the data (Fig. 3C).

The process we have used to fit the ECM and titin’s segments makes use of data within modest normalized CE lengths (2.35-4.46*µ*m, or 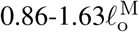 [26]). Scenarios in which the CE reaches extremely long lengths, such as during injury or during Leonard et al.’s experiment [8], require fitting titin’s force-length curve beyond the typical ranges observed in-vivo. The WLC model has been used successfully to model the force-length relation of individual titin segments [65] at extreme lengths. In this work, we consider two different extensions to 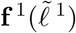 and 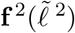 : a linear extrapolation, and the WLC model. Since the fitted **f** ^PE^ curve is linearly extrapolated, so too are the 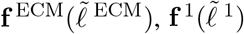, and 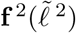 curves by default. Applying the WLC to our titin curves requires a bit more effort.

We have modified the WLC to include a slack length 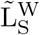 (the superscript W means WLC) so that the WLC model can made to be continuous with 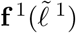 and 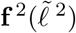. The normalized force developed by our WLC model is given by

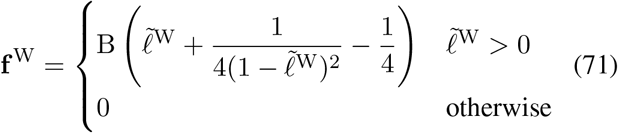

where B is a scaling factor and the normalized segment length 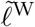 is defined as

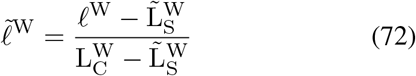

where 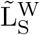 is the slack length, and 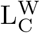 is the contour length of the segment. To extend the 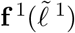 curve to follow the WLC model, we first note the normalized contour length of the *ℓ* ^1^ segment

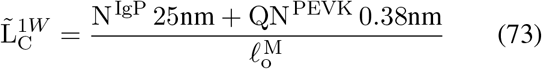

by counting the number of proximal Ig domains (N ^IgP^), at the number of PEVK residues (QN ^PEVK^) associated with *ℓ* ^1^ and by scaling each by the maximum contour length of each Ig domain (25nm [88]), and each PEVK residue (between 0.32 [65] and 0.38 nm [90] see pg. 254). This contour length defines the maximum length of the segment, when all of the Ig domains and PEVK residues have been unfolded. Similarly, the contour length of 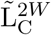 is given by

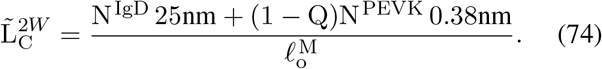

Next, we define the slack length by linearly extrapolating backwards from the final fitted force (1 − P)

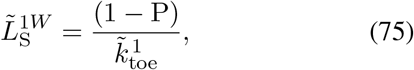

and similarly

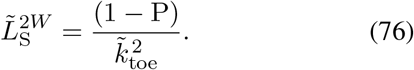

We can now solve for B in Eqn. 71 so that 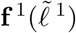 and 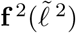 are continuous with each respective WLC extrapolation. However, we do not use the WLC model directly because it contains a numerical singularity which is problematic during numerical simulation. Instead, we add an additional Bézier segment to fit the WLC extension that spans between forces of (1 − P) and twice the normalized failure force 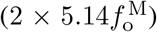 noted by Leonard et al. [8]. To fit the shape of the final Bézier segment, we adjust the locations of the internal control points to minimize the _W_ squared differences between the modified WLC model and ^S^ the final Bézier curve (Fig. 10A). The final result is a set of curves 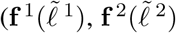, and 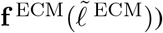 which, between forces 0 and (1 − P), will reproduce **f** ^PE^, Trombitás et al.’s measurements [26], and do so with a reasonable titin-ECM balance [58]. For forces beyond (1 − P), the curve will follow the segment-specific WLC model up to twice the expected failure tension noted by Leonard et al. [8].

### B.4 Fitting the XE’s Impedance

With the passive curves established, we can return to the problem of identifying the normalized maximum stiffness 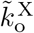 and damping 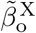 of the lumped XE element. Just prior to discussing titin, we had evaluated the impedance of the cat soleus CE in Kirsch et al.’s [5] Figure 12 to be *k* ^M^ =2.90 N/mm and *β* ^M^ =0.020 Ns/mm at a nominal active tension of 5N. The normalized stiffness *k* ^M^ can be found by taking the partial derivative of Eqn. 15 with respect to 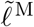

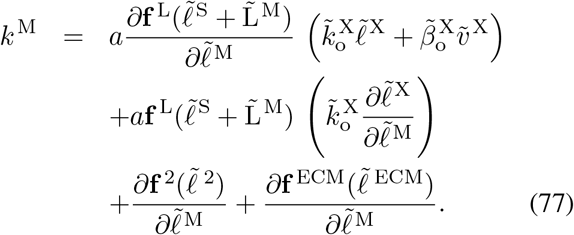

By noting that all of our chosen state variables in Eqn. 13 are independent and by making use of the kinematic relationships in Eqns. 9 and 10 we can reduce Eqn. 77 to

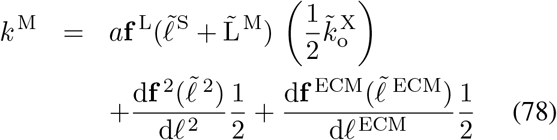

and solve for 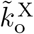

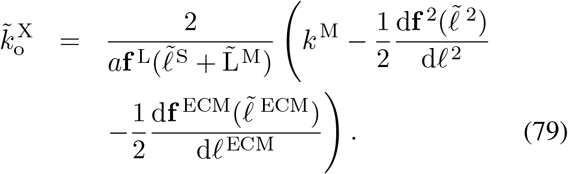

When using to the data from Figure 12 in Kirsch et al. [5], we end up with 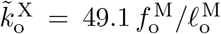 for the elastictendon model, and 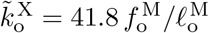 for the rigid-tendon model. When this procedure is repeated for Figure 3 of Kirsch et al. [5] (from a different specimen) we are left with 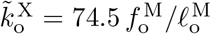 for the elastic-tendon model and 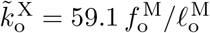 for the rigid-tendon model. The value for 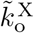 is much larger than *k* ^M^ because the *a* needed to generate 5N is only 0.231. Similarly, we can form the expression for the normalized damping of the CE by taking the partial derivative of Eqn. 15 with respect to 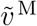

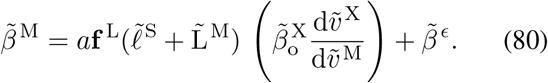

As with *k* ^M^, the expression for 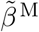 can be reduced to

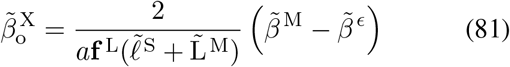

which evaluates to 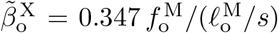 for both the elastic and rigid-tendon models using Kirsch et al.’s [5] Figure 12 data. The damping coefficients of the elastic and rigid-tendon models is similar because the damping coefficient of the musculotendon is dominated by the damping coefficient of CE. When the data from Kirsch et al.’s [5] Figure 3 is used, the damping coefficients of the elastic and rigid-tendon models are 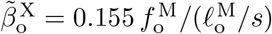 and 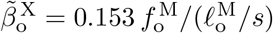 respectively.

The dimensionless parameters 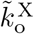 and 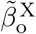 can be used to approximate the properties of other MTUs given 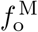 and 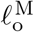. The stiffness and damping of the lumped cross-bridge element will scale linearly with 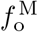 and inversely with 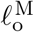 provided the impedance properties of individual crossbridges, and the maximum number of crossbridges attached per sarcomere, is similar between a feline’s skeletal muscle sarcomeres and those of the target MTU. This approximation is rough, however, since the values for 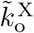 and 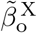 (Table 3) have a relative error of 41% and 76% when evaluated using Kirsch et al.’s [5] Figure 3 and Figure 12. In addition, when simulated, the stiffness and damping of the LTI system of best fit may differ from 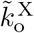 and 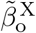 at low frequencies because the movement of the attachment point has been ignored in Eqns. 79 and 81. This approximation explains why the VEXAT’s stiffness profile (Fig. 7 A. and C) is below Kirsch et al.’s [5] data, despite having used this data to fit the *k* ^M^ and 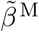 terms in Eqns. 79 and 81. The accuracy of this approximation, however, improves at higher frequencies (Fig. 6 E and F) because the attachment point’s movements become increasingly limited due to the time constant *τ* ^S^ in Eqn. 16. Unfortunately this is a trade-off due to the formulation of Eqn. 16: the VEXAT mode can fit Kirsch et al.’s [5] data at low frequencies, or high frequencies, but not both simultaneously.

**Table 3:** Normalized titin and crossbridge parameters fit to data from the literature.

| Symbol | Value | Unit | Source |
| --- | --- | --- | --- |
| $\tilde{k}^{\text{Ti}}$ | 3.88 | $f_o^M / \ell_o^M$ | [7] [58] |
| $\tilde{k}^1$ | 5.17 | $f_o^M / \ell_o^M$ | [26] |
| $\tilde{k}^2$ | 8.42 | $f_o^M / \ell_o^M$ | [26] |
| $\tilde{k}_o^X$ | 74.5 | $f_o^M / (\ell_o^M)$ | [5] (Fig. 3) |
| $\tilde{\beta}_o^X$ | 0.155 | $f_o^M / (\ell_o^M / s)$ | [5] (Fig. 3) |
| $\tilde{k}_o^X$ | 49.1 | $f_o^M / (\ell_o^M)$ | [5] (Fig. 12) |
| $\tilde{\beta}_o^X$ | 0.347 | $f_o^M / (\ell_o^M / s)$ | [5] (Fig. 12) |

**Table 4:** Mean normalized stiffness coefficients (A.), mean normalized damping coefficients (B.), VAF (C.), and the bandwidth (D.) of linearity (coherence squared *>* 0.67) for models with elastic-tendons. Here the proposed model has been fitted to Figure 12 of Kirsch et al. [5], while the experimental data from Kirsch et al. [5] comes from Figures 9 and 10. Experimental data from Figure 12 from Kirsch et al. has not been included in this table because it would only contribute 1 entry and would overwrite values from Figures 9 and 10. The impedance experiments at each combination of perturbation amplitude and frequency have been evaluated at 10 different nominal forces linearly spaced between 2.5N and 11.5N. The results presented in the table are the mean values of these ten simulations. The VAF is evaluated between the model and the spring-damper of best fit, rather than to the response of biological muscle (which was not published by Kirsch et al. [5]). Finally, model values for the VAF (C.) and the bandwidth of linearity (D.) that are worse than those published by Kirsch et al. [5] appear in bold font.

| A. Norm. Stiffness ($\frac{K}{F}$) | Kirsch et al. | | | Model | | | Hill | | |
| --- | --- | --- | --- | --- | --- | --- | --- | --- | --- |
|  | 15Hz | 35Hz | 90Hz | 15Hz | 35Hz | 90Hz | 15Hz | 35Hz | 90Hz |
| 0.4 mm | 0.56 | 0.85 | 0.87 | 0.33 | 0.32 | 0.28 | 0.45 | 1.02 | 1.68 |
| 0.8 mm | 0.46 |  |  | 0.30 | 0.30 | 0.24 | 0.30 | 0.66 | 1.37 |
| 1.6 mm | 0.28 | 0.38 | 0.50 | 0.22 | 0.23 | 0.19 | 0.18 | 0.36 | 0.96 |
| B. Norm. damping $\frac{\beta}{F}$ | 15Hz | 35Hz | 90Hz | 15Hz | 35Hz | 90Hz | 15Hz | 35Hz | 90Hz |
|  | 15Hz | 35Hz | 90Hz | 15Hz | 35Hz | 90Hz | 15Hz | 35Hz | 90Hz |
| 0.4 mm | 0.0118 | 0.0049 | 0.0038 | 0.0059 | 0.0044 | 0.0039 | 0.0196 | 0.0105 | 0.0029 |
| 0.8 mm | 0.0118 | 0.0049 | 0.0038 | 0.0060 | 0.0044 | 0.0039 | 0.0157 | 0.0098 | 0.0033 |
| 1.6 mm | 0.0118 | 0.0049 | 0.0038 | 0.0062 | 0.0045 | 0.0039 | 0.0112 | 0.0079 | 0.0029 |
| C. VAF (%) | 15Hz | 35Hz | 90Hz | 15Hz | 35Hz | 90Hz | 15Hz | 35Hz | 90Hz |
|  | 15Hz | 35Hz | 90Hz | 15Hz | 35Hz | 90Hz | 15Hz | 35Hz | 90Hz |
| 0.4 mm |  |  |  | **64** | **74** | 86 | **28** | **44** | **75** |
| 0.8 mm |  | 78-99% |  | **74** | 85 | 94 | **37** | **38** | **68** |
| 1.6 mm |  |  |  | **76** | 91 | 97 | **48** | **35** | **62** |
| D. Bandwidth (Hz) s.t.<br>Coherence<sup>2</sup> $> 0.67$ | 15Hz | 35Hz | 90Hz | 15Hz | 35Hz | 90Hz | 15Hz | 35Hz | 90Hz |
|  | 15Hz | 35Hz | 90Hz | 15Hz | 35Hz | 90Hz | 15Hz | 35Hz | 90Hz |
| 0.4 mm | 4-15 | 4-35 | 4-90 | 4-15 | 4-35 | 4-90 | 4-15 | 4-35 | 4-90 |
| 0.8 mm | 4-15 | 4-35 | 4-90 | 4-15 | 4-35 | 4-90 | 4-15 | 4-35 | **7-90** |
| 1.6 mm | 4-15 | 4-35 | 4-90 | 4-15 | 4-35 | 4-90 | 4-15 | 4-35 | **7-90** |

**Table 5:** Mean normalized stiffness coefficients (A.), mean normalized damping coefficients (B.), VAF (C.), and the bandwidth (D.) of linearity (coherence squared *>* 0.67) for models with rigid tendons. All additional details are identical to those of Table except the tendon of the model is rigid.

| A. Norm. Stiffness ($\frac{K}{F}$) | Kirsch et al. | | | Model | | | Hill | | |
| --- | --- | --- | --- | --- | --- | --- | --- | --- | --- |
|  | 15Hz | 35Hz | 90Hz | 15Hz | 35Hz | 90Hz | 15Hz | 35Hz | 90Hz |
| 0.4 mm | 0.56 | 0.85 | 0.87 | 0.32 | 0.31 | 0.28 | 0.05 | 0.01 | 0.01 |
| 0.8 mm | 0.46 |  |  | 0.30 | 0.29 | 0.25 | 0.05 | 0.00 | 0.01 |
| 1.6 mm | 0.28 | 0.38 | 0.50 | 0.23 | 0.23 | 0.20 | 0.04 | 0.00 | 0.02 |
| B. Norm. damping $\frac{\beta}{F}$ | 15Hz | 35Hz | 90Hz | 15Hz | 35Hz | 90Hz | 15Hz | 35Hz | 90Hz |
|  | 15Hz | 35Hz | 90Hz | 15Hz | 35Hz | 90Hz | 15Hz | 35Hz | 90Hz |
| 0.4 mm | 0.0118 | 0.0049 | 0.0038 | 0.0054 | 0.0042 | 0.0038 | 0.0217 | 0.0172 | 0.0125 |
| 0.8 mm | 0.0118 | 0.0049 | 0.0038 | 0.0055 | 0.0042 | 0.0038 | 0.0148 | 0.0111 | 0.0078 |
| 1.6 mm | 0.0118 | 0.0049 | 0.0038 | 0.0057 | 0.0043 | 0.0039 | 0.0094 | 0.0068 | 0.0046 |
| C. VAF (%) | 15Hz | 35Hz | 90Hz | 15Hz | 35Hz | 90Hz | 15Hz | 35Hz | 90Hz |
|  | 15Hz | 35Hz | 90Hz | 15Hz | 35Hz | 90Hz | 15Hz | 35Hz | 90Hz |
| 0.4 mm |  |  |  | 86 | 96 | 99 | 95 | 92 | 88 |
| 0.8 mm |  | 78-99% |  | 85 | 96 | 99 | 89 | 86 | 83 |
| 1.6 mm |  |  |  | 82 | 95 | 99 | 85 | 82 | 79 |
| D. Bandwidth (Hz) s.t.<br>Coherence<sup>2</sup> $> 0.67$ | 15Hz | 35Hz | 90Hz | 15Hz | 35Hz | 90Hz | 15Hz | 35Hz | 90Hz |
|  | 15Hz | 35Hz | 90Hz | 15Hz | 35Hz | 90Hz | 15Hz | 35Hz | 90Hz |
| 0.4 mm | 4-15 | 4-35 | 4-90 | 4-15 | 4-35 | 4-90 | 4-15 | 4-35 | **13-90** |
| 0.8 mm | 4-15 | 4-35 | 4-90 | 4-15 | 4-35 | 4-90 | 4-15 | 4-35 | **13-90** |
| 1.6 mm | 4-15 | 4-35 | 4-90 | 4-15 | 4-35 | 4-90 | 4-15 | 4-35 | **13-90** |

**Table 6:** The VEXAT and Hill model’s fitted rabbit psoas fibril MTU parameters. As in Table 6, parameters shared by the VEXAT and Hill model are highlighted in grey. Short forms are used to save space: length ‘len’, velocity ‘vel’, acceleration ‘acc’, half ‘h’, activation ‘act’, segment ‘seg’, threshold ‘thr’, and stiffness ‘stiff’. The letter preceding a reference indicates the experimental animal:’C’ for cat, ‘H’ for human, while nothing at all is rabbit skeletal muscle. Letters following a reference indicate how the data was used to evaluate the parameter: ‘A’ for arbitrary for simulating Leonard et al. [8], ‘n/a’ for a parameter that is not applicable to a fibril model, ‘—’ value taken from the cat soleus MTU, ‘C’ calculated, ‘F’ fit, ‘E’ estimated, ‘S’ scaled, and ‘D’ for default if a default value from another model was used. Only parameters that do not affect the outcome of our simulation of Leonard et al. [8] are marked ‘A’. Clearly the parameters that appear in this Table do not represent a generic rabbit psoas fibril model, but instead a rabbit psoas fibril model that is sufficient to simulate the experiment of Leonard et al. [8]. Finally, values for N ^IgP^, N ^PEVK^, and N ^IgD^ were obtained by taking a 70% and 30% average of the values for 3300 kD and 3400 kD titin to match the composition of rabbit psoas titin as closely as possible.

| Parameter |  | Value | Source |
| --- | --- | --- | --- |
| A. Basic parameters |  |  |  |
| Max iso force | $f_o^M$ | 1 N | A |
| Opt CE len | $\ell_o^M$ | 1 mm | A |
| Pen angle | $\alpha$ | 0° | A |
| Act time const | $\tau_A$ | 10 ms | A,H[17]D |
| De-act time const | $\tau_D$ | 40 ms | A,H[17]D |
| B. Force-velocity relation: $\mathbf{f}^V(\hat{v}^M)$ | | | |
| Max shortening vel | $v_{\max}^M$ | $4.5 \frac{\ell_o^M}{s}$ | H[17]D |
| $\mathbf{f}^V$ at $-\frac{1}{2}v_{\max}^M$ | $\tilde{f}_1^V$ | $0.1 f_o^M$ | H[17]D |
| $\mathbf{f}^V$ at $\hat{v}^M = +0$ | $\tilde{f}_2^V$ | $1.3 f_o^M$ | H[17]D |
| $\mathbf{f}^V$ at $v_{\max}^M$ | $\tilde{f}_3^V$ | $1.45 f_o^M$ | H[17]D |
| $v_{\max}^M$ scaling | $s^V$ | 0.950 | — |
| C. Active force-length relation: $\mathbf{f}^L(\tilde{\ell}^M)$ | | | |
| Opt sarcomere len | $\tilde{L}_o^M$ | $2.46 \mu\text{m}$ | [60] |
| Actin len | $\tilde{L}^A$ | $0.455 \ell_o^M$ | [60] |
| Myosin h-len | $\tilde{L}^M$ | $0.331 \ell_o^M$ | [60] |
| Myosin bare h-len | $\tilde{L}^B$ | $0.0163 \ell_o^M$ | [60] |
| Offset | $\Delta^L$ | $-\frac{2}{k_o^X} \ell_o^M$ | C |
| D. Passive force-length relation: $\mathbf{f}^{\text{PE}}(\tilde{\ell}^M)$ | | | |
| Slack len | $\tilde{\ell}_s^{\text{PE}}$ | $1 \ell_o^M$ | [8]F |
| Toe len | $\tilde{\ell}_{\text{toe}}^{\text{PE}}$ | $1.71 \ell_o^M$ | [8]F |
| Toe force | $\tilde{f}_{\text{toe}}^{\text{PE}}$ | $0.31 f_o^M$ | [8]F |
| Toe stiffness | $\tilde{k}_{\text{toe}}^{\text{PE}}$ | $0.870 \frac{f_o^M}{\ell_o^M}$ | [8]F |
| E. Compressive force-length relation: $\mathbf{f}^{\text{KE}}(\tilde{\ell}^M)$ | | | |
| Slack len | $\tilde{\ell}_s^{\text{PE}}$ | $\frac{1}{10} \ell_o^M$ | E |
| Toe len | $\tilde{\ell}_{\text{toe}}^{\text{PE}}$ | $0.00 \ell_o^M$ | E |
| Toe force | $\tilde{f}_{\text{toe}}^{\text{PE}}$ | $1.00 f_o^M$ | E |

| F. XE viscoelastic model |  |  |  |
| --- | --- | --- | --- |
| Stiffness | $\tilde{k}_o^X$ | $49.1 \frac{f_o^M}{\ell_o^M}$ | — |
| Damping | $\tilde{\beta}_o^X$ | $0.347 \frac{f_o^M}{\ell_o^M/s}$ | — |
| Acc. time const | $\tau^S$ | $1.00\text{e-}3 \text{ s}$ | — |
| Num acc damping | D | 1.00 | — |
| Low act threshold | $a_L$ | 0.0500 | — |
| Len tracking gain | $G_L$ | $1000 \frac{1}{s}$ | — |
| Vel tracking gain | $G_V$ | 1000 | — |
| G. Titin & ECM Parameters |  |  |  |
| ECM fraction | P | 0 | E |
| PEVK attach pt | Q | 0.675 | [8]F |
| Z-line-T12 len | $\tilde{L}^{\text{T12}}$ | $0.0407 \ell_o^M$ | H[65] |
| IgD rigid h-len | $\tilde{L}^{\text{IgD}}$ | $\tilde{L}^M$ | [60] |
| No IgP domains | $N^{\text{IgP}}$ | 45.1 | [58]C |
| No PEVK residues | $N^{\text{PEVK}}$ | 695 | [58]C |
| No IgD domains | $N^{\text{IgD}}$ | 22 | [58]C |
| Active damping | $\beta_A^{\text{PEVK}}$ | $975 \frac{f_o^M}{\ell_o^M}$ | [8]F |
| Passive damping | $\beta_P^{\text{PEVK}}$ | $0.1 \frac{f_o^M}{\ell_o^M}$ | — |
| Length threshold | $\tilde{\ell}_s^M$ | $\frac{1}{2} \tilde{\ell}_s^{\text{PE}}$ | — |
| Act threshold | $A_o$ | 0.05 | — |
| Step transition | R | 0.01 | — |
| H. Titin's force-length relations: $\mathbf{f}^1(\tilde{\ell}^1)$ & $\mathbf{f}^2(\tilde{\ell}^2)$ | | | |
| $\mathbf{f}^1(\tilde{\ell}^1)$ slack len | $\tilde{\ell}_s^1$ | $0.137 \ell_o^M$ | H[65]S,[8]F |
| $\mathbf{f}^1(\tilde{\ell}^1)$ toe len | $\tilde{\ell}_{\text{toe}}^1$ | $0.264 \ell_o^M$ | H[65]S,[8]F |
| $\mathbf{f}^1(\tilde{\ell}^1)$ toe force | $\tilde{f}_{\text{toe}}^1$ | $0.163 f_o^M$ | H[65]S,[8]F |
| $\mathbf{f}^1(\tilde{\ell}^1)$ toe stiff | $\tilde{k}_{\text{toe}}^1$ | $2.55 \frac{f_o^M}{\ell_o^M}$ | H[65]S,[8]F |
| $\mathbf{f}^2(\tilde{\ell}^2)$ slack len | $\tilde{\ell}_s^2$ | $0.067 \ell_o^M$ | H[65]S,[8]F |
| $\mathbf{f}^2(\tilde{\ell}^2)$ toe len | $\tilde{\ell}_{\text{toe}}^2$ | $0.129 \ell_o^M$ | H[65]S,[8]F |
| $\mathbf{f}^2(\tilde{\ell}^2)$ toe force | $\tilde{f}_{\text{toe}}^2$ | $0.163 f_o^M$ | H[65]S,[8]F |
| $\mathbf{f}^2(\tilde{\ell}^2)$ toe stiff | $\tilde{k}_{\text{toe}}^2$ | $5.25 \frac{f_o^M}{\ell_o^M}$ | H[65]S,[8]F |

## C Model Initialization

Solving for an initial state is challenging since we are given *a, ℓ* ^P^, and *v* ^P^ and must solve for *v* ^S^, *ℓ* ^S^, and *ℓ* ^1^ for a rigid-tendon model, and additionally *ℓ* ^M^ if an elastic-tendon model is used. The type of solution that we look for is one that produces no force or state transients soon after a simulation begins in which activation and path velocity is well approximated as constant. Our preliminary simulations found that satisfactory solutions were found by iterating over both 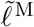 and 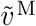 using a nested bisection search that looks for values which approximately satisfies Eqn. 22, result in small values for 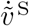 from Eqn. 16, and begins with balanced forces between the two segment titin model in Eqn. 20.

In the outer loop, we iterate over values of 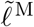. Given *a, ℓ* ^P^, *v* ^P^, and a candidate value of 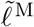, we can immediately solve for *α* and *ℓ* ^T^ using the pennation model. We can numerically solve for the value of another state, *ℓ* ^1^, using the kinematic relationship between *ℓ* ^M^ and *ℓ* ^1^ and by assuming that the two titin segments are in a force equilibrium

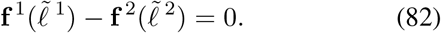

In the inner loop, we iterate over values of 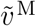 between 0 and *v* ^P^ cos *α* (we ignore solutions in which the sign of *v* ^M^ and *v*^T^ differ) to find the value of 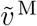 that best satisfies Eqn. 22. Prior to evaluating Eqn. 22, we need to set both 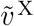 and 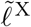. Here we choose a value for 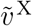 that will ensure that the XE is not producing transient forces

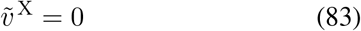

and we use fixed-point iteration to solve for 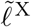 such that Eqn. 16 evaluates to zero. Now the value of 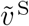 can be directly evaluated using the candidate value of 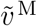, the first derivative of Eqn. 9, and the fact that we have set 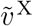 to zero. Finally, the error of this specific combination of 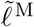 and 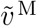 is evaluated using Eqn. 22, where the best solution leads to the lowest absolute value for of 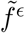 in Eqn. 22. If a rigid-tendon model is being initialized the procedure is simpler because the inner loop iterating over 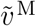 is unnecessary: given *v* ^P^ and 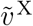 are zero, the velocities 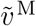 and 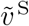 can be directly solved using the first derivative of Eqn. 9. While in principle any root solving method can be used to solve this problem, we have chosen to use the bisection method to avoid local minima.

## D Evaluating a muscle model’s frequency response

To analyze the the frequency response of a muscle to length perturbation we begin by evaluating the length change

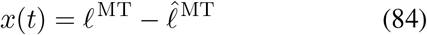

and force change

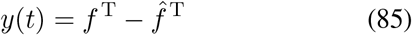

with respect to the nominal length 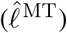 and nominal force 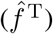. If we approximate the muscle’s response as a linear time invariant transformation *h*(*t*) we can express

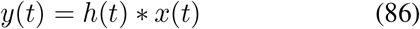

where *∗* is the convolution operator. Each of these signals can be transformed into the frequency-domain [77] by taking the Fourier transform *ℱ* (·) of the time-domain signal, which produces a complex (with real and imaginary parts) signal. Since convolution in the time-domain corresponds to multiplication in the frequency-domain, we have

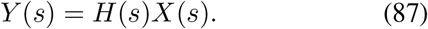

In Eqn. 87 we are interested in solving for *H*(*s*). While it might be tempting to evaluate *H*(*s*) as

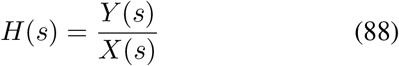

the result will poorly estimate *H*(*s*) because *Y* (*s*) is only approximated by *H*(*s*) *X*(*s*): *Y* (*s*) may contain nonlinearities, non-stationary signals, and noise that cannot be described by *H*(*s*) *X*(*s*).

Using cross-spectral densities, Koopmans [78] (p. 140) derived the estimator

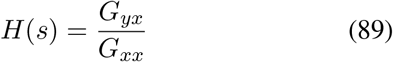

that minimizes the squared errors between *Y* (*s*) and its linear approximation of *H*(*s*) *X*(*s*). The cross-spectral density *G*_*xy*_ between *x*(*t*) and *y*(*t*) is given by

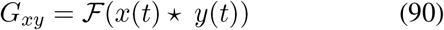

the Fourier transform of the cross-correlation (⋆) between *x*(*t*) and *y*(*t*). When the order of *x*(*t*) and *y*(*t*) are reversed in Eqn. 90 the result is *G*_*yx*_, while *G*_*xx*_ and *G*_*yy*_ are produced by taking the Fourier transform of *x*(*t*) *⋆ x*(*t*) and *y*(*t*) *⋆ y*(*t*) respectively.

Though Koopmans’s [78] estimator is a great improvement over Eqn. 88, the accuracy of the estimate can be further improved using Welch’s method [91]. Welch’s method [91] breaks up the time domain signal into K segments, transforms each segment into the frequency domain, and returns the average across all segments. Using Welch’s method [91] with K segments allows us to evaluate

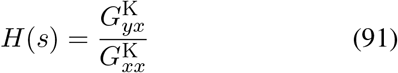

which has a lower frequency resolution than Eqn. 89, but an improved accuracy in *H*(*s*). Now we can evaluate the gain of *H*(*s*) as

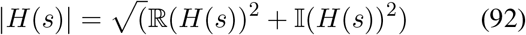

while the phase of *H*(*s*) is given by

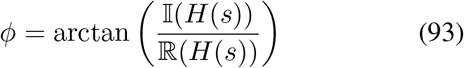

where ℝ (*H*(*s*)) and 𝕀 (*H*(*s*)) are the real and imaginary parts of *H*(*s*) respectively.

The transfer function estimated in Eqn. 91 is meaningful only when *y*(*t*) can be approximated as a linear time-invariant function of *x*(*t*). By evaluating the coherence [79] (p. 137) between *x*(*t*) and *y*(*t*)

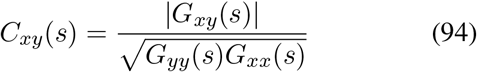

we can determine the strength of the linear association between *X*(*s*) and *Y* (*s*) at each frequency. When *C*_*xy*_ is close to 1 it means that *Y* (*s*) is well approximated by *H*(*s*) *X*(*s*). As *C*_*xy*_ approaches 0, it means that the approximation of *Y* (*s*) by *H*(*s*) *X*(*s*) becomes poor.

Kirsch et al. [5] analyzed a bandwidth that spanned from 4 Hz up to the cutoff frequency of the low-pass filter applied to the input signal *x*(*t*) (15 Hz, 35 Hz, and 90 Hz).

Unfortunately, we cannot use this bandwidth directly when analyzing model output because we have no guarantee that the simulated output is sufficiently linear in this range. Instead, to strike a balance between accuracy and consistency with Kirsch et al. [5], we analyze the bandwidth that is common to Kirsch et al.’s [5] defined range and has the minimum acceptable (*C*_*xy*_)^2^ of 0.67 that is pictured in Fig. 3 of Kirsch et al.

## E Simulation summary data of Kirsch et al

## F Supplementary plots: Gain and phase response rigid-tendon muscle models

## G Supplementary plots: active lengthening on the descending limb

## H Rabbit psoas fibril model parameters

**Figure 17.**
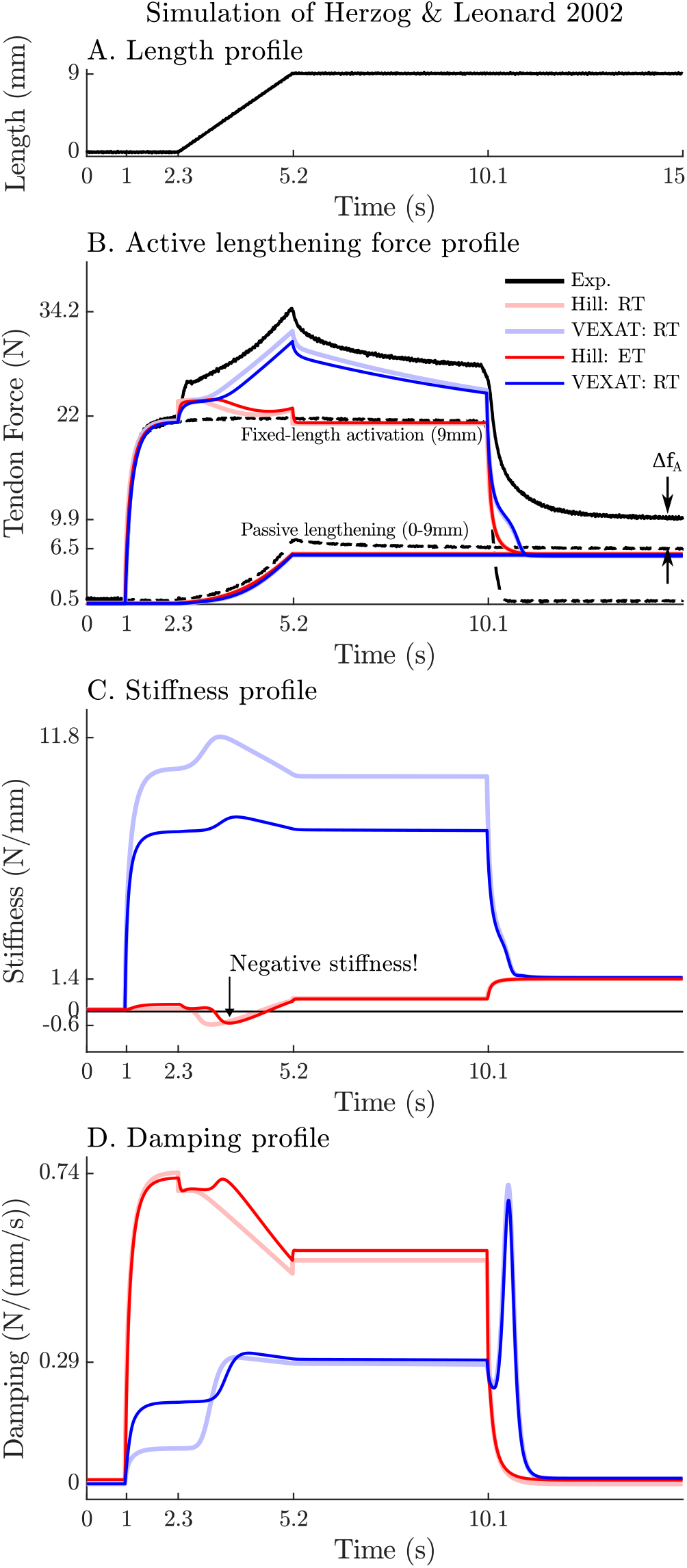
Simulation results of the 3 mm/s (A.) active lengthening experiment of Herzog and Leonard [7] (B.). As with the 9 mm/s trial, the Hill model’s force response drops during the ramp due to a small region of negative stiffness introduced by the descending limb of the force-length curve (C.), and a reduction in damping (D.) due to the flattening of the force-velocity curve. Note: neither model was fitted to this trial.

**Figure 18.**
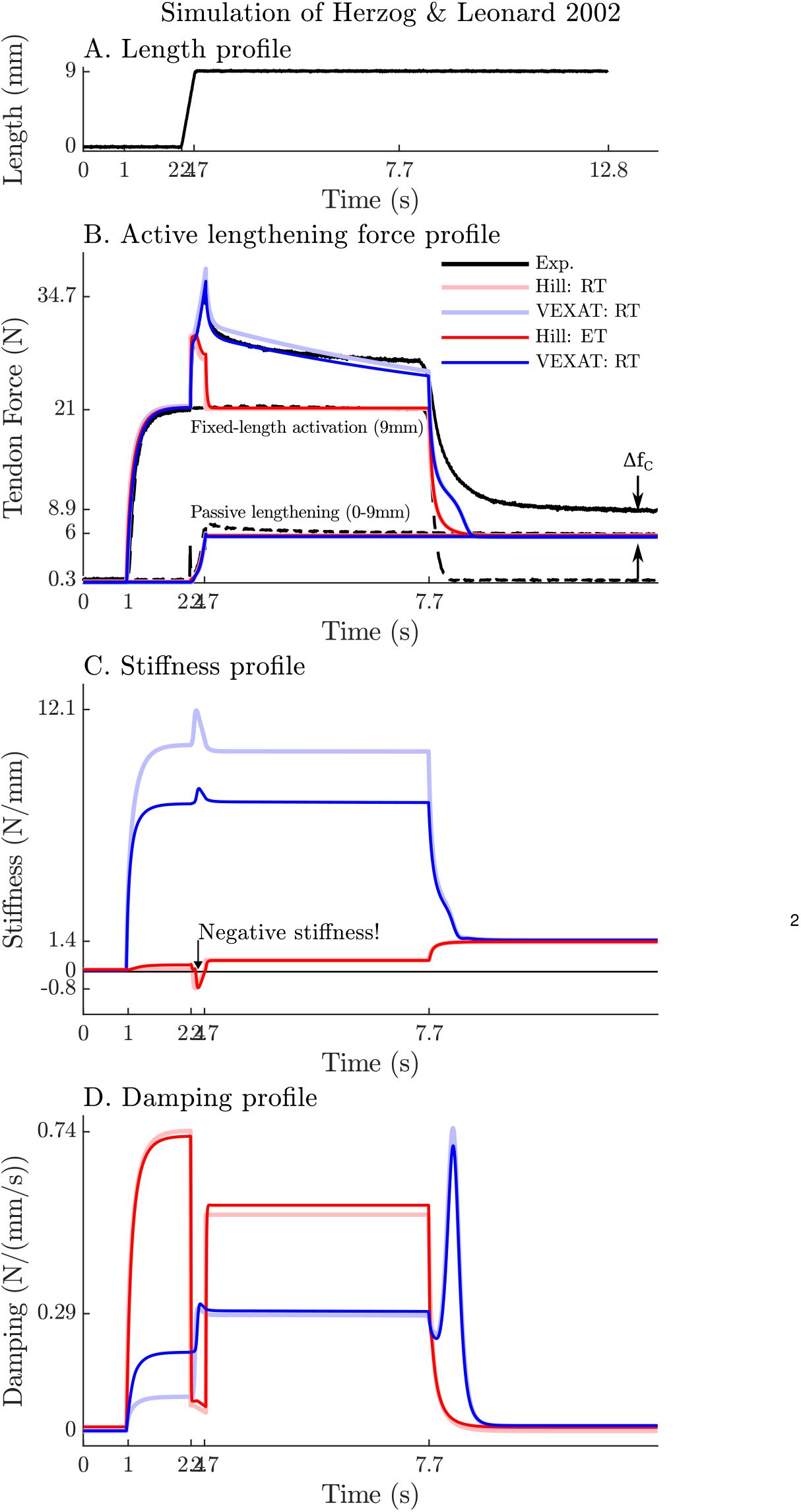
Simulation results of the 27 mm/s (A.) active lengthening experiment of Herzog and Leonard [7] (B.). As with the prior simulations the Hill model exhibits a small region of negative stiffness introduced by the descending limb of the force-length curve (C.) and a drop in damping (D.). Note: neither model was fitted to this trial.

## Footnotes

1 Small in the context of an LTI system is larger than the short-range of Rack and Westbury’s [6] short-range-stiffness: the response of an LTI system can include both length and velocity dependence, while Rack and Westbury’s [6] short-range ends where velocity dependence begins.

2 A Matlab implementation of the model and all simulated experiments are available from https://github.com/mjhmilla/Millard2023VexatMuscle under the branch *elife2023*.

3 The term rheological is used because the model includes a component that deforms with plastic flow in response to an applied force.

4 a change of *±*4 mm to a typical cat soleus with an 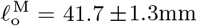 [19]

5 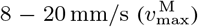 for a muscle with a maximum shortening velocity of 180 mm/s [18]

6 Although activation normally refers to the presence of Ca^2+^ ions in the sarcomere, Ca^2+^ ions alone are insufficient to cause titin to develop enhanced lengthening forces. In addition, crossbridge attachment appears to be necessary: when crossbridge attachment is inhibited titin is not able to develop enhanced forces in the presence of Ca^2+^ during lengthening [8].

7 Which means that the second derivative of the curve is continuous.

8 For readers who require an activation model with continuity to the second-derivative, the model of De Groote et al. [64] is recommended.

9 Note that we have used the symbols D, and not *β*, because the D terms damp the acceleration of actin-myosin movement and as such cannot be interpreted as a viscous damping term. In contrast, viscous damping terms are indicated using the *β* symbol.

10 Physically this assumption is equivalent to treating the CE and the tendon as massless. In general, this assumption is quite reasonable since a cubic centimeter of muscle has a mass of roughly 1.0 g but can generate tensions of between 35-137 N [71]. With such a low mass and a high maximum isometric force, the cubic centimeter of muscle would have to be accelerated at an incredible 3,500-13,700 m*/*s^2^ before the inertial forces would be within 10% of the maximum isometric tension. Since everyday movements require comparatively tiny accelerations, ignoring inertial forces of muscle results in relatively small errors.

11 The impedance (*z*) of two serially connected components (*z*_1_ and *z*_2_) is given by 1*/z* = 1*/z*_1_ + 1*/z*_2_, or *z* = (*z*_1_ *z*_2_)*/*(*z*_1_ + *z*_2_)

12 Kirsch et al. [5] note on page 765 a VAF of 88-99% for the medial gastrocnemius, and 8-10% lower for the soleus.

13 For brevity we will refer to the -3 dB frequency of the perturbation waveform rather than the entire bandwidth

14 See the *elife2023* branch of https://github.com/mjhmilla/Millard2021ImpedanceMuscle

15 See main_ActinMyosinAndTitinStiffness.m in the elife2023 branch of accompanying code repository for details.

16 Figure 8 of Prado et al. [58] shows titin’s contribution ranging from values ranging from (24%-57%) which means that the ECM’s contribution ranges from (43%-76%)

17 Referred to as contour lengths in a worm-like chain model [65]

18 Rabbit psoas titin [58] attaches at the Z-line with a 100nm rigid segment that spans to T12 epitope, is followed by 50 Ig domains, 800 PEVK residues, and another 22 Ig domains until it attaches to the 800 nm half-myosin filament which can also be considered rigid. If the Ig domains were all unfolded (adding around 25 nm [88]) and each PEVK residue could reach a maximum length of between 0.32nm [65] (see Fig. 5: 700nm/2174 residues is 0.32 nm per residue) to 0.38 nm [90] (see pg. 254), two titins in series would reach a length of 2(100nm + 72(25nm) + 800(0.32nm-0.38nm) + 800 nm) = 5192-6008nm. Since rabbit sarcomeres have an 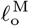 of 2.2*µ*m a sarcomere could be stretched to a length between 5192-6008nm, or 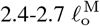, before the contour lengths of the tandem Ig and PEVK segments is reached.

